# SON-dependent nuclear speckle rehabilitation alleviates proteinopathies

**DOI:** 10.1101/2024.04.18.590103

**Authors:** William Dion, Yuren Tao, Maci Chambers, Shanshan Zhao, Riley K. Arbuckle, Michelle Sun, Syeda Kubra, Matthew A. Schaich, Yuhang Nie, Megan Ye, Imran Jamal, Mads B. Larsen, Daniel Camarco, Eleanor Ickes, Haokun H. Wang, C. DuPont, Bingjie Wang, Silvia Liu, Shaohua Pi, Bennet Van Houten, Bill B. Chen, Yuanyuan Chen, Xu Chen, Bokai Zhu

**Affiliations:** Aging Institute of UPMC, University of Pittsburgh School of Medicine, Pittsburgh, PA, U.S.A; Department of Neuroscience, School of Medicine, University of California, San Diego, CA, U.S.A; Department of Ophthalmology, University of Pittsburgh School of Medicine, PA, U.S.A; Department of Human Genetics, University of Pittsburgh Graduate School of Public Health, Pittsburgh, PA, USA; UPMC Hillman Cancer Center, University of Pittsburgh, PA, U.S.A; Department of Pharmacology and Chemical Biology, University of Pittsburgh School of Medicine, PA, U.S.A; Pittsburgh Liver Research Center, University of Pittsburgh, Pittsburgh, PA, U.S.A; Department of Pathology, University of Pittsburgh School of Medicine, Pittsburgh, PA, U.S.A; Division of Pulmonary, Allergy and Critical Care Medicine, Department of Medicine, University of Pittsburgh School of Medicine, Pittsburgh, PA, U.S.A; Division of Endocrinology and Metabolism, Department of Medicine, University of Pittsburgh School of Medicine, Pittsburgh, PA, U.S.A

## Abstract

Current treatments targeting individual protein quality control pathways have limited efficacy in alleviating proteinopathies, highlighting the prerequisite for a common druggable target capable of global proteostasis modulation. Building upon our prior research establishing nuclear speckles as pivotal membrane-less organelles for transcriptional control of proteostasis, we aim to alleviate proteinopathies through nuclear speckle rehabilitation. We identified pyrvinium pamoate as a nuclear speckle rehabilitator that enhances protein quality control gene expression and suppresses YAP1 transcriptional activity via decreasing the surface/interfacial tension of nuclear speckle condensates through interaction with the intrinsically disordered region of nuclear speckle scaffold protein SON. In pre-clinical models, nanomolar pyrvinium pamoate protected against retinal degeneration and tauopathy mainly by promoting autophagy and ubiquitin-proteasome activity in a SON-dependent manner without causing stress. Aberrant nuclear speckle morphology, reduced protein quality control and increased YAP1 activity were observed in human tauopathies. Our study provides proof-of-principle of targeting nuclear speckles to ameliorate proteinopathies.

## Introduction

Proteinopathies often arise from a decline in various proteostasis pathways, including the ubiquitin-proteasome system (UPS), the ER-Golgi protein secretory pathways, and autophagy lysosomal pathway (ALP) ^1, 2^. However, therapies targeting singular pathways have limited efficacy, indicating an incomplete understanding of disease mechanisms. We recently discovered that under physiological conditions, the network of proteostasis pathways manifests as cell-autonomous 12-hour (12h) ultradian rhythms, regulated by a dedicated 12h oscillator, independent from the 24h circadian clock and the cell cycle ^3, 4, 5, 6, 7, 8, 9, 10, 11, 12, 13, 14^. By studying this 12h oscillator, we uncovered an unexpected role of nuclear speckles in global proteostasis control ^9^. Nuclear speckles are membrane-less organelles important for mRNA processing and gene regulation^15, 16, 17, 18, 19^, and their liquid-liquid phase separation (LLPS) dynamics dictate the global transcriptional capacity of proteostasis genes ^9^. Moderate overexpression of the nuclear speckle scaffolding protein SON is sufficient to decrease nuclear speckle sphericity, increase the recruitment of nuclear speckles to chromatin, amplify proteostasis gene expression, and reduce protein aggregation ^9^. Conversely, reducing SON level leads to much more spherical and stagnant speckles, sequesters nuclear speckles away from chromatin, blunts proteostasis gene expression and subsequently elevates intracellular protein aggregates ^20^. More importantly, SON expression decreases with age in various tissues in mice, concomitant with more spherical and smaller nuclear speckles (**Supplementary Fig. 1a-d**). In addition, reduced SON expression was also observed in aging human lungs, and in brain tissues of human subjects with Alzheimer’s disease (**Supplementary Fig. 1e-g**) ^9, 21^. Based upon these results, we herein propose that enhancing SON expression or function, either genetically or pharmacologically, under aging or disease conditions, could potentially restore nuclear speckle morphology and function. This, in turn, would bolster the entire protein quality control system, thereby delaying or even reversing the progression of both aging-related and inherited proteinopathies. We introduce this concept as ‘SON-dependent nuclear speckle rehabilitation/rejuvenation’ (**Fig. 1a**).

**Fig. 1.**
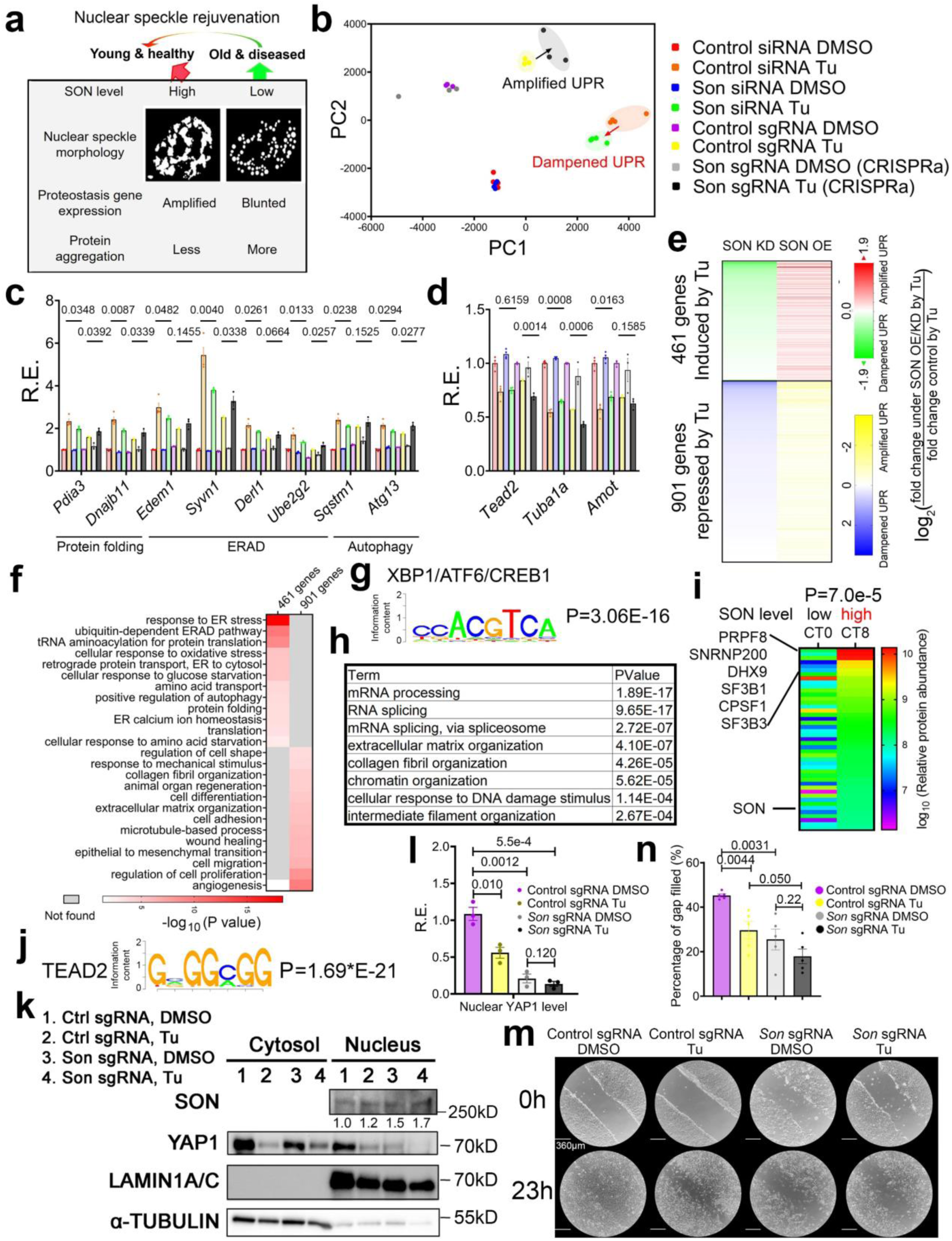
Genetic rehabilitation of nuclear speckle transcriptionally reprograms global proteostasis and YAP1 signaling in an opposing manner. (**a**) The diagram of the approach of nuclear speckle rehabilitation to alleviate proteinopathy. (**b-m**) SON KD, OE and their respective control MEFs were treated with DMSO or 100ng/ml Tu for 6h. (**b**) PCA of global transcriptional response to Tu in the presence of SON OE (via CRISPRa) or KD (via siRNA). (**c, d**) Relative expression (R.E.) of representative proteostasis genes (**c**) and YAP1 target genes (**d**) in response to Tu in the presence of SON OE/KD (n=4 and 3 biologically independent samples for SON KD and SON OE, respectively). (**e**) Heat map of relative fold change of gene expression by Tu in SON OE/KD cells compared to control cells. Only those genes with at least 1.4-fold induction by Tu (log _2_ > 0.5) with a p value smaller than 0.05 in control condition are included. Among these genes are 461 genes whose induction by Tu are further amplified by SON OE (induced more) and dampened (induced less) *by* SON KD; and 901 genes whose repression by Tu are further amplified by SON OE (repressed further) and dampened (repressed less) by SON KD. (**f**) GO analysis of those 461 and 901 genes showing enriched KEGG pathways. (**g**) Enriched XBP1s binding motif ACGTCA on the promoters of 461 genes. (**h**) Top enriched GO terms for top 500 most abundant proteins that are detected in hepatic XBP1s interactome at CT8, as previously reported ^10^. (**i**) Heatmap showing relative abundance of 45 proteins involved in mRNA splicing and processing within the XBP1s interactome at CT0 and CT8, respectively. (**j**) Enriched TEAD2 binding motif GGCGG on the promoters of 901 genes. Western blot (**k**) and quantification (**l**) of nuclear level of YAP1 in control and SON OE cells in response to Tu (n=3 biologically independent samples). Scratch assay with representative images (**m**) and quantification (**n**) of cell migration rate in control and SON OE cells in response to Tu (n=5 biologically independent samples collected from two independent experiments). All data mean ± standard error of the mean (S.E.M.). Statistical tests used: unpaired two-tailed Student’s t-test for c, d, l, and n. Paired two-tailed Student’s t-test for i. All biologically independent samples used for Western blot analyses were collected from at least two independent experiments. The corresponding raw blot images are provided in the source data file.

In this study, we initially characterized the comprehensive transcriptome changes resulting from SON-mediated nuclear speckle rehabilitation and unexpectedly discovered a broader proteostasis framework that incorporates SON-mediated nuclear speckles condensation, unfolded protein response (UPR) transcription factors (TF)-mediated proteostasis gene activation and repression of YAP1 transcriptional activity. Via a high-throughput drug screen, we identified pyrvinium pamoate (PP) as a small molecule nuclear speckle rehabilitator that recapitulates the transcriptome changes elicited by SON overexpression. Mechanistically, via combining cell-free nuclear speckle reconstitution and optical tweeze biophysical experiments, we demonstrated that PP exerts transcriptional reprogramming via reducing the surface/interfacial tension of nuclear speckle condensates and promoting their wetting of genomic DNA via targeting the intrinsically disordered region (IDR) of SON. In preclinical models, PP exhibited strong efficacy in protecting against both tauopathy and retinal degeneration at nanomolar concentration without inducing cellular stress. Lastly, we showed that both the reduction of protein quality control gene expression and increase of YAP1 transcriptional activity is associated with retinal degeneration in mice and tauopathy in humans.

## Result

### Genetic rehabilitation of nuclear speckles transcriptionally reprograms global proteostasis and YAP1 activity in an opposing manner

To determine the extent by which SON transcriptionally reprograms gene expression under both basal and proteotoxic stress conditions, we performed bulk mRNA-Seq on immortalized mouse embryonic fibroblasts (MEFs) with either SON knockdown (KD) by siRNA or overexpression (OE) via CRISPRa^22^, in the absence or presence of the ER stress inducer tunicamycin (Tu) as previously described (**Supplementary Table1**) ^9^. Principal component analysis (PCA) on total mRNA level indicated that while SON manipulation has little effects on global gene expression under basal condition, SON OE and KD significantly amplified and dampened the global transcriptional response to ER stress, respectively (**Fig. 1b**). These include 461 genes that are normally induced, and 901 genes repressed by Tu under normal SON expression condition (**Fig. 1c-e, Supplementary Fig. 2a**). For both groups of genes, we further observed a strong correlation between the relative fold induction or repression for each gene under SON OE and KD conditions (**Supplementary Fig. 2b**), further demonstrating the robustness of bidirectional control on proteostasis gene expression by SON.

As expected, gene ontology (GO) analysis revealed that those ER stress-induced 461 genes are strongly enriched in protein quality control pathways, including protein folding, ER/Golgi quality control, tRNA aminoacylation, ER-associated protein degradation (ERAD), and autophagy (**Fig. 1f, Supplementary Fig. 2a**). To shed light on the mechanisms by which SON transcriptionally amplifies gene activation in response to ER stress, we performed both Landscape In Silico deletion Analysis (LISA) ^23^ and motif analysis to infer the transcriptional regulators that may mediate nuclear speckles interactions with chromatin. Both analyses revealed basic leucine zipper (bZIP) transcription factors (TFs), including ATF6, XBP1, ATF4 and CREB1 as the top candidates (**Fig. 1g and Supplementary Fig. 2c**). ChIP-qPCR analysis revealed enhanced recruitment of nuclear speckles to the 3’ regions of specific proteostasis genes, which are direct targets of XBP1, following SON overexpression under both basal and ER stress conditions (**Supplementary Fig. 2d, e**). To support our in vitro findings, we analyzed a recently published murine in vivo hepatic XBP1s interactome dataset ^10^. We observed a strong enrichment of proteins involved in mRNA splicing and processing within the XBP1s interactome at CT8, a time point when hepatic SON expression peaks (**Fig. 1i**) ^9^. Notably, the three most abundant proteins that interact with XBP1s at CT8—PRPF8, SNRNP200, and DHX9 (**Fig. 1i**)—all have been previously shown to localize in nuclear speckles ^24, 25^. In contrast, at CT0, when SON expression is at the lowest, the interaction between XBP1s and splicing proteins is significantly reduced (**Fig. 1i**). The observed decreased recruitment of splicing proteins to XBP1s at CT0 is not due to reduced XBP1s level itself, as the hepatic XBP1s expression at CT0 is in fact higher compared to CT8 ^3, 26^. These results thus reinforce the notion that nuclear speckles rehabilitation by SON OE is sufficient to amplify the global proteostasis transcriptional activation, likely via facilitating physical interactions between nuclear speckles and UPR TF like XBP1s.

Compared to induced genes, much less is known about the gene programs that are repressed under ER stress. GO analysis revealed that the 901 genes repressed by ER stress are significantly enriched in the Hippo-YAP1 signaling pathway, which regulates a wide range of biological processes, including angiogenesis, axon guidance, epithelial-to-mesenchymal transition (EMT), wound healing, cell adhesion, migration, and extracellular matrix organization. Notable examples of canonical YAP1 target genes include *Tuba1a, Amot* and its transcriptional partner *Tead2* ^27, 28, 29^ (**Fig. 1c-f**). YAP1 and TEAD2 were further predicted to be transcriptional regulators of these 901 ER stress-repressed genes via both LISA and motif analysis (**Fig. 1j and Supplementary Fig. 2c**) and the nuclear YAP1 level was significantly reduced in response to either Tu or SON OE and further decreased upon the combination of the two (**Fig. 1k, l**). We further performed TEAD luciferase reporter assay and found that both SON OE and Tu significantly reduced the TEAD response element-driven luciferase activity in MEFs, with the lowest observed in SON OE cells under ER stress (**Supplementary Fig. 3a**). Scratch assays further confirmed that both SON OE and Tu significantly reduced cell migration in MEFs, with the lowest observed in SON OE cells under ER stress (**Fig. 1m, n**). To rule out the possibility that the global repression of YAP1 transcriptional output during ER stress is specific to MEFs or Tu, we further analyzed a recent transcriptome dataset in the human astrocytoma-derived LN-308 cell line in response to both Tu and thapsigargin (Thap) (another ER stress inducer) treatments ^30^, and observed a strong downregulation of genes involved in YAP1 signaling under both treatments that progressed with time (**Supplementary Fig. 3b-d**). Together, our data indicates that the downregulation of YAP1 transcriptional output is an integral component of the global transcriptional response to proteotoxic stress, and it is also under nuclear speckles control.

Given the established roles of nuclear speckles in mRNA processing ^31, 32, 33^, we next investigated whether SON also regulates mRNA splicing dynamics. While SON manipulation has no effects on the overall transcriptional state of mature mRNA under basal DMSO condition (**Supplementary Fig. 4a bottom**) (consistent with total mRNA shown in **Fig. 1a**), it exerted moderate effects on the pre-mRNA level (**Supplementary Fig. 4a top**), indicating a change in splicing dynamics. By either estimating the relative splicing rates among different groups under basal condition using a simple first-order kinetic model of transcription (see Materials and Methods) or quantifying global intron retention events using the iRead algorithm ^34^, we found that SON increases the splicing rates and improves the splicing fidelity of genes involved in proteostasis and RNA metabolism, and negatively regulates those involved in YAP1-related processes of cell migration, axon guidance, cell adhesion and EMT (**Supplementary Figs. 4b-e, 5a-d and 6a-d**). The significant enrichment in mRNA processing genes themselves under SON control is consistent with known potent autoregulation of splicing factors ^35, 36^. Furthermore, we also found that SON can activate and repress the mature mRNA expression of a select set of proteostasis (albeit very modestly) and cell growth and migration genes, respectively, under basal DMSO conditions (**Supplementary Fig. 7a-c**). Finally, we observed that SON can also activate an anti-viral response gene signature under basal DMSO condition (**Supplementary Fig. 7b**), suggesting a potential broader implication of nuclear speckle rehabilitation in boosting innate immunity. Collectively, our results demonstrated that genetically rejuvenating nuclear speckles reprograms global proteostasis and YAP1 transcriptional output in an opposing manner, under both basal and proteotoxic stress conditions.

Our results thus far demonstrated a tripartite network where nuclear speckle rehabilitation by SON boosts proteostasis and suppresses YAP1, with two possible topologies. In topology one, nuclear speckles can signal both proteostasis and YAP1 signaling directly (**Supplementary Fig. 8a, model 1**), while in topology 2, nuclear speckles repress YAP1 downstream of increased proteostasis gene program (**Supplementary Fig. 8a, model 2**). To distinguish between the two topologies, we examined a recently published RNA-seq dataset in HEK293T cells treated with DMSO, Thap or a very specific XBP1s small molecule activator IXA4 ^37, 38, 39^. While IXA4 can induce a robust proteostasis gene signature similar to that of Thap, it failed to repress YAP1 transcriptional output genes as Thap did (**Supplementary Fig. 8b-d**). Collectively these results support the first topology where nuclear speckles can program proteostasis gene expression and YAP1 transcriptional output in parallel, likely via promoting physical interaction between nuclear speckles and XBP1s for the former and triggering YAP1 nuclear exclusion for the latter (**Supplementary Fig. 8e**). We speculate the opposing changes in proteostasis and YAP1 signaling may reflect an energetic trade-off between proteostasis and the control of cell dynamics under proteotoxic stress (**Supplementary Fig. 8f**).

### High-throughput screen (HTS) identified pyrvinium pamoate (PP) as a SON-dependent nuclear speckle rehabilitator

Having established the proof-of-principle of nuclear rehabilitation via SON OE, we next explore the feasibility of rejuvenating nuclear speckles pharmacologically. Since SON OE and KD reduced and increased the sphericity of nuclear speckles ^20^, respectively, putative nuclear speckle rehabilitators are expected to reduce the sphericity of speckles. We started with a library of over 2,500 U.S. Food and Drug Administration (FDA)-approved drugs and ran a primary HTS to identify compounds that could reduce nuclear speckles sphericity, followed by a secondary screen to identify those further capable of amplifying *Perk-*promoter driven dGFP expression (*Perk* is a UPR target) in a dose-dependent manner (**Fig. 2a**). To narrow down the five final candidates - the tyrosine kinase inhibitors nintedanib (NB) and ponatinib (PB), the anti-microbial proflavine hemisulfate (PH) and proflavine, and the anthelmintic pyrvinium pamoate (PP) (**Supplementary Fig. 9a-e**), RNA-seq analysis was performed to compare the transcriptomes of these compounds to those of SON OE and KD. In the end, PP was identified as the most likely nuclear speckle rehabilitator (**Fig. 2a**). A strong dose-response of PP in reducing the nuclear speckle sphericity was observed up to 0.3µM (**Fig. 2b and Supplementary Fig. 9c**). PP also increased the perimeter of nuclear speckles (**Fig. 2c**), indicating that PP can increase the surface area of nuclear speckles in the three-dimensional space of the nucleus.

**Fig. 2.**
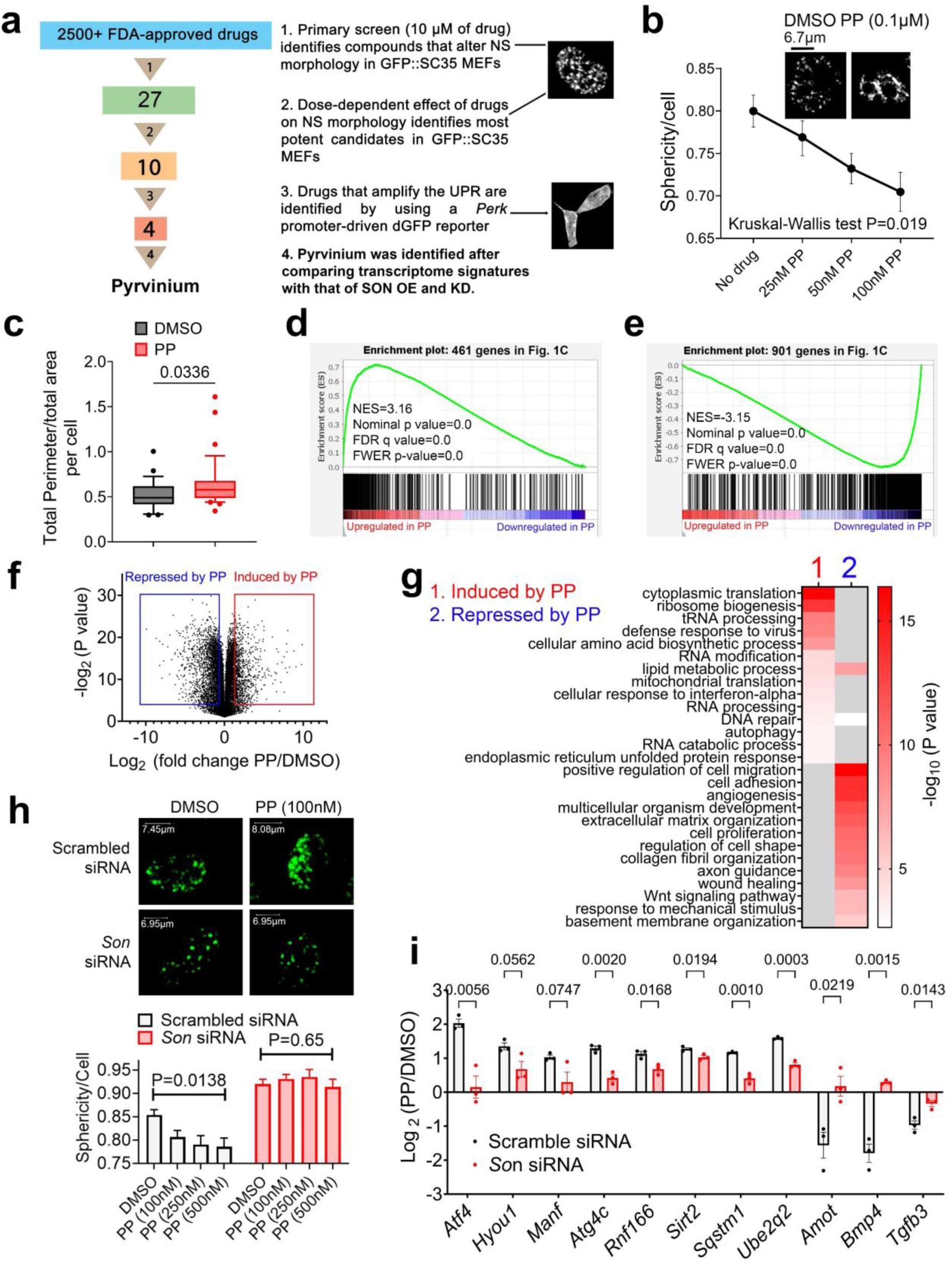
HTS identifies PP as a SON-dependent nuclear speckle rehabilitator. (**a**) Workflow detailing our initial drug screen and subsequent steps to identify nuclear speckle rehabilitators. (**b**) Dose-dependent effect on nuclear speckles morphology by PP, with a representative image of nuclear speckles under DMSO or 0.1µM of PP (n=25∼57 cells collected evenly across three separate wells for each treatment). (**c**) Quantification of total area-normalized perimeter of nuclear speckles in control and 1µM PP per cell (n=26 and 35 cells for DMSO and PP, respectively, collected from three independent experiments). (**d**) GSEA showing a similar transcriptome signature between PP-upregulated genes and 461 genes further amplified by SON OE during ER stress. (**e**) GSEA analysis showing a similar transcriptome signature between PP-downregulated genes and 901 genes further repressed by SON OE during ER stress. (**f**) Volcano plot showing fold change by PP versus log transformed p values. Genes induced or repressed by at least 1.41-fold with a p value smaller than 0.05 are boxed. (**g**) GO analysis of differentially expressed genes by PP. (**h**) Representative images and quantification of sphericity of GFP signal from GFP::SRSF2 MEFs with scrambled or *Son* siRNA treated with DMSO or increasing concentration of PP for 25 hours (n=51 and 31 cells for scrambled and *Son* siRNA, respectively, collected from two independent experiments). (**i**) Log_2_ normalized fold change in response to PP treatment (0.3 µm) for 24 hours in control and SON KD MEFs (n=3 biologically independent samples). All data mean ± S.E.M. Statistical tests used: unpaired two-tailed Student’s t-test for c and i. Kruskal-Wallis test for b, and Ordinary one-way ANOVA for h.

PP triggers a transcriptional response with strong resemblance to SON OE cells in response to ER stress (**Fig. 2d, e and Supplementary Fig. 10a-d**), including both upregulated genes implicated in protein quality control and downregulated genes involved in the regulation of cell dynamics (**Fig. 2f, g and Supplementary Fig. 10a-d**). LISA analysis on differentially expressed genes by PP revealed bZIP TFs ATF4 and YAP1 among top transcriptional regulators of upregulated and downregulated genes, respectively (**Supplementary Fig. 10e**). 1µm PP elevated ATF4 expression and, to a lesser extent, XBP1s (**Supplementary Fig. 10f, g),** while also inducing a modest reduction of nuclear YAP1 level (**Supplementary Fig. 10h)**, without altering SON levels (**Supplementary Fig. 10i**).

We performed additional comparative transcriptome analysis to further validate PP as a SON-dependent nuclear speckle rehabilitator. First, when comparing the fold induction or repression of gene expression by PP and Tu, the signature of PP is more similar to that of Tu under SON OE compared to under SON KD condition (p= 0.00195 by Chow tests) (**Supplementary Fig. 11a**). Secondly, similar to SON OE (**Supplementary Fig. 7b**), PP also induced expression of genes involved in anti-viral response (**Supplementary Fig. 11b-f**). Thirdly, GSEA indicated a strong resemblance of gene signatures repressed by PP and SON OE that are enriched in the control of cell dynamics, under basal conditions in the absence of ER stress (**Supplementary Fig. 12a-e**). Lastly, like SON (**Supplementary Figs. 5d and 6d**), PP also improves the splicing fidelity of splicing genes themselves (**Supplementary Fig 13a-f**), again reflecting the autoregulation of splicing factors. To experimentally confirm that PP rejuvenates nuclear speckles in a SON-dependent manner, we knocked down *Son* via siRNA in MEFs. Son knockdown leads to smaller and more spherical speckles, consistent with our previous study ^20^ (**Fig. 2h**). Importantly, PP’s ability to reduce nuclear speckle sphericity is abolished in *Son* knockdown MEFs (**Fig. 2h**). Subsequently, *Son* knockdown impaired PP’s ability to both activate protein quality control gene expression and repress YAP1 transcriptional output (**Fig. 2i and Supplementary Fig. 13g**). Taken together, these results indicate that PP is a *bona fide* SON-dependent nuclear speckle rehabilitator.

### PP reduces the surface tension of SON IDR condensates

To determine whether PP can physically interact with SON in MEFs, we performed cellular thermal shift assay (CETSA), which is based on ligand-induced thermal stabilization of target proteins, whereas unbound proteins denature, aggregate and precipitate at elevated temperatures, ligand-bound proteins remain soluble due to increased stability ^40, 41^. Using a SON-specific antibody (**Supplementary Fig. 14a**), we found that PP induced a thermal shift of SON with a direction consistent with stabilization (**Fig. 3a**). As negative controls, we found that PP does not stabilize SRSF2 (SC35), another nuclear speckle protein, or the ISR/UPR TF ATF4, whose expression is nonetheless significantly increased by PP (**Fig. 3b and Supplementary Fig. 14b**).

**Fig. 3.**
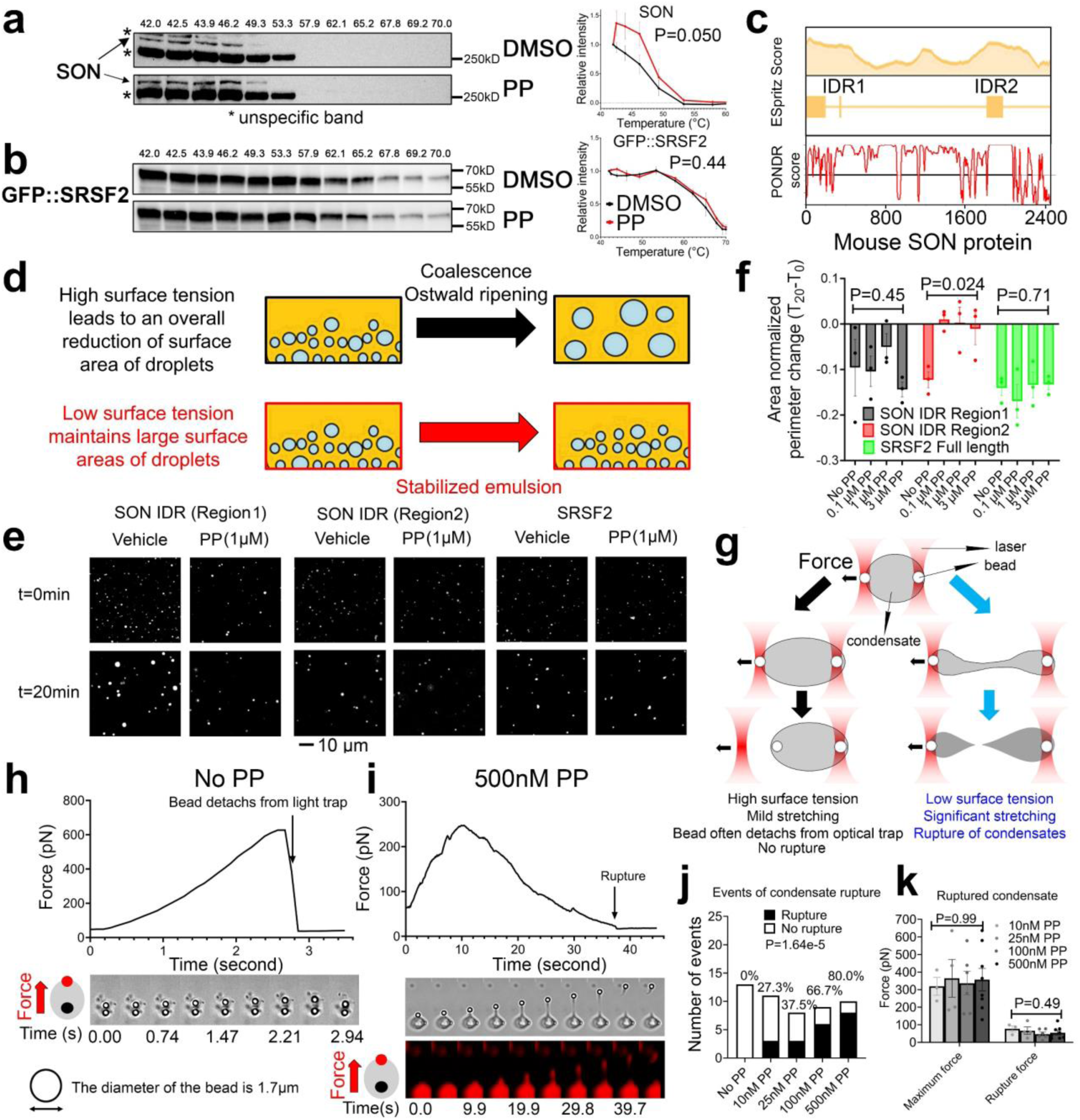
PP reduces the surface tension of SON IDR2 condensate. (**a, b**) CETSA of SON (**a**) and GFP::SRSF2 (**b**) with 3µM PP. Both representative blot and quantification are shown (SON: n=6 and 5 quantifications collected from six and five independent CETSAs for DMSO and PP, respectively; GFP::SRSF2: n=3 quantifications collected from three independent CETSAs for both DMSO and PP). (**c**) Computational prediction of IDR in mouse SON. (**d**) Diagram illustrating how surface tension influences droplets coalescence kinetics. (**e, f**) Representative images of droplet formation assay with different recombinant proteins (**e**) and quantification (**f**) of area-normalized perimeter changes in the time span of 20 minutes with 125mM NaCl (n=3 quantifications collected from three independent droplet formation assays for each treatment). (**g**) Diagram of the optical tweezer set up. Image created, in part, with BioRender.com (**h**) Force recording as well as time lapse bright field images of condensate stretching without PP. (**i**) Force recording as well as time lapse fluorescent and bright field images of condensate stretching with 500nM PP. (**j, k**) Quantification of the percentage (**j**) and the maximum and rupture force (**k**) (n=3∼8 ruptured events quantified from three independent C-Trap experiments for No PP and 500nM PP, and two independent C-Trap experiments for 10, 25 and 100nM PP) of ruptured SON IDR2 condensates with increasing concentration of PP. All data mean ± S.E.M. Statistical tests used: mixed-effects analysis for a and b. Ordinary one-way ANOVA for f. Chi-square test for trend for j. Kruskal-Wallis test for k. Western blot raw blot images are provided in the source data file.

SON is the central scaffold protein of nuclear speckles and essential for their formation ^42, 43, 44^,. By contrast, SRSF2 (SC35) is one of the subunits of the spliceosomes and has a broader spatial distribution also occupying the periphery of nuclear speckles, at the interface between nuclear speckles and chromatin and is dispensable for speckle formation ^45, 46^. PP generated a less spherical nuclear speckle with larger surface area (**Fig. 2b, c**), suggesting that PP could reduce the surface tension of speckles (surface tension is the tendency of liquid droplets to minimize the total surface area, therefore an increased surface area is suggestive of reduced surface tension^47^). Given the CETSA data indicating PP can bind to SON directly, we next tested whether PP can directly impact the condensates formation of two nuclear speckle proteins, SRSF2 and SON, using an *in vitro* droplet formation assay ^48^. Using different computational algorithms to search for intrinsically disordered region (IDR) ^49, 50, 51^, we identified two IDRs at the N and C terminals of mouse SON (**Fig. 3c**). While both SON IDRs are low complexity mixed charge domains (MCD) containing both negative (D/E) and positive charged (K/R) amino acids ^52^, the C-terminal IDR of SON (SON IDR2) is significantly more enriched in arginine than the N-terminal SON IDR1 (**Supplementary Fig. 14c, d**). We separately cloned the regions encoding both SON IDRs and the full-length SRSF2 into the C-terminal of mCherry and purified recombinant proteins from *E. coli* (**Supplementary Fig. 15a, b**). Purified recombinant proteins were added to buffers containing 10% crowding reagents PEG-8000 to form droplets, as previously described ^48^. Confocal fluorescence microscopy revealed mCherry positive, micron-sized spherical droplets freely moving in solution and wetting the surface of the glass coverslip (**Supplementary Movies 1, 2**). All droplets were highly spherical, exhibited fusion/coalescence behaviors (**Supplementary Movies 1, 2**), and scaled in size and number positively with increasing concentration of proteins and negatively with increasing salt concentration (**Supplementary Fig. 15c, d**), all properties expected for liquid-like droplets ^48^.

Due to surface tension, small droplets will eventually morph into a fewer number of large droplets, resulting in a net decrease of surface area, either via coalescence or Ostwald ripening^53^ (**Fig. 3d**), which was seen for all protein droplets after 20 minutes of time lapse imaging (**Fig. 3e, f, and Supplementary Fig. 15e**). Addition of nanomolar concentration of PP to SON IDR2, but not SON IDR1 and SRSF2 condensates, significantly reduced the kinetics of this process (**Fig. 3d-f, and Supplementary Fig. 15e, f**). The significance of SON IDR2 is reinforced by the substantial evolutionary conservation of its sequences from flies to humans (**Supplementary Fig. 15g**).

To verify that PP could indeed reduce the surface tension of SON IDR2 condensates, we further used an optical tweezer setup to optically manipulate two polystyrene beads attached to opposite poles of each SON IDR2 condensate. By applying force outward to one bead while keeping the other fixed in place, we could directly observe condensate deformation in real-time and measure the force needed to stretch and rupture them (**Fig. 3g**), the latter of which linearly correlates with the condensate’s surface/interfacial tension force ^54^. In the absence of PP, no rupturing events were observed. Instead, the beads frequently detached from the optical trap once the condensate’s surface tension exceeded the maximum optical force limit (**Fig. 3h, Supplementary Fig. 16a, and Supplementary Movies 3-5**). However, as PP concentration increased, rupturing events became more frequent, reaching an 80% rupture rate with 500 nM PP (**Fig. 3i, j, Supplementary Fig. 16a, and Supplementary Movies 6-13**). Notably, the average force needed to rupture each condensate (∼60pN) remained unchanged even as PP concentration increased (**Fig. 3k**). These findings—an increase in rupture probability without a decrease in rupture force—argues against the site-specific interaction between each PP and each SON IDR2 molecule. Instead, it suggests that each SON IDR2 condensate interacts with PP as a collective unit: the free energy of PP partitioning into SON IDR2 condensate emerges only upon the phase separation of the latter (**Supplementary Fig. 16b**) ^55^. In sum, these results strongly indicate that PP can reduce the surface tension of SON IDR2 condensates in a cell-free system.

### PP promotes the wetting of DNA by nuclear speckle condensates *in vitro*

To better recapitulate the heterogeneous compositions of nuclear speckles in the cell-free system, we further supplemented recombinant mCherry-SON IDR2 with nuclear extract (NE) from HeLa cells that include all active components of transcription and splicing factors (**Fig. 4a**) ^48^. Mass spectrometry confirmed that HeLa NE-supplemented mCherry-SON IDR2 condensates preferentially compartmentalized splicing factors (including SRSF2), with twelve of the top fifteen enriched proteins previously identified in nuclear speckles ^24, 25^ (**Supplementary Fig. 17a, b**). By contrast, other nuclear proteins like proteasome subunits, DNA repair factors or general transcription factors were not enriched in the reconstituted condensates (**Supplementary Fig. 17a, b**). These condensates further exhibited less spherical morphology, had increased number and total size (**Supplementary Fig. 17c, d**), features expected from nuclear speckle-like condensates with viscoelastic properties ^56^. Importantly, the addition of PP reduced both the sphericity and surface tension of HeLa NE-supplemented SON IDR2 condensates in a dose-dependent manner (**Fig. 4b, c**).

**Fig. 4.**
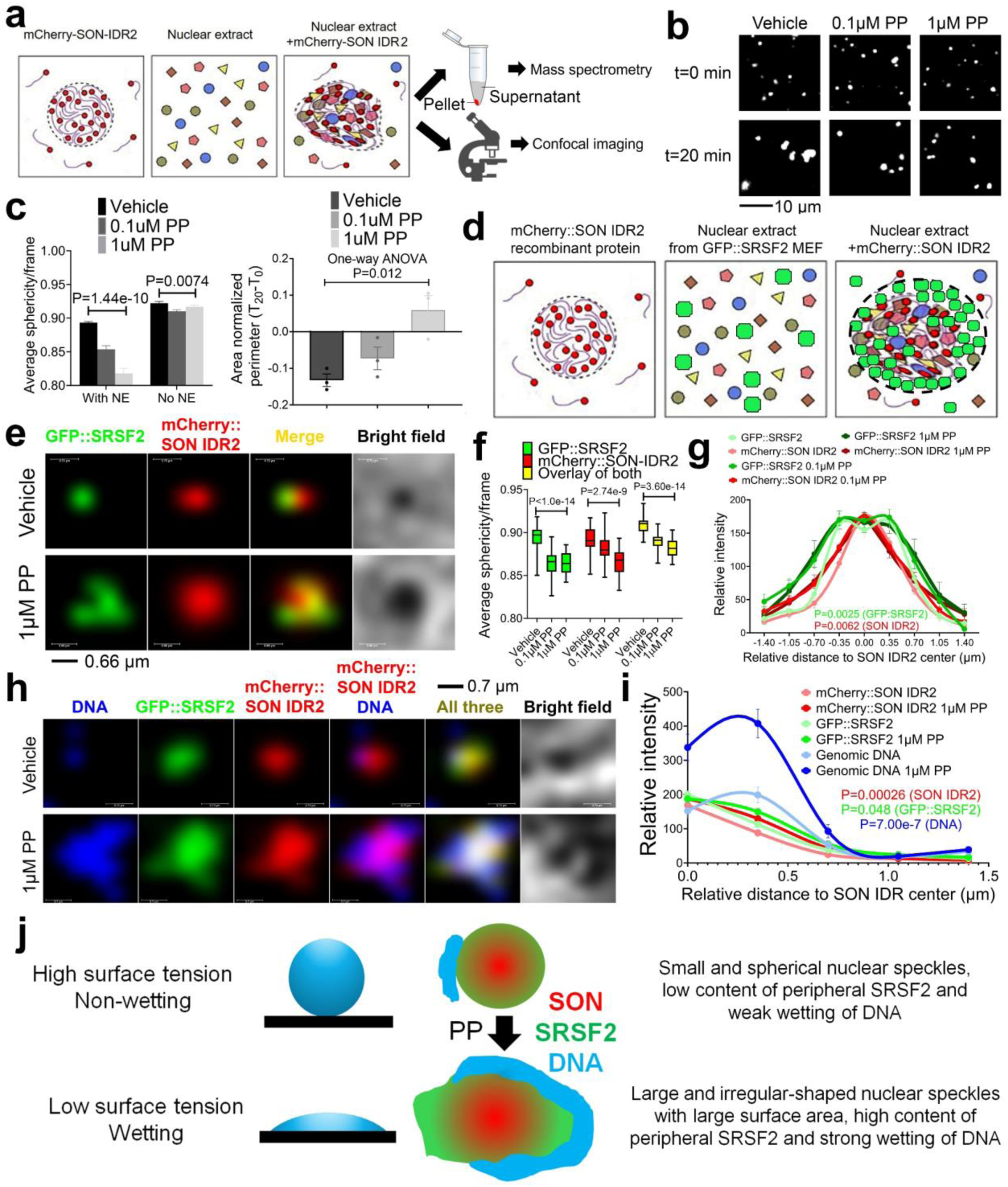
PP promotes the wetting of genomic DNA by nuclear speckle condensates. (**a**) Diagram showing NE-supplemented SON IDR2 condensates are expected to compartmentalize splicing factors and exhibit less spherical morphology. (**b, c**) Representative images of droplet formation with HeLa NE-supplemented SON IDR2 with increasing concentration of PP (**b**) and quantification of sphericity (n=12 quantifications from 12 image frames for NE and 42 quantifications from 42 image frames for without NE, collected evenly across three independent droplet formation assays for each treatment) (**c left**) and area-normalized perimeter changes (n=3 quantifications collected from three independent droplet formation assays for each treatment) (**c right**) in the time span of 20 minutes. (**d-g**) Diagram of droplet formation assay where SON IDR2 is expected to compartmentalize splicing factors, including GFP::SRSF2 into the nuclear speckle-like condensates. GFP::SRSF2 is expected to exhibit a broader spatial distribution than the SON IDR2 core (**d**). Representative images (**e**) and quantification of sphericity (n=30 quantifications from 30 image frames per channel, collected evenly across three independent droplet formation assays for each treatment) (**f**) and spatial distribution of mCherry::SON IDR2 and GFP::SRSF2 (n=15∼41 nuclear speckles collected evenly across three independent droplet formation assays) (**g**). (**h, i**) Mouse genomic DNA was further added to the solution. Representative images (**h**) and quantification (n=60∼61 nuclear speckles collected evenly across three independent droplet formation assays) (**i**) of spatial distribution of mCherry::SON IDR2, GFP::SRSF2 and DNA. (**j**) Diagram showing how PP reduces the surface tension of nuclear speckle condensate to promote wetting of genomic DNA. All data mean ± S.E.M. Statistical tests used: mixed-effects analysis for g. Ordinary one-way ANOVA for c and f. Two-way RM ANOVA for i.

To investigate whether PP may directly affect the relative spatial distribution of SON and SRSF2 within nuclear speckles, we further supplemented recombinant mCherry-SON IDR2 with NE from GFP::SRSF2-expressing MEFs (**Fig. 4d**). The resulting nuclear speckle-like condensates recapitulated the anticipated spatial distribution of SON and SRSF2 proteins, with the former located at the center, and the latter exhibiting a broader distribution with its highest concentration often observed at 350nm away from the SON IDR2 center (**Supplementary Fig. 17e and Supplementary Movie 14**). While PP does not alter the relative spatial distribution of SON IDR2 and SRSF2, it reduced their sphericity, and markedly increased SRSF2 content at the periphery of nuclear speckle-like condensates (**Fig. 4e-g**). To determine whether PP may also influence the wetting of genomic DNA by nuclear speckles ^57^, we further added mouse genomic DNA into the droplet solution. In accordance with observations in intact cells, nuclear speckle condensates largely don’t mix with but can wet the DNA (**Supplementary Fig. 17f**). Interestingly, the addition of PP more than doubled the wetting of genomic DNA by the reconstituted nuclear speckles (**Fig. 4h, i**).

To validate that PP can alter the nuclear speckle LLPS properties *in vitro* in the context of intact cells, we further performed 1,6 hexanediol sensitivity assay ^58^ using the same EGFP::SRSF2 MEFs ^9^. A short term 1,6 hexanediol treatment preferentially dissolves liquid but not solid condensates, thus a change in the sensitivity to 1,6 hexanediol reflects an alteration in the viscoelastic property of a given condensates ^58^. As demonstrated in **Supplementary** Fig. 18a, b, PP desensitized SRSF2 to the increasing concentrations of 1,6 hexanediol. This effect was not observed on two other biomolecular condensates, the nuclear MED1 ^48^ and cytosolic GW182 present in P-bodies ^59^ (**Supplementary Fig. 18c, d**). Taken together, these results suggest a mechanism where PP can reduce the surface/interfacial tension of nuclear speckles via targeting SON IDR2, leading to larger surface areas with increased SRSF2 content at the periphery, increased wetting of genomic DNA and subsequently a higher portion of spliceosomes stably engaging in active RNA processing and transcription elongation of proteostasis genes (**Fig. 4j**). Since no active transcription occurs in the *in vitro* droplet formation assay, these results demonstrate that PP-mediated nuclear speckle LLPS change is a cause, rather than a consequence of or response to, global transcriptional reprogramming.

### PP reduces cellular pathological Tau and rhodopsin levels by boosting ALP and UPS, at the expense of YAP1 signaling

Since PP leads to a global increase of protein quality control gene expression, we went on to determine the effects of PP on global protein synthesis, and degradation via UPS and ALP with two different concentrations: 0.1 and 1µM. By performing puromycin incorporation assay and examining markers of ISR ^60^, we found that 0.1 µM of PP did not alter the global protein synthesis (**Fig. 5a, b, and Supplementary Fig. 19a, b**). To quantify the UPS activity, we treated MEFs with PP alone or in combination with the proteasome inhibitor MG132 and blotted for high molecular weight poly-ubiquitinated proteins. 0.1 µM PP led to a significant reduction of poly-ubiquitinated protein levels (**Fig. 5c, d**), which is completely restored by MG132 co-treatment, indicating increased UPS flux (**Fig. 5c, d**). We further directly measured the activity of proteasome complex and found 0.1 µM PP significantly increased the proteasome activity in a SON-dependent manner (**Supplementary Fig. 19c**). To quantify the autophagic flux, we utilized a tandem LC3 reporter mCherry-GFP-LC3 where an increase in the number of red-fluorescent cytosolic puncta indicates increased autolysosome formation and autophagic flux (**Fig. 5e**) ^61^, and found that 0.1 µM PP markedly increased the formation of autolysosomes (**Fig. 5f**). To verify this result, we further blocked autophagy at the late stage autophagosome-lysosome fusion step using Bafilomycin A1 (BafA) ^62^, and quantified the level of LC3I and LC3II with or without PP. 0.1 µM PP resulted in a reduced level of LC3II, which was completely restored by BafA co-treatment, again indicating increased autophagic flux (**Supplementary Fig. 19d,e**). Unlike 0.1µM PP, 1µM PP leads to a global translation repression (**Supplementary Fig. 20a, b**) concomitant with ATF4 induction (**Supplementary Fig. 10g**), but is still able to promote both UPS and ALP (**Supplementary Fig. 20c-f)**. In sum, these findings demonstrate that nanomolar concentrations of PP effectively rejuvenate nuclear speckles, enhancing both UPS and ALP without inducing cellular stress. However, at higher micromolar concentrations, PP also induces cellular stress, likely due to its known inhibitory effects on mitochondrial activity^63, 64^.

**Fig. 5.**
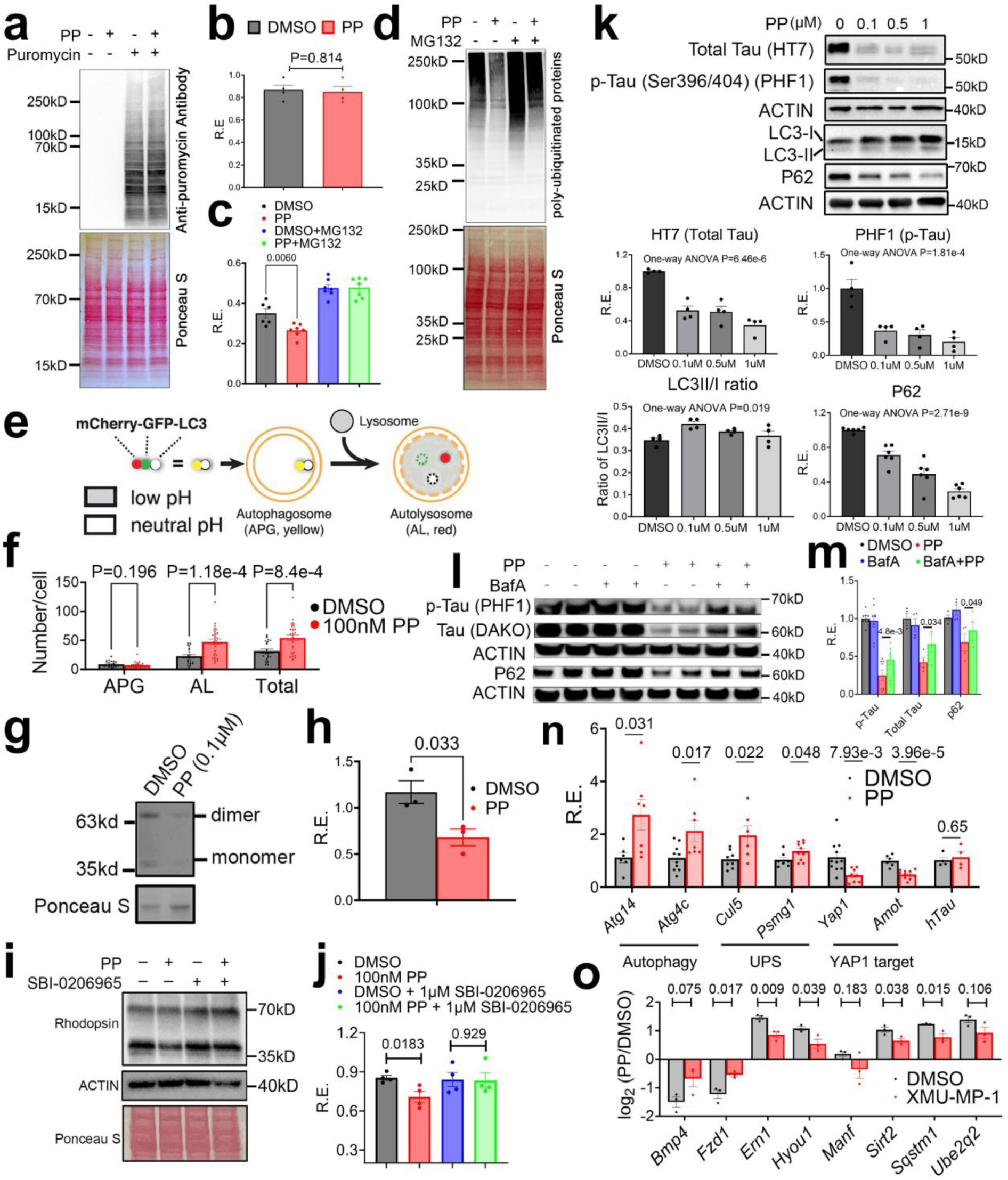
PP reduces pathological Tau and Rhodopsin level by boosting autophagy and UPS activity. (**a-d**) MEFs were treated with DMSO or 0.1µM PP for ∼24 hours and then co-treated with or without puromycin (10 μg/mL for 30 minutes) (n=4 biologically independent samples), or MG132 (10μM for 110 minutes) (n=6∼7 biologically independent samples). Western blot and quantification of puromycin-incorporated proteins (**a, b**), and poly-ubiquitinated protein (**c, d**). (**e, f**) MEFs were treated with DMSO or 0.1µM PP for ∼24 hours and autophagic flux was measured via a mCherry::GFP::LC3 fusion protein reporter. Chimeric proteins comprising LC3B fused with both GFP and mCherry offer a method to track autophagic flux. Autophagosomes (APG) marked by mCherry::GFP::LC3 exhibit both mCherry and GFP signals. Following fusion with lysosome to form autolysosome (AL), GFP signals diminish significantly in the acidic environment, while mCherry signals remain relatively stable. (**e**). Quantification of the number of APG, AL and total vesicles (**f**). n=23 and 29 cells for DMSO and PP, respectively, collected evenly across four separate wells for each treatment from two independent experiments. (**g, h**) NIH3T3 RHO^P23H^ cells were treated with 0.1µM PP for 24 hours and Western blot (**g**) and quantification (**h**) of RHO^P23H^ level (n=3 biologically independent samples). (**i, j**) 0.1µM PP-treated NIH3T3 RHO^P23H^ cells were co-treated with or without 1μM SBI-0206965 for 24 hours. Western blot (**i**) and quantification (**j**) of RHO^P23H^ level (n=4 biologically independent samples). (**k-m**) Tau (P301S)-expressing primary neurons were treated with increasing concentration of PP for 24 hours, and western blot and quantification of different proteins (n=4 biologically independent samples for HT7, PHF1 and LC3, and n=6 biologically independent samples for P62) (**k**) or treated with 0.1µM PP for 12 hours in the presence or absence of BafA (50nM) in the last hour and western blot (**l**) and quantification (n=4 biologically independent samples for total Tau and P62, and n=8 biologically independent samples for p-Tau) (**m**) of different proteins. (**n**) Tau P301S-expressing neurons were treated with DMSO or 0.1µM PP for 12 hours and qPCR of selective proteostasis and YAP1 target genes (n=6∼11 biologically independent samples) and hTau (n=4 biologically independent samples). (**o**) MEFs were treated with DMSO, PP (1µM), XMU-MP-1 (1µM) or XMU-MP-1+PP for 24 hours, and qPCR of protein quality control and YAP1s output gene expression quantified as log _2_ transformed fold change under DMSO or XMU-MP-1 condition by PP (n=3 biologically independent samples). All data mean ± S.E.M. Statistical tests used: unpaired two-tailed Student’s t-test for b, c, f, h, m, n and o. Ordinary one-way ANOVA for k. Paired (biological repeats were run separately) two-tailed Student’s t-test for m. All biologically independent samples used for Western blot analyses were collected from two to three independent experiments. The corresponding raw blot images are provided in the source data file.

To determine whether PP can protect against proteinopathies via boosting UPS and ALP, we focused on two different diseases, a genetic form of Retinitis Pigmentosa (RP) with a proline to histidine (P23H) mutation in Rhodopsin (RHO) protein, and tauopathy. Recently, several studies suggested that boosting protein quality control can protect against mouse models of RP by increasing elimination of the mutant RHO^P23H^ protein ^65, 66, 67, 68^. To determine if PP can also reduce RHO^P23H^ level, we used a NIH3T3 cell line ectopically expressing RHO^P23H^ protein ^69^. Since nanomolar concentration of PP is sufficient to boost both UPS and ALP without triggering cellular stress, we selected 0.1 µM as the working concentration. Treating this cell line with 0.1 µM of PP for 24 hours led to a reduction of RHO^P23H^ protein in a SON-dependent manner (**Fig. 5g, h, and Supplementary Fig. 21a, b**). Blocking autophagy with ULK1/2 inhibitor SBI-0206965 or BafA, and UPS with MG132, respectively, abolished the effect of PP on reducing RHO^P23H^ protein level (**Fig. 5i, j, and Supplementary Fig. 21c-f**), suggesting that both increased ALP and UPS are responsible for the increased elimination of RHO^P23H^ by PP.

Both UPS and ALP are also involved in the degradation of tau protein in tauopathies ^70, 71, 72, 73^. To test the effect of PP on tau proteostasis in mouse primary neuronal cultures, we overexpressed human Tau carrying P301S mutation – a frontal temporal dementia (FTD)-causing mutation in the human *MAPT* gene (Tau) ^74^. Nanomolar PP significantly reduced both total and p-Tau (Ser396/404) protein in a dose-dependent manner (**Fig. 5k**), without inducing any observable signs of neuronal toxicity (**Supplementary Fig. 21g**). 100nM PP also promoted autophagic flux in neurons (**Fig. 5k and Supplementary Fig. 21h, i**) ^75^. Blocking autophagic flux with BafA dampened the effects of PP on reducing p-Tau and total Tau (**Fig. 5l, m**). As observed in fibroblasts, 100nM PP also promoted UPS activity in neurons; however, inhibiting proteasome activity with MG132 has minimal effects on PP’s ability to reduce Tau level (**Supplementary Fig. 21j, k**). As a negative control, we found that PP does not reduce the transcription of hTau gene (**Fig. 5n**). Consistent with PP’s ability to boost UPS and ALP, 100nM PP increased the expression of genes involved in both pathways in P301S hTau-expressing neurons (**Fig. 5n**). Collectively, these results indicate that increased ALP partially underlies PP’s effect in reducing Tau level in mouse primary neurons.

Nuclear speckle rehabilitation by SON OE or PP increases global protein quality control at the cost of reduced YAP1 signaling in both MEFs (**Fig. 2i**) and neurons (**Fig. 5n**). To address whether YAP1 downregulation also contributed to PP’s efficacy in alleviating proteinopathy, we restored YAP1 signaling with previously published YAP1 activators XMU-MP-1 and/or TRULI ^76, 77,78^. We found that while XMU-MP-1 antagonized the downregulation of YAP1 target genes by PP as expected, it also potently dampened the upregulation of proteostasis genes (**Fig. 5o**). In addition, both XMU-MP-1 and TRULI negated PP’s effect on reducing RHO^P23H^ level in NIH3T3 cells (**Supplementary Fig. 22a-d**). In addition, knocking down MST1, a kinase that inhibits YAP1 activity ^79^, similarly blocked the ability of PP to reduce RHO^P23H^ level (**Supplementary Fig. 22e, f**). Similarly, TRULI also blocked PP’s effect on reducing Tau in primary neurons (**Supplementary Fig. 22g, h**). These results suggest that maintaining reduced YAP1 activity is also essential for nuclear speckle rehabilitation to achieve the maximum effect in alleviating proteinopathy.

### PP protects against mouse retina degeneration *ex vivo* and alleviates tauopathy in *Drosophila*

Next, we assessed the efficacy of PP in ameliorating proteinopathies by utilizing animal models of RP and tauopathy. To determine whether PP has the potential to restore gene expression changes in the retina of *Rho^P23H/+^* mice, we performed RNA-seq in the retina of one and three months old wild-type and *Rho^P23H/+^*mice (including both sexes) and compared the gene signatures of *Rho^P23H/+^* retina with that of PP (**Fig. 6a-e**). In the retina of one month-old *Rho^P23H/+^* mice, we observed a significant downregulation of protein transport and autophagy gene expression that showed large convergence with those upregulated by PP (**Fig. 6a, b**). This includes *Reep6* gene, which regulates protein trafficking in the ER and its loss-of-function mutation causes autosomal-recessive RP in both humans and mice (**Fig. 6e**) ^80^. By three months, the downregulation of proteostasis gene expression persists in *Rho^P23H/+^* mice retina, concomitant with a significant increase of YAP1-mediated cell dynamics gene expression that also overlaps with PP-downregulated genes (**Fig. 6c-e**). To directly test the efficacy of PP in protecting against RP, we treated retina explants isolated from *Rho^P23H/+^* mice (including both sexes) with 0.2 and 0.5 µM PP for 10 days, and visible light optical coherence tomography (vis-OCT) imaging ^81^ revealed a compelling efficacy of PP in safeguarding the mouse *Rho^P23H/+^* retina explants from degeneration, represented by the preserved total retinal volume and average retinal thickness from Day 0 to 10 (**Fig. 6f-h**). Notably, the cell counts in the outer nuclear layer closely resembled that of the WT retina explant control (**Fig. 6f-j**), indicating a near complete protection against degenerative processes by PP.

**Fig. 6.**
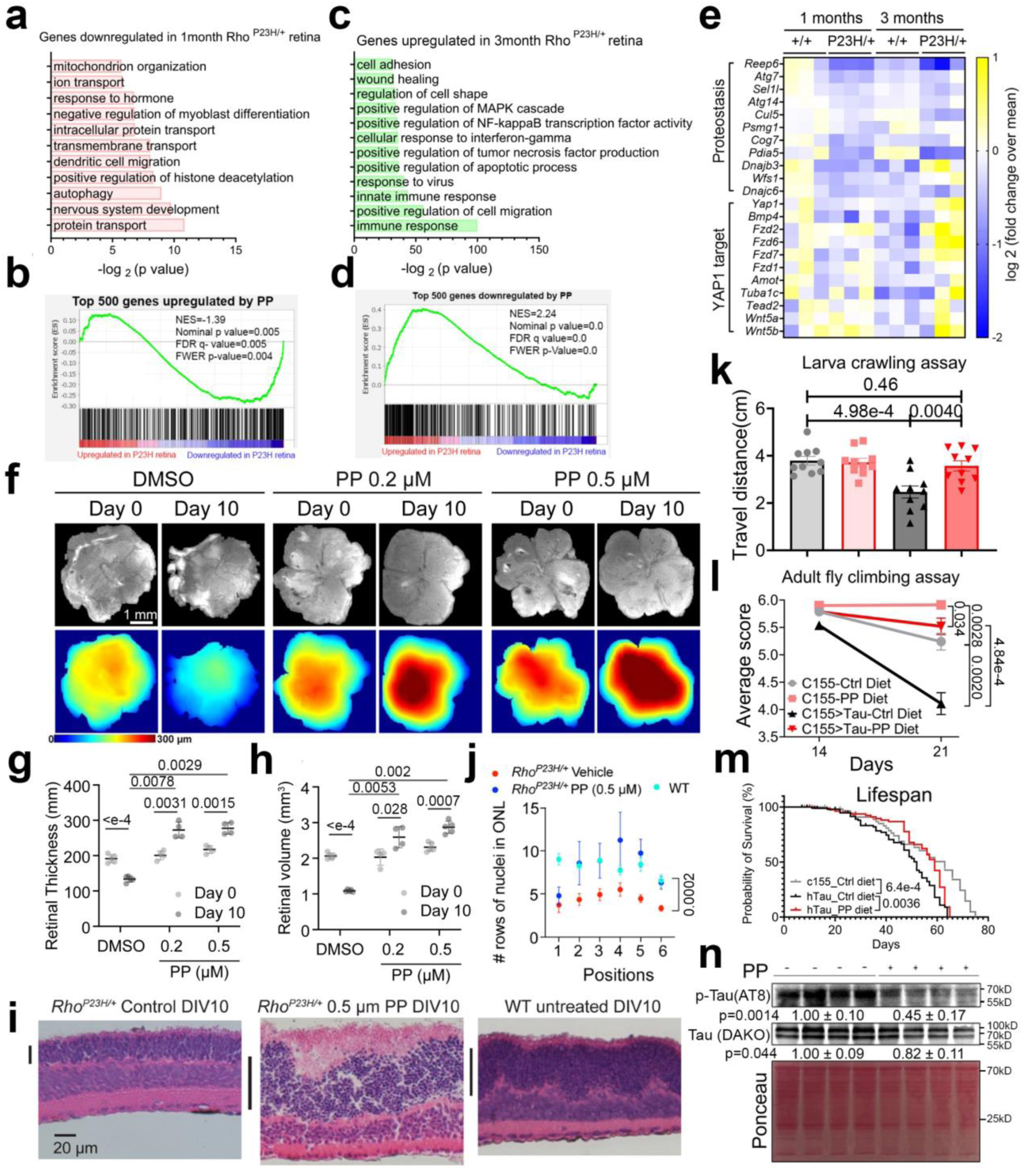
PP alleviates proteinopathies in preclinical models. (**a-e**) RNA-seq was performed in the retina of 1 and 3 months-old wild-type and Rho ^P23H/+^ mice. GO of differentially expressed genes (DEG) (FDR<0.1) (**a, c**) and GSEA comparing these DEG with that of PP (**b, d**). Heatmap of selective genes (**e**). (**f-j**) Retina explants isolated either from *Rho^P23H/+^* mice P15 and cultured with PP or DMSO vehicle control or from wild-type mice cultured for 10 days *ex vivo*. (**f**) The morphology retinae were imaged and scanned before (Day 0) and after treatment (Day 10) by a webcam (top) and visible light optical coherence tomography (vis-OCT) with tissue thickness shown as a heatmap with a color legend indicating thickness from 0-300 µm (bottom). Scale, 1 mm. (**g**) and (**h**) are bar plots of retinal thickness and volume, respectively, measured from the vis-OCT scanning data. n=4∼5 biologically independent samples. (**i**) Representative retinal histology images of the retina explants cultured for 10 days with black bars showing the outer nuclear layer (ONL). (**j**) The nuclei count in the outer nuclei layer (ONL) along six horizontal positions at peripheral-central-peripheral positions across each cross-section in (**i**). n=3-4 biologically independent samples. Similar results were confirmed in at least three independent experiments. (**k-n**) Male C155>UAS-hTau1.13 and C155 flies were fed with either standard diet or diet supplemented with 25µM PP. Quantification of distance travelled from larval crawling assay (n=10 biologically independent samples) (**k**), climbing index score from adult fly climbing assay at 14 days and 21 days (n=5 biologically independent cohorts/group) (**l**) of age, and lifespan assay (**m**) (n=32 flies for C155-Ctrl, n=65 flies for hTau-Ctrl and n=44 flies for hTau-PP). (**n**) Western blot and quantification of p-Tau (AT8 antibody) and total Tau (DAKO antibody) level in 21 days C155>UAS-hTau1.13 flies fed with control or PP diet (n=4 biologically independent samples). All data mean ± S.E.M. Statistical tests used: two-way ANOVA and Tukey multiple comparison for g and h, two-way ANOVA for j, unpaired two-tailed Student’s t-test for k, l (day 21) and n. Log-rank (Mantel-Cox) test for m. The Western blot raw blot images are provided in the source data file.

Pan-neuronal expression of wildtype human *MAPT* gene in *Drosophila* recapitulates essential features of tauopathies, including hyperphosphorylated and misfolded tau, age-dependent neuron loss, and reduced life span^82^, and SON IDR2 sequence is also conserved in flies (**Supplementary Fig. 15g**). Thus, we next tested the efficacy of PP in ameliorating tauopathy in male flies that express 2N4R isoform of human Tau (*MAPT*) pan-neuronally [*elav*^c1^^55^-Gal4: UAS-hTau1.13 (C155>UAS-hTau1.13)] as well as in control *elav* ^c1^^55^-Gal4 (C155) flies ^83^. Both C155>UAS-hTau1.13 and C155 flies were fed with either standard diet or diet supplemented with 25µM PP, which did not affect the normal development and growth of flies despite its effect in attenuating WNT and YAP1 signaling ^84^. We quantified disease progression with both larval crawling and adult fly climbing assay at 14 and 21 days of age. PP feeding preserved motor function in C155>UAS-hTau1.13 third instar larvae and adult flies, with their locomotor performance restored to a level similar to or even slightly higher than control C155 flies fed with a standard diet (**Fig. 6k, l**). Notably, PP also enhanced the motor function of adult wild-type control (C155) flies at 21 days of age (**Fig. 6l**). This improvement is likely linked to PP’s ability to promote overall proteostasis, particularly protein turnover rates, a process known to prolong health and lifespan in flies ^85^. PP further extended the median lifespan of C155>UAS-hTau1.13 flies by 16% from 51 to 59 days (**Fig. 6m**). Consistent with the overall phenotypes, PP significantly reduced the level of p-Tau and total Tau in the brains of 21 days-old C155>UAS-hTau1.13 flies (**Fig. 6n**).

### PP has the potential for treating tauopathy in humans

To determine the potential of PP for treating tauopathy in humans, we studied whether gene expression signatures that are opposite of PP can be observed in brain regions of human subjects with Alzheimer’s diseases (AD). We initially performed a post-hoc analysis of a total of nineteen bulk RNA-seq datasets encompassing hippocampus, entorhinal cortex, temporal cortex and frontal cortex regions in control and AD human subjects ^86^, and found that genes repressed by PP have increased expression in all four brain regions of human subjects with AD (such as *YAP1, TEAD1 and AMOT*) (**Supplementary Figs. 23 and 24a**). By contrast, genes that were upregulated by PP displayed significantly decreased expression in temporal cortex (such as genes involved in ERAD: *EDEM1, SEL1L*, autophagy: *ATG13*, protein folding: *HYOU1*, and tRNA aminoacylation: *GARS*, *IARS*) (**Supplementary Figs. 23 and 24a**). To validate these findings, we further analyzed an independent single-nucleus RNA-seq (snRNA-Seq) dataset in the prefrontal cortex regions of human individuals with varying degrees of AD pathology (**Supplementary Fig. 24b**) ^87^. We found that proteostasis genes upregulated by PP are consistently downregulated in all cell types with strong prominence in neurons and oligodendrocytes in both early and late-stage human AD subjects (**Supplementary Fig. 24c, d**). Genes that are downregulated by PP (those enriched in regulation of cell dynamics by YAP1) are initially downregulated in all cell types in the early stage but significantly upregulated during the late stage of AD in all cell types but inhibitory neurons (**Supplementary Fig. 24e, f**). This early to late AD progression is concomitant with strong increase of tauopathy, but not the amyloid burden in these individuals (**Supplementary Fig. 24b**). Consistent with *in vivo* data, we also observed a significant decrease of proteostasis gene expression as well as an increase of YAP1-TEAD2 target gene expression in human induced pluripotent stem cells (iPSC)-derived neurons that express the P301S 4R-Tau, compared to wild-type 4R-Tau control cells (**Fig. 7a-c**) ^73^. These upregulated YAP1-TEAD2 target genes also overlap with those repressed by PP (**Fig. 7b**).

**Fig. 7.**
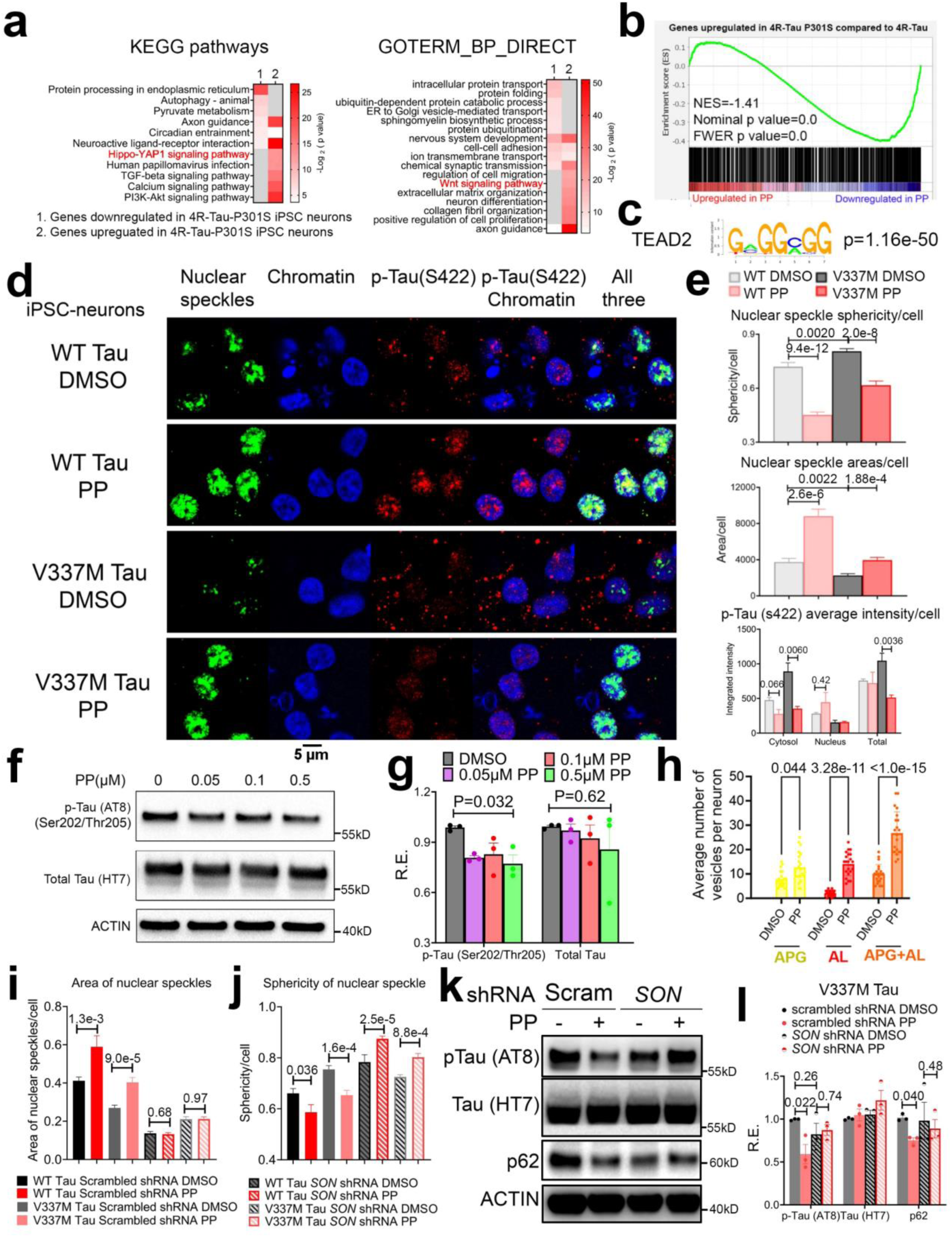
PP rejuvenates nuclear speckles and alleviates tau burden in human iPSC-neurons expressing mutant Tau. (**a**) GO analysis of up and downregulated genes in 4R-Tau P301S iPSC neurons reported in ^73^. (**b**) GSEA comparing genes upregulated in 4R-Tau P301S iPSC neurons with those downregulated by PP. (**c**) Motif analysis of promoters of genes upregulated in 4R-Tau P301S iPSC neurons compared to 4R-Tau (FDR<0.05) revealed top enriched motif of TEAD2. (**d, e**) Wild-type and V337M Tau-expressing iPSC-neurons were treated with DMSO or PP (10nM) for 12 hours, and IF against nuclear speckle (Ab11826 against SRRM2), p-Tau (Ser422) and chromatin (DAPI) were performed. Representative images (**d**) and quantification of nuclear speckle sphericity and area, and intensity of nuclear, cytosol and total level of p-Tau (Ser422) (**e**) (n=17∼32 cells collected evenly across four separate wells for each treatment from two independent experiments). (**f, g**) V337M Tau-expressing iPSC-neurons were treated with DMSO or increasing concentrations of PP for 12 hours. Representative western blot (**f**) and quantification (**g**) of total and p-Tau (Ser202/Thr205) (n=3 biologically independent samples). (**h**) V337M Tau-expressing iPSC-neurons were treated with DMSO or 100nM PP for 12 hours, and autophagy flux was quantified by the autophagy reporter (n=21 cells collected evenly across three separate wells for each treatment from three independent experiments). (**i, j**). WT and V337M Tau iPSC-neurons were infected with scrambled or SON shRNA lentivirus and treated with DMSO or 100nM PP for 12 hours, and IF against SRRM2 were performed. Quantification of the area (**i**) and sphericity of nuclear speckles (**j**) (n=10∼29). (**k, l**) V337M Tau iPSC-neurons were infected with scrambled or SON shRNA lentivirus and treated with DMSO or 100nM PP for 12 hours, and western blot analysis was performed. Representative blot (k) and quantification (l) of the level of different proteins (n=3 biologically independent samples). All data mean ± S.E.M. Statistical tests used: one-way ANOVA for g, unpaired two-tailed Student’s t-test for e, h, I, j and l. All biologically independent samples used for Western blot analyses were collected from three independent experiments. The corresponding raw blot images are provided in the source data file.

Finally, to directly test the efficacy of PP in reducing tauopathy in human, we utilized human iPSC-neurons harboring homozygous FTD-causing MAPT V337M mutation (herein referred to as V337M) and isogenic wild-type control cells ^88^ and treated both cell lines with 100nM PP for 12 hours, which did not induce toxicity or cellular stress (**Supplementary Fig. 25a-d**). Immunofluorescence against nuclear speckles marker SRRM2 revealed that compared to wildtype Tau controls, V337M Tau iPSC-neurons exhibited aberrant nuclear speckle morphology characterized by smaller size and more spherical shape, and 12 hours of 10nM PP treatment fully restored nuclear speckle morphology to normal size and diffuseness (**Fig. 7d, e**). Consequently, PP markedly reduced the level of V337M p-Tau, concomitant with increased autophagic flux (**Fig. 7d-h**). To confirm that PP rejuvenates nuclear speckles and reduces V337M p-Tau in a SON-dependent manner, we knocked down SON using lentiviral shRNA and repeated the experiment. As demonstrated in **Fig. 7i-l and Supplementary Fig. 25e-h**, PP was unable to rejuvenate nuclear speckles, enhance autophagy, or lower p-Tau levels in V337M Tau iPSC neurons in the presence of SON knocking down. Compared to V337M Tau, PP and SON had no effects on modulating wild-type Tau levels (**Fig. 7d, e and Supplemental Fig. 25g, h**), again demonstrating the lack of toxicity associated with nuclear speckle rehabilitation. Taken together, these findings indicate that rehabilitation of nuclear speckles by PP has great potential to normalize gene expression patterns and reduce Tau burden in AD/ADRD-affected humans with severe tauopathy. Further, these results provide strong support for the decline of nuclear speckle function as a driver of tauopathy.

## Discussion

Several recent studies indicate that both the decline of nuclear speckle functions and dysregulated mRNA splicing are associated with proteinopathies in humans, including tauopathy, RP and amyotrophic lateral sclerosis (ALS) ^89, 90, 91, 92, 91, 93, 94^, thus providing the rationale for nuclear speckle rehabilitation as a strategy for counteracting various proteinopathies. Exploring the therapeutic potential of targeting biomolecule condensates represents an exciting avenue for research and drug development ^95, 96^. Our study is proof of principle demonstrating that nuclear speckle LLPS can also be therapeutically targeted. Manipulating nuclear speckle LLPS through SON overexpression is a conceptually viable approach. However, the practical implementation of this strategy presents significant challenges due to the large size of the human SON open reading frame, making it technically difficult to design gene therapy targeting SON. That said, we cannot rule out the possibility that overexpression of specific truncated SON domains, such as SON IDR2, may be sufficient to rejuvenate nuclear speckles, and future efforts will be directed toward exploring such possibilities.

Through a high-throughput drug screen, we identified PP as a small nuclear speckle rehabilitator by directly interacting with SON and modulating nuclear speckle LLPS dynamics. While we don’t yet know the full detailed mechanisms by which PP modulates nuclear speckles dynamics and boosts proteostasis gene transcription, several lines of evidence suggest that it does so in part by reducing the surface tension and consequently increasing the surface areas of nuclear speckles via a SON IDR2 (but not SON IDR1)-dependent manner. Our in vitro reconstitution system further showed that reduced nuclear speckles surface tension by PP further facilitates nuclear speckles wetting of chromatin ^47^. Thus, since spliceosomes reside at the interfacial boundary between nuclear speckles and nucleoplasm/chromatin^45^, larger surface areas also entails a higher probability of spliceosome stably engaging in mRNA processing and transcription elongation. The mechanisms by which PP reduces the surface tension of nuclear speckles remain incompletely understood. Findings from our optical tweezer experiments suggest that PP does not interact with each SON IDR2 molecule in a site-specific manner. Instead, the local physicochemical properties of the SON IDR2 condensate—particularly its abundance of positively charged arginine and negatively charged aspartate and glutamate residues—likely facilitate the interaction of PP with SON IDR2 condensate as a collective unit via weak multivalent interactions. These interactions likely occur between the aromatic rings of PP and the positively charged arginine residues (through π-cation interactions) as well as the positive charged pyridinium nitrogen of PP and negatively charged amino acids on SON IDR2 (via electrostatic interactions). The collective strength of these newly formed attractive interactions is likely weaker than the native attractive cohesive forces (between D/E and R, for example) within SON IDR2 condensate alone, leading to a reduction in surface tension. The importance of arginine in nuclear speckle condensation (which is not enriched in SON IDR1), has been underscored in previous research ^52^.

Nuclear speckles play a vital role in coordinating the opposing changes observed in proteostasis and cell dynamics regulation. Elevated SON expression facilitates increased physical interactions between nuclear speckles and XBP1s, leading to augmented transcription of proteostasis genes. To explore the possibility of a direct interaction between XBP1s and SON, we employed Alphafold3^97^ and obtained iPTM and pTM scores of 0.21 and 0.22, respectively—both significantly below the recommended confidence thresholds of 0.6 and 0.5. These results suggest that direct interaction between XBP1s and SON is unlikely. Further research is needed to elucidate the precise mechanism by which XBP1s is recruited to interact with rejuvenated nuclear speckles. On the other hand, much less is clear on how nuclear speckle rehabilitation represses YAP1 transcription activity. Upon SON overexpression and PP treatment, we found a significantly reduced level of nuclear YAP1 protein and a lower nucleus/cytosol ratio, indicating an active nuclear exclusion of YAP1 protein. Like nuclear speckles, YAP1 can also form biomolecular condensates, and a recent study reported that YAP1 nuclear condensates and the nuclear speckles showed limited nuclear co-localization ^98^, suggesting a low level of wetting of these two condensates under normal physiological conditions. Pending further investigation, we speculate herein that nuclear speckles rehabilitation may further reduce the wetting of these two condensates, resulting in the alteration of YAP1 condensate composition, and ultimately its nuclear exclusion.

While both proteostasis and YAP1 signaling are downstream of nuclear speckles, direct antagonistic reciprocal interactions between these two are likely to be present as well (**Supplementary Fig. 8e**). A recent study reported that in *Drosophila*, the proteostasis output gene Bip can sequester the fly YAP1 ortholog Yorkie, in the cytoplasm to restrict Yorkie transcription output ^99^. Conversely, in undifferentiated pleomorphic sarcoma, YAP1 can suppress PERK and ATF6-mediated UPR target expression, and treatment with the YAP1 inhibitor Verteporfin upregulated the UPR and autophagy ^100^. The latter is further consistent with our findings showing that restoring YAP1 activity dampened the efficacies of PP on activating protein quality control gene expression and reducing proteinopathies. These findings indicate that in order to maximize the effectiveness of nuclear speckle rehabilitation, it is crucial to maintain elevated levels of protein quality control while simultaneously reducing YAP1 activity. Thus, a delicate balance between protein quality control and YAP1 activity appears essential for effective nuclear speckle rehabilitation. This observation may also explain why therapies merely aimed at activating protein quality control pathways often have limited efficacies.

## Materials and methods

### Mice

For retinal explant studies, littermates of wildtype C57BL/6J and Rho*^P23H/+^* knock-in mice (including both sexes) (Jackson Laboratory Strain #017628) were euthanized by CO2 and retinae were isolated for culture. For aging study, C57BJ/6J mice (including both sexes) were housed under regular 12L/12D cycles fed *ad libitum*. Liver tissues were harvested at different ages for qPCR, Western blot and immunofluorescence studies. The animal studies were carried out in accordance with the National Institutes of Health guidelines and were granted formal approval by the University of Pittsburgh’s Institutional Animal Care and Use Committee (approval numbers IS00013119, IS00023112, IS00025267 and IS00020197).

### Larva crawling assay

PP solubilized in DMSO were diluted directly into the fly medium at the final concentration of 25 µM and vortexed extensively to obtain homogeneous culture. Crawling assays were performed on 1.5% agarose plates made with a 2.3:1 combination of grape juice and water. A sample size of 10 to 15 larvae were selected for each genotype and assays were done using larvae in the third instar state. The larvae were first removed from vials and gently placed into a petri dish containing deionized water to allow for residual food to be washed off the body. After 15 seconds, the larvae were transferred to a petri dish containing the 1.5% agarose mixture and were given one minute to rest. They were then transferred to a second dish filled with the 1.5% agarose mixture and timed immediately for one minute, during which their crawling performance was measured. A transparent plastic lid was placed on top of the plates and the crawling path of the larvae were traced. Observations of the crawling activity were done under a light microscope. The brightness and distance of the light source above the plates were kept constant across all trials and genotypes. The crawling paths of the larvae were measured using FIJI ImageJ and the average distance traveled was taken for each genotype.

### Adult fly climbing assay

Male adult Drosophila melanogaster flies at 14 and 21 days of age fed with a normal diet or diet supplemented with 25 µM PP were used for assessing climbing ability. Flies were grouped into cohorts of the same sex, pre-mated, and age-matched, with a maximum of 20 individuals per vial (usually 5-15). All flies used in each trial were hatched within a 3-day window. The evening prior to each assay, flies were gently transferred to fresh tubes to allow for grooming and access to food. To ensure consistent conditions, assays were conducted at approximately the same time of day with a consistent ambient light setting. A custom climbing vial was employed, divided into six compartments, each labeled with a number (1 to 6) to denote climbing speed. The vial was positioned against a white background to enhance visibility during photography. Flies were transferred from their housing vial to the climbing vial, which was covered with a plastic plate on top. To initiate the assay, the flies were gently tapped to the bottom of the vial and allowed 10 seconds to climb. A cell phone camera was used to capture a photograph of the vial. Care was taken to ensure the camera was level with the vial, all flies were visible, and the background was free from stains or spots. The number of flies in each compartment of the climbing vial was counted at each time point and recorded on a dedicated worksheet. Each cohort of flies underwent five consecutive trials, with approximately 1 minute of rest between each trial. The average score of each cohort was determined by dividing the total score by the total number of flies.

### Fly lifespan Assay

Male Tau flies (C155-Gal4>UAS-htau1.13) and control (C155-Gal4) flies were collected from the crossing vials and separated by sex, genotype, and diet. For each condition/group, 20 flies were collected into one vial and there were 5 vials for each condition, which makes a total of 100 flies in each group. Day 0 was defined by the day that new adult flies were hatched. Flies were transferred to vials containing control diet and PP diet (25µM) on Day 1-2 and transferred to a fresh vial every two days. The number of deaths in each vial was checked every two days. Accidental loss of flies was censored from the analysis. The survival curve was graphed in GraphPad with the Kaplan-Meier analysis.

### Fibroblast cell culture and drug treatment

SV40-immortalized MEFs ^101^ and Rho P23H-expressing NIH 3T3 cells ^69^ were cultured at 37°C and 5% CO_2_ in Dulbecco’s Modified Eagle’s Medium (DMEM, glucose 4.5 g/L with phenol red) and supplemented with 10% fetal bovine serum (FBS), 1 mM sodium pyruvate (Gibco), and penicillin (100 U/mL)-streptomycin (100 μg/mL) (Gibco). Methods for the manipulation of *Son* (transient knockdown or constitutive overexpression) and validation of changes to protein (SON) levels with regards to the mRNA-Seq data are previously described in ^9^. For Tu treatment, 100 ng/mL Tu (in DMSO) for six hours was used unless otherwise noted. NB (HY-50904), PB (HY-12047), PH (HY-B0883), PP (HY-A0293), MG-132 (HY-13259), SBI-0206965 (HY-16966) and XMU-MP-1 (HY-100526) were purchased through MedChemExpress and BafA (1334) were purchased from Tocris. All drugs were handled per manufacturer instruction.

### siRNA Transient Transfections

MEFs were transfected with 10µM of different siRNAs for 24∼48 hours with Lipofectamine RNAiMAX reagents (Life technologies) per the manufacturer’s instructions. Sources of siRNA are as follows: siGENOME non-targeting siRNA pool (Dharmacon, D-001206-1305), siGENOME SMARTpool son siRNA (Dharmacon, L-059591-01-0005), siGENOME SMARTpool Stk3/Mst1 siRNA (Dharmacon, L-040440-00-0005), and siGENOME SMARTpool Stk4/Mst2 siRNA (Dharmacon, L-059385-00-0005).

### Primary neuron cell culture and P301S-Tau virus infection

The cerebral cortices of 3-4 neonate mice (P0) were dissected on ice, the meninges were removed and placed in the cold dissection medium (DM), consisting of 6 mM MgCl2 (Sigma M1028-100 ml), 0.25 mM CaCl_2_ (Sigma C7902), 10 mM HEPEs (100X), 0.9% Glucose, 20 µM D-AP5 (Cayman, NC1368401), and 5 µM NBQX (Tocris Bioscience, 10-441-0). After dissection, the brain tissues were washed with DM 1∼2 times and incubated with 13mL of DM containing papain (Worthington, LK003176) in 37°C water bath for 20 min. The suspension was shaken every 5 min. 10mL media containing 18 ml DM + 2 ml low OVO + 133 ul DNase I (dilute 10X low OVO and 150x DNase I to DM) were added into the suspension to stop the digestion in 37°C water bath for 5 min. The solution was taken off and 10 ml fresh solution was added in. Then the tissues were triturated until there were no visible chunks, and the solution was filtered through the 70 µm cell strainer. The cell solution was then centrifuged at 1000 rpm for 10 min and the supernatant was discarded. The cell pellet was gently resuspended in 20 ml B27/NBM/High glucose media, and the suspension was centrifuged at 850 rpm for 5 min. After that, the supernatant was taken off and B27/NBM (1 ml/mouse brain) was added to resuspend the cells until single cell solution. The cells were counted and plated onto the coverslips at 250k in 24-well plates for imaging or 800k in 12-well plates for qRT-PCR or Western blots. AAV-P301S hTau (Viro-vek) were infected at DIV1 at 100 MOI.

### Human iPSC-derived neurons culture

Human iPSC-derived neurons were pre-differentiated and differentiated as described ^88^. Briefly, iPSCs were pre-differentiated in Matrigel-coated plates or dishes in N2 Pre-Differentiation Medium containing the following: KnockOut DMEM/F12 as the base, 1× MEM non-essential amino acids, 1× N2 Supplement (Gibco/Thermo Fisher Scientific, cat. no. 17502-048), 10 ng/ml of NT-3 (PeproTech, cat. no. 450-03), 10 ng/ml of BDNF (PeproTech, cat. no. 450-02), 1 μg/ml of mouse laminin (Thermo Fisher Scientific, cat. no. 23017-015), 10 nM ROCK inhibitor and 2 μg/mlof doxycycline to induce expression of mNGN2. After 3 d, on the day referred to hereafter as Day 0, pre-differentiated cells were re-plated into BioCoat poly-D-lysine-coated plates or dishes (Corning, assorted cat. no.) in regular neuronal medium, which we refer to as +AO neuronal medium, containing the following: half DMEM/F12 (Gibco/Thermo Fisher Scientific, cat. no. 11320-033) and half neurobasal-A (Gibco/Thermo Fisher Scientific, cat. no. 10888-022) as the base, 1× MEM non-essential amino acids, 0.5× GlutaMAX Supplement (Gibco/Thermo Fisher Scientific, cat. no. 35050-061), 0.5× N2 Supplement, 0.5× B27 Supplement (Gibco/Thermo Fisher Scientific, cat. no. 17504-044), 10 ng/ml of NT-3, 10 ng/ml of BDNF and 1 μg/ml of mouse laminin. Neuronal medium was half-replaced every week. For SON knocking down, differentiated iPSC neurons were infected with lentivirus encoding either scrambled shRNA (pLKO.1 Puro shRNA Scramble, addgene #162011)^102^ or human SON shRNA (pLKO.1-shSON(3’UTR): Sigma TRCN0000083723) for seven days before treatment with DMSO or PP.

### Efficacy test of PP in retina explant culture

Wild type and *Rho^P23H/+^* mice were euthanized at P15, and retina explants were isolated and cultured as previously described^103, 104^. Briefly, eyeballs were enucleated and incubated in Ames solution containing 0.22 mM L-cysteine (Sigma-Aldrich) and 20 U papain (Worthington, Freehold NJ, USA) at 37 °C for 30 min. The digestion was stopped by transferring the eyes to Dulbecco’s modified Eagle’s medium (DMEM; Gibco) containing 10% fetal calf serum (FCS; Gibco) and penicillin & streptomycin antibiotics (1x, GenClone) at 4 °C for 5 min. The eye cup was made by gently removing the cornea, iris and lens. Each eye cup was flattened by four radio cuts and the sclera was then carefully peeled off from the retina:RPE complex. The retina:RPE explant was transferred to a trans well insert with 0.4-micron pore polycarbonate membrane (ThermoFisher) sitting on the surface of 1.5 mL of neurobasal-A plus medium (Gibco) containing 2% B27 supplement (Gibco) in a 6-well cell culture plate, and the RPE layer was facing the transwell membrane. The retinal explants were cultured at 37 °C with 5% CO_2_. The medium was replaced with fresh medium containing 0.5 µM PP after 24 h, which was replaced again every 2 days until 10 days in culture (DIV). A visible light optical coherence tomography (vis-OCT) prototype ^105^ was utilized to monitor the explants noninvasively at day 0 and day 10. Retinal layers were segmented automatically using a deep learning method and then manually corrected by a customized software to calculate the retinal thickness. Retina explants were collected at 10 DIV and processed for fixation, dehydration, paraffin embedding, cross-sections, dewaxing, rehydration and hematoxylin and eosin (H&E) staining^106^. H&E-stained slides were imaged by regular light microscopy with a color camera, and the number of nuclei in the outer nuclear layer (ONL) was calculated manually.

### Autophagy reporter assay

To express the LC3 reporter in the neurons, the primary mouse neuronal cultures were infected with the homemade lentivirus-mCherry-GFP-LC3 for 7 days. The florescent signal from the vacuoles at different stages were acquired by confocal imaging. The mCherry-GFP-LC3 fluorescence images were acquired with a Leica TCS SP8 confocal system using 63x oil-immersion objective. 488 nm and 568 nm laser were used to excite the GFP and mCherry, respectively. Images were taken with the same confocal settings. Minor image adjustment (brightness and/or contrast) was performed in ImageJ. The GFP and mCherry signal collected were merged into one image to quantify the red, green, and yellow vacuoles for quantification of different types of vacuoles. The different colored fluorescent signal was manually counted in each cell, and each point represents the average number of the specific vacuole for one cultured cell. For autophagy reporter assay in MEFs, pCDH-EF1a-mCherry-EGFP-LC3B was a gift from Sang-Hun Lee (Addgene plasmid # 170446; http://n2t.net/addgene:170446; RRID:Addgene_170446)^107^ and purchased from addgene. Lentivirus was packaged from HEK293T cells as previously described ^20^ and was used to infect MEFs with a MOI of 3 three times. The quantification was performed essentially the same way as in neurons.

### TEAD luciferase reporter assay

MEFs with the CRISPRa system either overexpressing *Son* or serving as controls used in this assay were previously described in ^108^. Briefly, cells were seeded at a density of 7000 cells per well in a 96-well plate with a clear bottom and white walls. Cells were then transfected using the Lipofectamine 3000 transfection kit (ThermoFisher #L3000015) for 22 hours with the 8xGTIIC-luciferase plasmid, a gift from Stefano Piccolo (Addgene plasmid # 34615; http://n2t.net/addgene:34615; RRID:Addgene_34615) ^109^. The Dual-Glo Luciferase Assay System (Promega #E2920) was used with a SpectraMax i3x plate reader (Molecular Devices) to measure firefly luciferase signal (500ms integration time, 1mm from the plate read height).

### Proteasome activity assay

The proteasome activity assay was performed per manufacturer’s instruction (Sigma Aldrich, MAK172). Briefly, MEFs transfected with scrambled control or *Son* siRNA were treated with DMSO or 100nM PP for 46 hours. MEFs were then treated with DMSO (vehicle) or 100 nM PP for 24 hours in phenol-free DMEM. Assay reagents were then added directly to the cells. Proteasomal activity was then measured using the Cell-Based Proteasome-GloTM Chymotrypsin-Like Cell-Based Assay (Promega #G8660) per manufacturer instructions using a SpectraMax i3x plate reader (Molecular Devices) to measure total luminescence (500ms integration time, 1mm from the plate read height). The final signal was corrected by subtracting the luminescence background of the relevant blanks (media with either DMSO or 100 nM PP and without cells).

### Scratch assay

Cells were grown until they were 100% confluent, ER stress was induced as previously described, and then a single scratch was performed with a pipette tip per well. Cells were imaged immediately after scratching (0hr) and then after 23hr. The Cell Profiler ^110^ “Wound Healing” pipeline (https://cellprofiler.org/examples) was used to measure the “Percentage of Gap Filled”.

### Lactate Dehydrogenase (LDH) cytotoxicity assay

LDH-Glo™ Cytotoxicity Assay (Promega, J2380) was used to examine neural damage and death by measuring Lactate Dehydrogenase (LDH) release to the media. iPSC-neurons were changed to 1mL media by replacing half of the old media with fresh media 24 hours before the drug treatment. At the end of the drug treatment, the assay was performed according to the manufacturer’s protocol. Briefly, 50uL of the cell media in 1:10 dilution was transferred to 96-well opaque flat bottom plate and 50uL of the LDH Detection Reagent was added. After 60-minute incubation at room temperature in the dark, luminescence was recorded. For calculation of the results, background from blank media was subtracted and maximum LDH release was determined by lysing with 0.2% Triton for 15 minutes.

### Immunoblot

Different cells were harvested and fractionated to produce cytosolic and nuclear lysates using the NE-PER kit (Thermo Fisher Scientific). For whole cell lysates, cells were lysed in RIPA buffer. Both protease and phosphatase inhibitors were included in the respective lysis buffer. ∼47 μg of protein was separated on a 4%-15% gradient SDS-polyacrylamide gel (Bio-Rad) which were transferred to nitrocellulose membranes, stained with Ponceau S stain, washed, blocked with 5% non-fat milk, and incubated overnight at 4°C with the following primary antibodies: anti-α-Tubulin (Cell Signaling Technology (C.S.T.) #2144), anti-Lamin A/C (C.S.T. #4777), anti-SON [Abcam #121033 (Fig. 1k and Supplementary Fig. 1c) and LSBio LS-C803664 (Fig. 3a and Supplementary Fig. 14a)], anti-YAP1 (C.S.T. #12395), anti-GFP (C.S.T. #2956), anti-Tau (DAKO) (Sigma-Aldrich, #A0024), anti-Tau (HT7) (Invitrogen, #MN1000), anti-p-Tau (S396/S404) (PHF1) (a gift from Dr. Peters Davies), anti-p-Tau (S202/T205) (AT8) (Thermo Fisher, #MN1020), anti-β-actin (C.S.T. #4970), anti-puromycin (BioLegend 381502), anti-ubiquitin (C.S.T. #58395), anti-LC3-I/II (C.S.T. #2775), anti-p62 (C.S.T. #23214, C.S.T. #5114 and Abnova H00008878-M01), anti-ATF4 (C.S.T. #11815), anti-ATF6 (Novus 70B1413.1), anti-MST1 (C.S.T. #3682), anti-MST2 (C.S.T. #3952), anti-eIF2α (C.S.T. #5324), anti-p-eIF2α (Ser51) (C.S.T. #3597) and anti-XBP1s (BioLegend 658802). The 1D4 anti-rhodopsin antibody^111^ was obtained as a gift from Dr. Krzysztof Palczewski’s laboratory. Membranes were treated with the appropriate secondary antibody conjugated to horseradish peroxidase the following day and then ECL Prime Western Blotting Detection Reagent (Cytiva) was applied. A Bio-Rad ChemiDoc MP Imaging System was used to visualize the signal, and signal intensities were determined with ImageJ ^112^. For anti-Rhodopsin western blot, the protein samples were not boiled before loading. Since ACTIN levels may also vary under different treatments, for blots with available Ponceaus S stainings, we normalize the level of each protein to that of total ponceau staining intensity (after converting to gray scale image) of each sample lane (not just the partial ponceau staining image shown in each figure) and presented the expression as relative expression (R.E.).

### Cellular thermal shift assay (CETSA)

EGFP::SC35 MEFs with EGFP knocked into the N-terminal of mouse *Srsf2* locus (previously described in ^9^) were treated with either DMSO or 3μM PP for 50 minutes at 37°C. Cells were then trypsinized and resuspended in PBS with either DMSO or 3μM PP and 100 μL of the suspensions were distributed to PCR tubes for the thermal shift assay (three minutes at a range of temperatures). The temperatures used were: 42.0°C, 42.5°C, 43.9°C, 46.2°C, 49.3°C, 53.3°C, 57.9°C, 62.1°C, 65.2°C, 67.8°C, 69.2°C, and 70.0°C. After the samples were heated, they sat at 20°C for three minutes, were snap frozen in liquid nitrogen and thawed for three cycles to lyse the cells, and then spun at 20,000 x g for 20 minutes at 4°C. The supernatant was then removed, and immunoblotting was performed using anti-GFP (C.S.T. 2956), anti-ATF4 (C.S.T. #11815), and anti-SON (Lifespan Biosciences #LS-C803664-100), followed by appropriate secondary antibody. Band intensity on the blots were relative to the intensity of the 42°C band and were normalized so that this band’s (42°C) intensity was set equal to 1.

### Protein purification and *in vitro* droplet formation assay

Regions of SON and the entire SRSF2 protein were fused to mCherry. cDNA encoding the SON-IDR N terminal (region 1), SON-IDR C-terminal (region 2), and SRSF2 were each cloned into the expression vector pET21a (+)-Histag-mCherry (Addgene plasmid # 70719) (Niederholtmeyer et al., 2015). The plasmids obtained were transformed into C3013 *E. Coli* (NEB C3013I). Fresh bacterial colonies were inoculated into LB media containing ampicillin and grown overnight at 37°C. Overnight cultures were diluted in 500mL of LB broth with ampicillin and grown at 37°C until reaching OD 0.6. IPTG was then added to 2mM, and growth continued for 3h at 37°C. The cells were pelleted and stored frozen at −80°C. Bacterial pellets were resuspended in 15mL of Buffer A (50 mM Tris-HCl, 500 mM NaCl) containing protease inhibitors (Pierce, A32965) and 10mM imidazole. The suspension was sonicated on ice for 15 cycles of 30 sec on, 30 sec off. The lysate was centrifuged for 40 minutes at 15,000 RPM at 4°C to clear debris, then added to 2mL of preequilibrated Ni-NTA agarose beads (Qiagen cat no. 30210). The agarose lysate slurry incubated for 1.5hrs at 4°C while rocking, then allowed to flow through the column. The packed agarose was washed with 15mL of Buffer A with 10mM imidazole. Protein was eluted with 5mL of Buffer A containing 15mM imidazole, 10mL Buffer A containing 100mM imidazole, then 10mL Buffer A containing 200mM imidazole. All elutions were collected in 1mL fractions. Aliquots of the collected fractions were run on an SDS-PAGE gel and stained with Imperial Protein Stain to verify the amount and purity of the protein. Fractions containing protein were combined and dialyzed against Dialysis Buffer (50mM Tris-HCl, 500mM NaCl, 10% glycerol, 1mM DTT). Recombinant mCherry fusion proteins were concentrated and desalted to 200uM protein concentration and 125mM NaCl using Amicon Ultra centrifugal filters (MilliporeSigma cat no. UFC801024) following manufacturer’s instructions. 20uM of recombinant protein was added to Droplet Buffer (50mM Tris-HCl, 10% glycerol. 1mM DTT, 10% PEG) containing indicated final salt and pyrvinium concentrations. For droplet formation assay with nuclear extracts, nuclear extracts were isolated from either HeLa or MEFs as previously described ^113^. Briefly, cells were trypsinized, collected by centrifugation, and washed in PBS buffer. Cells were then swollen in 10 vol hypotonic buffer (10 mm Tris-HCl, pH 7.5; 1.5 mm MgCl_2_; 10 mm KCl) for 10 min on ice, collected by centrifugation at 1000 × *g*, and homogenized using loose pestle B in a glass Dounce homogenizer. Nuclei were pelleted by centrifugation at 3700 × *g* for 15 min and then resuspended in 0.5 vol low-salt buffer (20 mm Tris-HCl, pH 7.5; 1.5 mm MgCl_2_; 300 mm KCl; 0.2 mm EDTA; 25% glycerol). Nuclear proteins were extracted by drop-wise addition of high-salt buffer (same as low-salt buffer except 900 mm KCl was used), and nuclear debris were pelleted by centrifugation at 25,000 × *g* for 20 min at 4 C. Nuclear extracts were then dialyzed in BC-150 buffer (20 mm Tris-HCl, pH 7.5; 0.2 mm EDTA; 150 mm KCl; 20% glycerol) until extract KCl concentration reached 160 mm. Nuclear extracts were then cleared from precipitate by another 20-min round of centrifugation at 25,000 × *g*, aliquoted, and snap-frozen in liquid nitrogen. For droplet formation assay with HeLa nuclear extract supplementation, NE was added to different concentration of SON IDR2 at the final concentration of 1.5mg/ml in Droplet buffer (20mM HEPES, pH 7.9, 20% glycerol, 125mM KCl, 0.2mM EDTA, 0.5mM DTT, 10% DEG). For droplet formation assay with GFP::SRSF2 MEF nuclear extract supplementation, NE was added to 10uM SON IDR2 at the final concentration of 0.6mg/ml in Droplet buffer. A custom imaging chamber was created by placing strips of tape on a glass coverslip, forming a square. The protein solution was immediately loaded onto the center of the square and covered with a second glass coverslip. Slides were then imaged with a Leica confocal microscope with a 63x oil objective. The image series were taken over a 20-minute time span with 1 image every 30 seconds.

### Mass spectrometry to profile condensates composition

HeLa nuclear extract samples were thawed at room temperature, vortexed for 10 minutes, bath sonicated for 5 minutes and centrifuged at 13000g for 10 minutes at room temperature prior to quantification of total protein by a Pierce 660 Protein Assay (Thermo Scientific #22660). Condensates were obtained by centrifuging at 10,000xg for 10 minutes and resuspended in 5%SDS in 50 mM TEAB prior to total protein quantification. Protein digestion was carried out on 10 µg of protein from each sample on S-trap micro columns (Protifi) according to the manufacturer’s protocol. Following digestion, peptide samples were then dried in a speedvac and resuspended in a solution of 3% acetonitrile and 0.1% TFA and desalted using Pierce Peptide Desalting Spin Columns (Thermo Scientific # 89851). Eluants were dried in a speedvac and resusupended in a solution of 3% acetonitrile and 0.1% formic acid to a final concentration of 0.5 µg/µL. Mass spectrometry analysis was conducted on a Thermo Fisher QE-HFX coupled to a Vanquish Neo UHPLC. Approximately 1 µg of each sample was loaded onto an EASY-Spray PepMap RSLC C18 column (2 µm, 100A, 75µm x 50 cm) and eluted at 300 nl/min over a 120-minute gradient. MS1 spectra were collected at 120,000 resolution with a full scan range of 350 – 1400 m/z, a maximum injection time of 50ms and the automatic gain control (AGC) set to 3e6.

The precursor selection window was 1.4 m/z and fragmentation were carried out with HCD at 28% NCE. MS2 were collected with a resolution of 30,000, a maximum injection time of 50ms and the AGC set to 1e5 and the dynamic exclusion time set to 90s. The collected MS data were analyzed using MSFragger V4.0[1] and searched against the human SwissProt database. The search parameters were set as follows: strict trypsin digestion, missing cleavage up to 2, carbamidomethylation of cysteine as static modification, oxidization of methionine and protein N-terminal acetylation as variable modification, a maximal mass tolerance of 20 ppm for the precursor ions and 20ppm for the fragment ions, and false detection rate (FDR) was set to be 1%.

### 1,6-hexanediol treatment to examine effects of PP on LLPS

A 10% (w/v) 1,6-hexanediol (1,6-HD, MilliporeSigma) solution was prepared in Dulbecco’s Modified Eagle’s Medium (DMEM, glucose 4.5 g/L with phenol red) supplemented with 10% fetal bovine serum (FBS), 1 mM sodium pyruvate (Gibco), and penicillin (100 U/mL)-streptomycin (100 μg/mL) (Gibco). To examine NS LLPS dynamics, EGFP::SC35 MEFs (previously described in ^9^) were treated with DMSO or 1 μM PP for 30 minutes and then treated with 0, 1, 2, or 10% 1,6-HD for 20 minutes. Cells were fixed in 2% paraformaldehyde, stained with bisBenzimide H 33258 (Hoechst), and then imaged. Image analysis was completed in Cell Profiler; briefly, the process was to image the cells in the 405 (Hoescht), 488 (GFP), and 555 (high intensity to image whole cells) and then use the 405 channel to determine the nuclei boundaries, 488 to determine EGFP::SC35, and 555 to determine the area of the whole cells. We differentiated nuclear and cytosolic areas by subtracting the 405 signals from the 555 signals. 488 signal was then quantified in the aforementioned nuclear area and cytosolic area and compared. 20+ cells were measured for each condition. The average Manders coefficient was determine with ImageJ ^112^ by averaging the tM1 and tM2 values.

This same process was used to examine the effects of PP on GW182 and MED1 except in wildtype MEFs. The signal was identified by immunofluorescence (IF). Briefly, IF was performed by fixing cells with 4% paraformaldehyde in PBS, permeabilizing with 0.2% Triton X-100 in PBS, blocked with 2% bovine serum albumin in PBS, and then incubated with primary antibody (GW182 (ab156173, Abcam) or MED1 (ab60950, Abcam)) diluted per manufacturer’s recommendation overnight at 4°C. Cells were then treated with the appropriate 1:1000 secondary antibody overnight at 4°C, stained with Hoechst the following day, and then mounted with ProLong Gold Antifade (Invitrogen). Signal (either IF or endogenous GFP) sphericity was determined as previously described in ^9^. 20+ cells were measured for each condition.

### Optical tweezer C-Trap experiment

The condensate formation was essentially performed as previously described. 20 uM of mCherry-SON IDR2 was incubated in 125 mM KCl, protease inhibitor, 20 mM HEPES, 20% glycerol, 0.2 mM EDTA, 0.5 mM DTT, and 10% PEG. The total volume was 100 uL. After resuspension, samples were flowed into a custom flow-cell of a LUMICKS C-Trap and allowed 10-15 minutes to form droplets on the surface of the glass. In addition, we tested varying concentrations of our drug of interest, PP, by adding the appropriate amounts from our stock solutions to achieve final concentrations of 0 nM, 10 nM, 25 nM, and 100 nM and 500 nM. After generating condensates on the surface of the flow cell, 1.7 μm streptavidin-coated polystyrene beads were suspended in phosphate buffered saline and flowed into a separate channel of the custom flow cell from LUMICKS. During flow, the Z-position was set to approximately the center of the flow chamber, 50 μm from the surface to rapidly immobilize beads in the optical traps. After single beads were immobilized in trap one and two, the flow was stopped, and the microstage adjusted to move the traps to the channel containing condensate droplets. The Z-position was then lowered to ∼1-2 μm away from the surface of the glass, so that the beads could interact with the condensates without touching the glass surface. Trap stiffness was calibrated at this position to simultaneously measure the force exerted on the polystyrene beads by the condensates. The condensates were then screened for positive mCherry signal and roughly circular shape with diameter of ∼5-10 μm. Once identified, both trapped polystyrene beads were moved to the two opposite edges of the condensate to allow the protein to adhere, and then one trapped bead is moved away at a velocity of ∼0.5 μm/second to measure the viscoelastic properties of the condensate while simultaneously imaging in the brightfield and confocal channels. In cases without rupture, the adhesive force was sufficient to either remove the condensate from the surface of the glass or to remove the bead from the optical trap, and in cases with rupture, the condensate extended from the pulling force until it ruptured into two droplets. In cases where the condensate was removed from the surface, a second pull curve was generated by holding the condensate and bead in one trap and bringing the free bead to the position of the trapped condensate as before to again test for rupture or removal of the bead from the optical trap. mCherry-SON IDR2 was excited at 561 nm and emission collected in a 575-625 nm band pass filter. All data was collected with a 1.2 NA 60X water emersion objective and fluorescence measured with single-photon avalanche photodiode detectors. Confocal laser power was set to 5% power and continuously scanned to generate snapshots every 2-5 seconds, depending on the scan area needed. Data analysis was carried out using Lakeview software from LUMICKS and FIJI to visualize forces and extract stacks of .tiff images, respectively.

### Reverse transcription quantitative polymerase chain reaction (RT-qPCR)

For Reverse transcription-quantitative polymerase chain reaction (RT-qPCR), cDNA was produced using the Superscript III (Thermo Fisher) kit and qPCR was completed using the SYBR Green system (Thermo Fisher) in a CFX384 Real-Time System (Bio-Rad). The expression of each gene was normalized to that of *β-actin* and presented as relative expression (R.E.). The qPCR primer sequences (all are for mouse genes, unless otherwise indicated) were as follows:

*β-actin* forward: AAGGCCAACCGTGAAAAGAT

*β-actin* reverse: GTGGTACGACCAGAGGCATAC

*Amot* forward: CTGGAAGCAGATATGACCAAGT

*Amot* reverse: GGTGTTAGGAGAGTGGCTAATG

*Atf4* forward: CCACTCCAGAGCATTCCTTTAG

*Atf4* reverse: CTCCTTTACACATGGAGGGATTAG

*Atg4c* forward: GTGCGGAATGAGGCTTATCA

*Atg4c* reverse: CCAGACTTCTTCCCAAACTCTATC

*Bmp4* forward: AACGTAGTCCCAAGCATCAC

*Bmp4* reverse: CGTCACTGAAGTCCACGTATAG

*Ern1* forward: TCCTAACAACCTGCCCAAAC

*Ern1* reverse: TCTCCTCCACATCCTGAGATAC

*Fdz1* forward: GAGATCCACCTTCCAGCTTTAT

*Fzd1* reverse: CACTCCCTCTGAACAACTTAGG

*Hyou1* forward: GAGGCGAAACCCATTTTAGA

*Hyou1* reverse: GCTCTTCCTGTTCAGGTCCA

*Manf* forward: GACAGCCAGATCTGTGAACTAAAA

*Manf* reverse: TTTCACCCGGAGCTTCTTC

*Rnf166* forward: GAAGACACACTCCCGCTTTA

*Rnf166* reverse: CTGAGACCAACTCTCCTTGTG

*Sirt2* forward: CATAGCCTCTAACCACCATAGC

*Sirt2* reverse: GTAGCCTGTTGTCTGGGAATAA

*Sqstm1* forward: AACAGATGGAGTCGGGAAAC

*Sqstm1* reverse: AGACTGGAGTTCACCTGTAGA

*Tgfb3* forward: CCACGAACCTAAGGGTTACTATG

*Tgfb3* reverse: CTGGGTTCAGGGTGTTGTATAG

*Ube2q2* forward: TTCCTAAGCACCTGGATGTTG

*Ube2q2* reverse: CTCCTCCTCTTCCTCTTCTTCT

*Xbp1* forward: GGGTCTGCTGAGTCC

*Xbp1* reverse: CAGACTCAGAATCTGAAGAGG

*Cul5* forward: GAACACAGGCACCCTCATATT

*Cul5* reverse: AGTTACACTCTCGTCGTGTTTC

*Psmg1* forward: CCAGTGGTTGGAGAAGGTTT

*Psgm1* reverse: GGGTCTTGTAGTCTGTGATGTG

*Atg14* forward: CATTCCCTGGATGGGCTAAA

*Atg14* reverse: CCTCAGGAACAAGAAGGAAGAG

*Yap1* forward: CCAATAGTTCCGATCCCTTTCT

*Yap1* reverse: TGGTGTCTCCTGTATCCATTTC

Mouse *Son* forward: ttccgggaaatacaacagga

Mouse *Son* reverse: gggtggatttgtttcaccat

Human *SON* forward: GTTTAGCAGATCTCCCATCCG

Human *SON* reverse: ACACCAGCCTTAGCACAC

### Chromatin Immunoprecipitation (ChIP)

ChIP for SC35 was performed using anti-SC35 antibody (ab11826, Abcam) as previously described ^114^. Briefly, mouse liver samples were submerged in PBS + 1% formaldehyde, cut into small (∼1 mm3) pieces with a razor blade and incubated at room temperature for 15 minutes. Fixation was stopped by the addition of 0.125 M glycine (final concentration). The tissue pieces were then treated with a TissueTearer and finally spun down and washed twice in PBS. Chromatin was isolated by the addition of lysis buffer, followed by disruption with a Dounce homogenizer. The chromatin was enzymatically digested with MNase. Genomic DNA (Input) was prepared by treating aliquots of chromatin with RNase, Proteinase K and heated for reverse-crosslinking, followed by ethanol precipitation. Pellets were resuspended and the resulting DNA was quantified on a NanoDrop spectrophotometer. An aliquot of chromatin (10 μg) was precleared with protein A agarose beads (Invitrogen). Genomic DNA regions of interest were isolated using 4 μg of antibody. Complexes were washed, eluted from the beads with SDS buffer, and subjected to RNase and proteinase K treatment. Crosslinking was reversed by incubation overnight at 65 °C, and ChIP DNA was purified by phenol-chloroform extraction and ethanol precipitation. ChIP-qPCR for MEFs were essentially performed the same way as previously described with anti-SC35 (ab11826, Abcam) and anti-XBP1s antibody (Biolegend 658802), except that the MEFs were directly fixed with 1% formaldehyde before subject to nuclei isolation and chromatin immunoprecipitation. The primers used for ChIP-qPCR are as follows:

Gene desert forward primer: GCAACAACAACAGCAACAATAAC

Gene desert reverse primer: CATGGCACCTAGAGTTGGATAA

*Xbp1* promoter region forward primer: GGCCACGACCCTAGAAAG

*Xbp1* promoter region reverse primer: GGCTGGCCAGATAAGAGTAG

*Xbp1* gene body region forward primer: CTTTCTCCACTCTCTGCTTCC

*Xbp1* gene body region reverse primer: ACACTAGCAAGAAGATCCATCAA

*Manf* promoter region forward primer: ACAGCAGCAGCCAATGA

*Manf* promoter region reverse primer: CAGAAACCTGAGCTTCCCAT

*Manf* gene body region forward primer: CAACCTGCCACTAGATTGAAGA

*Manf* gene body region reverse primer: AGGCATCCTTGTGTGTCTATTT

*Hyou1* promoter region forward primer: GACTTCGCAATCCACGAGAG

*Hyou1* promoter region reverse primer: GACTTCTGCCAGCATCGG

*Hyou1* gene body region forward primer: TGGAAGAGAAAGGTGGCTAAAG

*Hyou1* gene body region reverse primer: TCCCAAGTGCTGGGATTAAAG

### HTS of FDA-approved drugs and quantitative imaging analysis

EGFP::SC35/SRSF2 MEFs (EGFP was knocked in to the N-terminus of endogenous *Srsf2* locus, which is a well-established marker for nuclear speckles ^9^) were seeded in black 384 well plate with glass bottom (Cellvis). The FDA-approved compound library (100nL per drug) was stamped to 384-well tissue culture plates using CyBio Well vario (Analytik Jena). Compound solutions were then added to the cell plate at the final concentrations of 10 μM using a BRAVO liquid handler. After 18 hours of treatment, culture media was removed, and cells were fixed with 4% PFA followed by DAPI staining. Cells were imaged using a GE INCELL 2200 with 60x lens. Maximum intensity projection images were captured, and nuclear speckles sphericity was quantified with CellProfiler as previously described in ^9^. Briefly, for speckle i the sphericity is defined as equation 1:

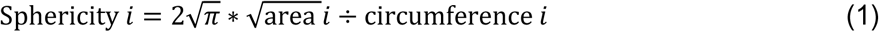

So that a perfect circle will have a sphericity of 1, and a line will have a sphericity of 0. To calculate the average sphericity of a given image that has k total speckles, we calculated the area-weighted average as described in equation 2.

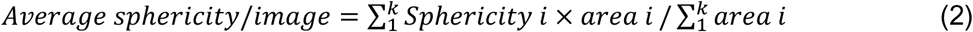

For the secondary screening to determine if compounds affected nuclear speckles morphology in a dose-dependent manner, specific compounds were selected using a TTP Mosquito X1 followed by serial dilutions of compounds that were prepared using a Bravo automated liquid handling platform (Agilent). Cells were treated, imaged, and analyzed according to the same protocol described above. For the tertiary screen we used MEFs with a *Perk* promoter-driven dGFP as previously described ^115^. Cells were treated with drugs of interest and either DMSO or Tu at the same time as described above and both GFP/cell and cell number were determined with CellProfiler as previously described ^3^. For both dose-response and *Perk* promoter-driven dGFP experiments, eight biological replicates were performed per drug per dose.

### Immunofluorescence

Immunofluorescence was performed as previously described ^116^. Briefly, liver OCT sections or cells cultured in chamber slide were fixed in cold acetone for 10 mins at −20 °C. The sections were then air dried, rehydrated with PBS and permeabilized with PBS+ 0.1% Triton X-100. The sections were then blocked with 10% goat serum at room temperature for 1 hour. For mouse liver tissues, primary antibodies against SC35/SRRM2 (Abcam, ab11826) were conjugated to Alexa-488, respectively per manufacture’s protocol and added to the OCT section at 1:1000 dilution overnight at 4 °C. Next day, sections were washed five times with PBS and counterstained with DAPI before mounting (with ProlongGold Glass) and imaging using Leica SP8 lightening confocal microscope (Leica Microsystems). For iPSC cell culture experiment, after incubation with anti-SC35/SRRM2 (Abcam, ab11826), anti-p-Tau (Ser422) (LifeTechnology, 44-764G) primary antibodies, Alex488 and Alex555-conjugated secondary antibodies were added, and the rest was performed essentially the same way.

### mRNA-seq and transcriptome analysis

For all mRNA-seq or RT-qPCR, total mRNA was isolated and purified from MEFs using the PureLink RNA Mini Kit (Thermo Fisher). For mRNA-seq, samples were submitted to the UPMC Genome Center for quality control, mRNA library preparation (Truseq Stranded mRNA (poly-A pulldown), and sequencing (paired-end 101 bp reads and ∼40 million reads per sample). The sequencing was performed on a NextSeq 2000 sequencer.

For the *son* overexpression or knockdown (OE/KD) sequencing data, the raw RNA-seq FASTQ files were analyzed by FastQC for quality control. Adaptors and low-quality reads were filtered by Trimmomatic ^117^. Then the processed reads were aligned by HISAT2 ^118^ against mouse reference mm10. For gene-level intron/exon quantification, bedtools software ^119^ was used to collect and count reads that aligned to any intron/exon of the given gene. If one read spans across multiple exons of the same gene, it will only be counted once. If one read spans intron/exon junction, it will only be counted as intron. The intron/exon count was normalized by gene length and total reads for FPKM normalization.

The analysis of drug-treated samples was done using Galaxy ^120^: Trim Galore was used for quality control ^121^ and Salmon ^122^ was used to normalize the paired-end reads (TPM method). The online 3D RNA-Seq ^123^ pipeline was used to determine upregulated and downregulated genes, in addition to generating the PCA plot.

For RNA-seq of *Rho^P23H/+^* and wild-type mice, total RNA was isolated from retinae isolated from mice at 1, 3, and 6 months of age using TRIzol organic extraction. RNA-seq was performed by QuickBiology Inc. RNA integrity was checked by Agilent Bioanalyzer. Libraries for RNA-seq were prepared with KAPA Stranded mRNA-Seq poly(A) selected kit (KAPA Biosystems, Wilmington, MA) using 250 ng toal RNAs for each sample. Paired end sequencing was performed on Illumina HighSeq 4000 (Illumina Inc., San Diego, CA).

The reads were first mapped to the latest UCSC transcript set using Bowtie2 version 2.1.0^124^ and the gene expression level was estimated using RSEM v1.2.15^125^. TMM (trimmed mean of M-values) was used to normalize the gene expression. Differentially expressed genes were identified using the edgeR program^126^. Genes showing altered expression with p < 0.05 and more than 1.5-fold changes were considered differentially expressed. Goseq was uesd to perform the GO enrichment analysis and Kobas was used to perform the pathway analysis.

### mRNA splicing rates analysis

The mRNA processing rate was estimated by the simple kinetic model (equation 3) where pre-mRNA was converted to mature mRNA with the mRNA processing rate *Kp* and the mature mRNA is subject to decay with a constant decay rate *Kd*.

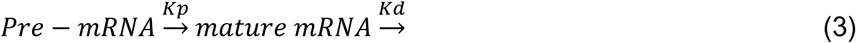

Under the basal condition (DMSO), we assume a steady state of mature mRNA expression whose level does not change over time; thus, we have:

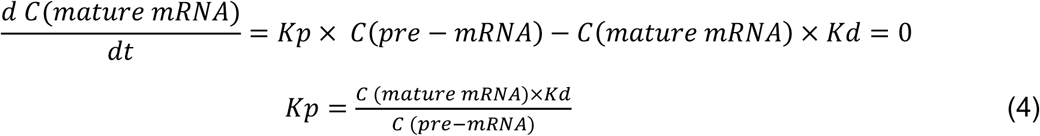

Under *Son* overexpression or knocking down condition (condition 2), if we assume the mature mRNA degradation rate does not change with *Son* OE/KD (*K1*d=*K2*d), then the ratio of splicing rates between basal condition 1 and *Son* OE/KD condition 2 is given by:

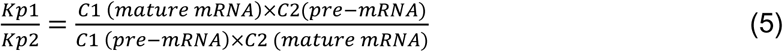

### Gene set enrichment analysis (GSEA)

GSEA was performed with software version 4.1.0. TPM quantification of transcriptome under different drugs or DMSO control was used as input for gene expression. Parameters used for the analysis: 1000 permutations, permutation type: gene set.

### Intron retention detection

Intron retention events were detected by iREAD ^34^. Intron retention events are selected with default settings T>=20, J>=1, FPKM>=2.

### Gene ontology analysis

DAVID (Version 2021) ^127^ (https://david.ncifcrf.gov) was used to perform Gene Ontology analyses. Briefly, gene names were first converted to DAVID-recognizable IDs using Gene Accession Conversion Tool. The updated gene list was then subject to GO analysis using all Homo Sapiens as background and with Functional Annotation Chart function. GO_BP_DIRECT, KEGG_PATHWAY or UP_KW_BIOLOGICAL_PROCESS was used as GO categories. Only GO terms with a p value less than 0.05 were included for further analysis.

### Motif analysis

Motif analysis was performed with the SeqPos motif tool (version 0.590) ^128^ embedded in Galaxy Cistrome using all motifs within the Mus musculus reference genome mm10 as background. LISA analysis was performed using webtool (http://lisa.cistrome.org/).

### Statistical Analysis

Data was analyzed and presented with GraphPad Prism software (version 10.0.0). All measurements were taken from distinct biological samples. Plots show individual data points and bars at the mean and ± the standard error of the mean (SEM). Individual tests used were provided at the end of each figure legend. Generally, two-tailed Student’s t-tests were used to compare differences between two groups, except in five cases where strong prior evidence supports a unidirectional change and a subsequent utilization of one-tailed Student’s t-test. Specifically, unpaired one-tailed Student’s t-tests were applied in Supplementary Fig. 21d, f, and Supplementary Fig. 22b, d, h, justified by previous data demonstrating a consistent and significant decrease in Rhodopsin levels following PP treatment in NIH3T3 RHOP23H cells (Fig. 5g–j, 7 biological replicates) and a consistent reduction in total and phosphorylated Tau levels in Tau P301S-expressing primary neurons (Fig. 5k–m, 8–12 biological replicates). One-way ANOVA was used to compare the difference of multiple treatments with one variable with equal sample size. In cases where the sample sizes were unequal, the Kruskal-Wallis test (followed by Dunn’s multiple comparison), a non-parametric alternative that does not assume normality or equal variances, was used, such as data in Figs. 2b, 3k and Supplementary Fig. 1f. Two-way ANOVA was used to compare the difference between two treatments with one additional variable, such as data in Fig. 4i. However, for data with unequal sample size, mixed effects analyses were used in lieu of Two-way ANOVA, such as data in Fig. 3a. In Supplementary Fig. 2e, Three-way ANOVA was used to determine how SON (first variable) and Tu (second variable) affect SC35 and XBP1s chromatin binding across three different regions (third variable), respectively. For multiple comparisons, we selected the appropriate method based on the comparison structure: Tukey’s test was used when comparing all row (or column) means to each other, and the Šídák method, offering slightly more power than Bonferroni, was used for a series of independent comparisons. For Fig. 3j, Chi-square test for trend, also known as the Cochran-Armitage test for trend, was used to assess whether there is a linear trend between two variables, where one variable is binary (rupture or not), and the other variable is an ordered categorical variable (increasing dose of PP). The chi-square test for trend was used to essentially answer the question whether the concentration of PP is associated with the presence or absence of condensates rupture. Log-rank (Mantel-Cox) test was used to assess lifespan differences in Fig. 6m.

## Author contributions

Conceptualization, Y.C., X.C., and B.Z.; Methodology, W.D., B.B.C., S.L., S.P., Y.C., B.V.H., X.C., and B.Z.; Investigation, W.D., Y.T., S.K., S.Z., M.C., M.A.S., R.K.A., M.S., Y.N., M.Y., I.J., M.B.L., D.C., E.I., C.D., H.H.W., S.L., S.P., B.B.C., Y.C., X.,C, and B.Z.; Writing – Original Draft, W.D., Y.C., X.C., and B.Z.; Writing – Review & Editing, all authors; Funding Acquisition, W.D., B.B.C., M.A.S., B.V.H., Y.C., X.C., and B.Z.; Resources, B.B.C. S.P., Y.C., B.V.H., X.C. and B.Z; Supervision, Y.C., X.C., and B.Z.

## Acknowledgements

We would like to thank Drs. Yvonne Eisele, Yuan Liu and Toren Finkel (University of Pittsburgh School of Medicine, Pittsburgh, PA, U.S.A.) for their technical assistance and/or comments on the manuscript. We thank Dr. Krzysztof Palczewski who generously shared the 1D4 anti-rhodopsin antibody. W.D. was supported by grant T32 HL082610 through the National Institutes of Health (NIH), the Diana Jacobs Kalman/AFAR Scholarship for Research in the Biology of Aging through the American Federation for Aging Research, and fellowship F31 AG080998 from NIA. B.B.C. was supported by grant 1R35HL139860 through the NIH. Y.C was supported by the R01 EY030991. X.C. was supported by the National Institutes of Health grants R01AG074273 and R01AG078185. B. Zhu was supported by grants 1DP2GM140924 and 1R21AG071893 through the NIH, and grants from Richard King Mellon foundation and AFAR/Hevolution foundation. This research was supported in part by the University of Pittsburgh Center for Research Computing through the resources provided. Specifically, this work used the HTC cluster, which is supported by NIH award number S10OD028483. This research project was supported in part by the Pittsburgh Liver Research Centre supported by NIH/NIDDK Digestive Disease Research Core Center grant P30DK120531, the Ophthalmology and Visual Sciences Research Center core grant P30 EY08098, the Eye and Ear Foundation of Pittsburgh and an unrestricted grant from Research to Prevent Blindness. Y.C and B.Z are also supported by Foundation Fighting Blindness Translational Research Program Award. This work was further supported by NIH R35ES031638 (BVH), F32ES034982 (MAS), S10OD032158 (BVH).

## Declaration of interests

B.B.C. is Co-founder of Koutif Therapeutic Inc., Co-founder and VP of Drug Discovery for Generian Pharmaceuticals, and Co-founder and C.S.O. of Coloma Therapeutics Inc..

## Data availability

All raw and processed sequencing data generated in this study have been submitted to the NCBI Gene Expression Omnibus (GEO; http://www.ncbi.nlm.nih.gov/geo/) under accession numbers GSE224275 and GSE281959. The mass spectrometry proteomics data have been deposited to the ProteomeXchange Consortium via the PRIDE partner repository with the dataset identifier PXD050371. All data needed to evaluate the conclusions in the paper are present in the paper and/or the Supplementary Materials.

## Code availability

No new codes were generated from this study.

## Supplementary Figures

**Supplementary Figure 1.**
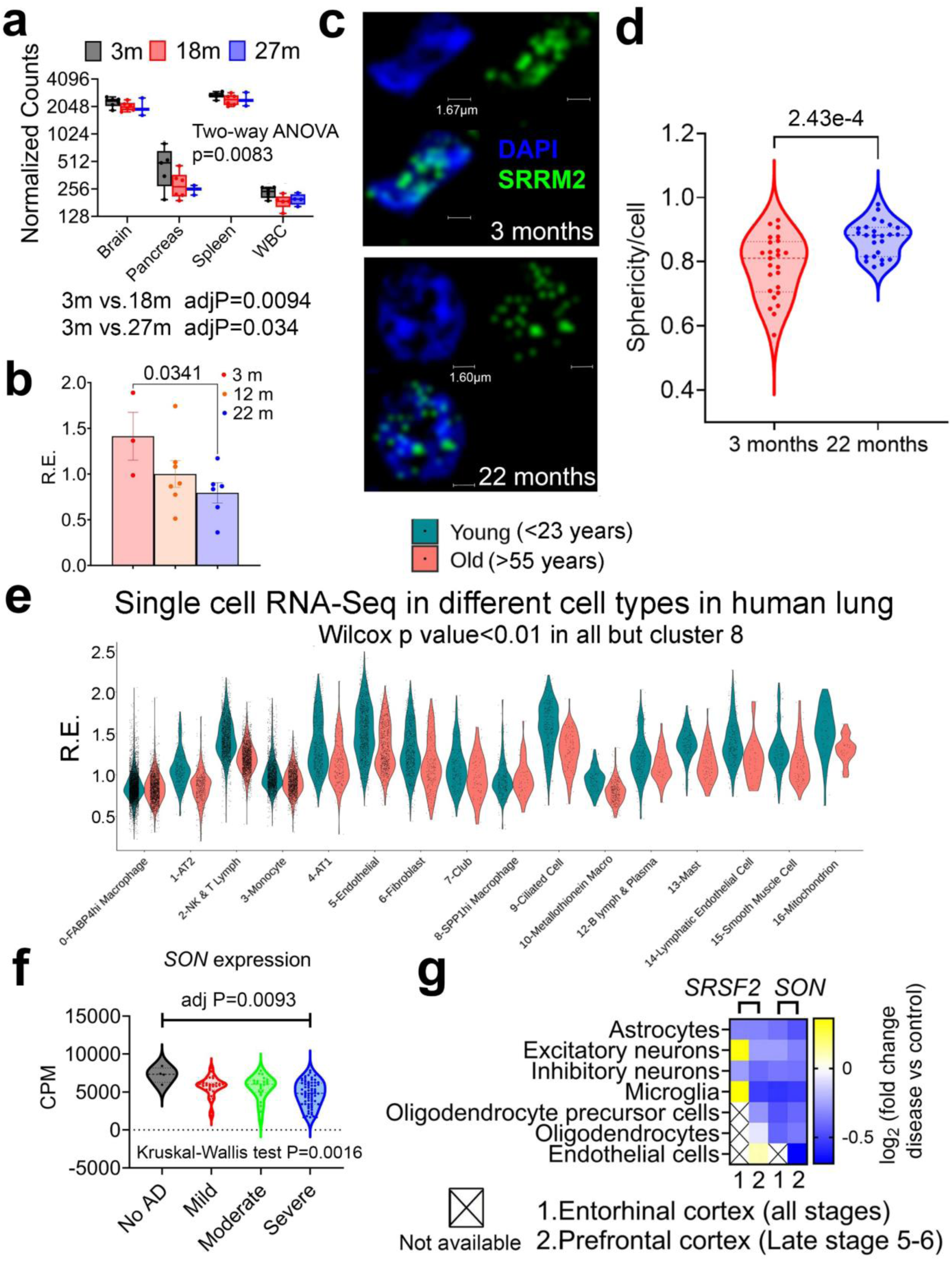
SON expression is reduced during aging in mice and humans. (**a**) Normalized counts of *Son* mRNA in different mouse tissues with different ages (WBC; White blood cells) according to Tabula Muris Senis database (n=3∼6 biologically independent samples) ^129, 130^. (**b**) qPCR of *Son* level in liver of mice with different ages (n=3∼7 biologically independent samples). (**c, d**) Representative immunofluorescence images of SRRM2 and DAPI in the liver of 3 and 22 months old male mice (**c**) and quantification of the sphericity of nuclear speckles (**d**). n=25 cells collected evenly across three mice for each age group. For b-d, both male and female mice are included. (**e**) Single cell RNA-seq data of *SON* mRNA level in different cell types in lung tissues of young and aging humans, reported from ^21^. (**f**) Counts per million normalized expressions of *SON* in brains of human AD compiled from RNA-seq data from the AMP-AD consortium (n=5 biologically independent samples for No AD, n=30 biologically independent samples for mild AD, n=24 biologically independent samples for moderate AD, and n=70 biologically independent samples for severe AD). (**g**) Heat map showing relative expression of *SRSF2* and *SON* in different cell types in the cortex of human AD subjects ^131^. Blue color indicates lower level in AD subjects compared to controls. Data: Mean ± S.E.M. Statistical tests used: Two-way ANOVA and Šídák’s multiple comparisons test for a, unpaired two-tailed Student’s t-test for b, and d. Wilcoxon test for e. Kruskal-Wallis test and Dunn’s multiple comparisons test for f.

**Supplementary Figure 2.**
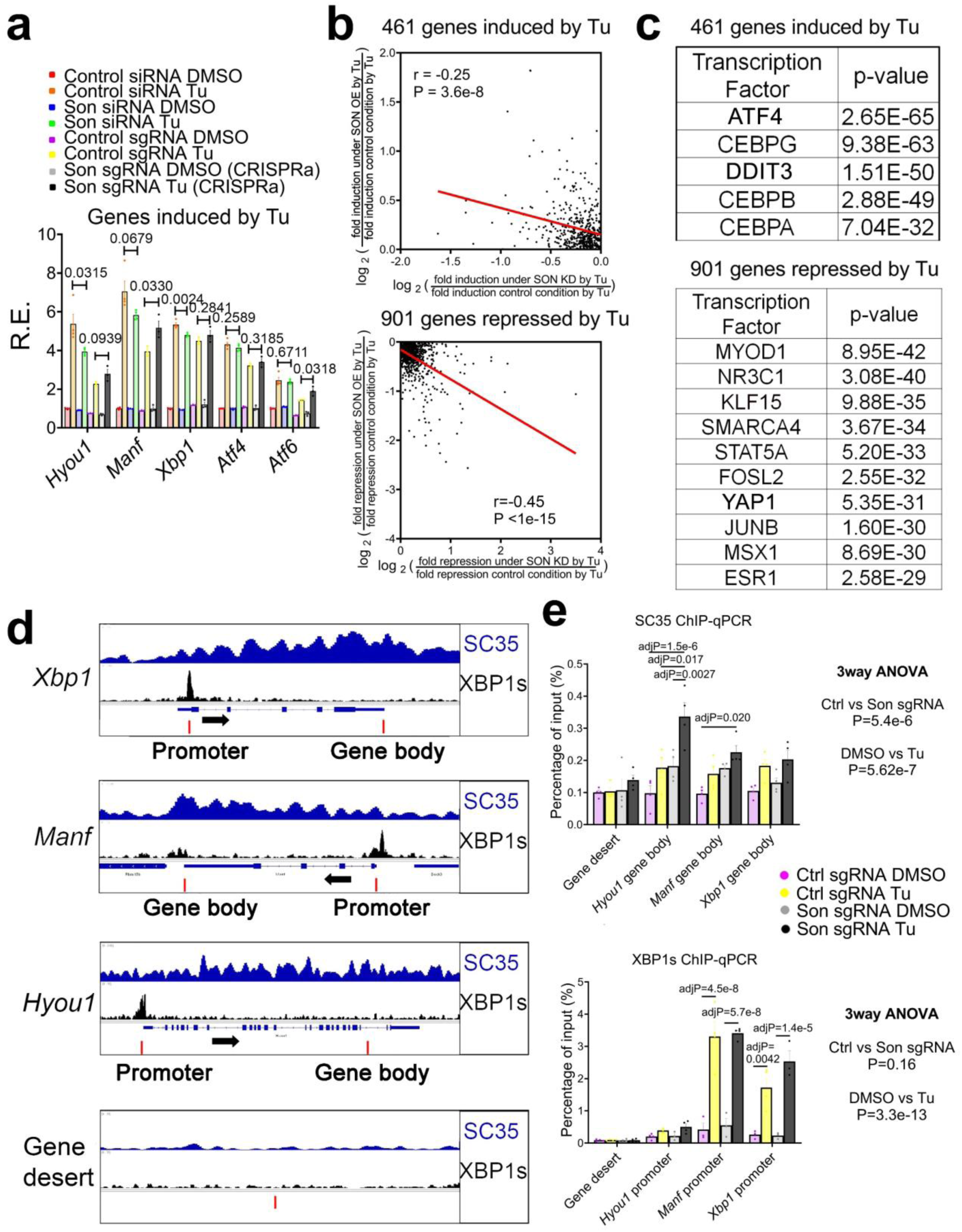
Genetic rehabilitation of nuclear speckles by SON. (**a**) Relative expression (R.E.) of representative proteostasis genes (top) and YAP1 target genes (bottom) in response to Tu (100ng/ml for 6h) in the presence of SON OE/KD (n=4 biologically independent samples for SON KD and n=3 biologically independent samples for SON OE). (**b**) Scatter plot showing relative fold change by *Son* KD versus SON OE for both Tu-induced and Tu-repressed genes. (**c**) Top predicted transcription regulators of 461 and 901 genes by the LISA Cistrome DB TR ChIP-Seq models. (**d, e**) Selected genes aligned for SC35 and XBP1s ChIP-seq signal from CT12 in XBP1*^Flox^*mice ^6^ (**d**) and ChIP-qPCR of XBP1s and SC35 on selected regions (indicated by red bars) (n=3∼4 biologically independent samples collected from two independent experiments) (**e**). Panel d taken from Figure 5d of ^20^. © The Authors, some rights reserved; exclusive licensee AAAS. Distributed under a Creative Commons Attribution License 4.0 (CC BY) https://creativecommons.org/licenses/by/4.0/. Data: Mean ± S.E.M. Statistical tests used: unpaired two-tailed Student’s t-test for a. Linear regression for b. Three-way ANOVA and Tukey’s multiple comparisons test for e.

**Supplementary Figure 3.**
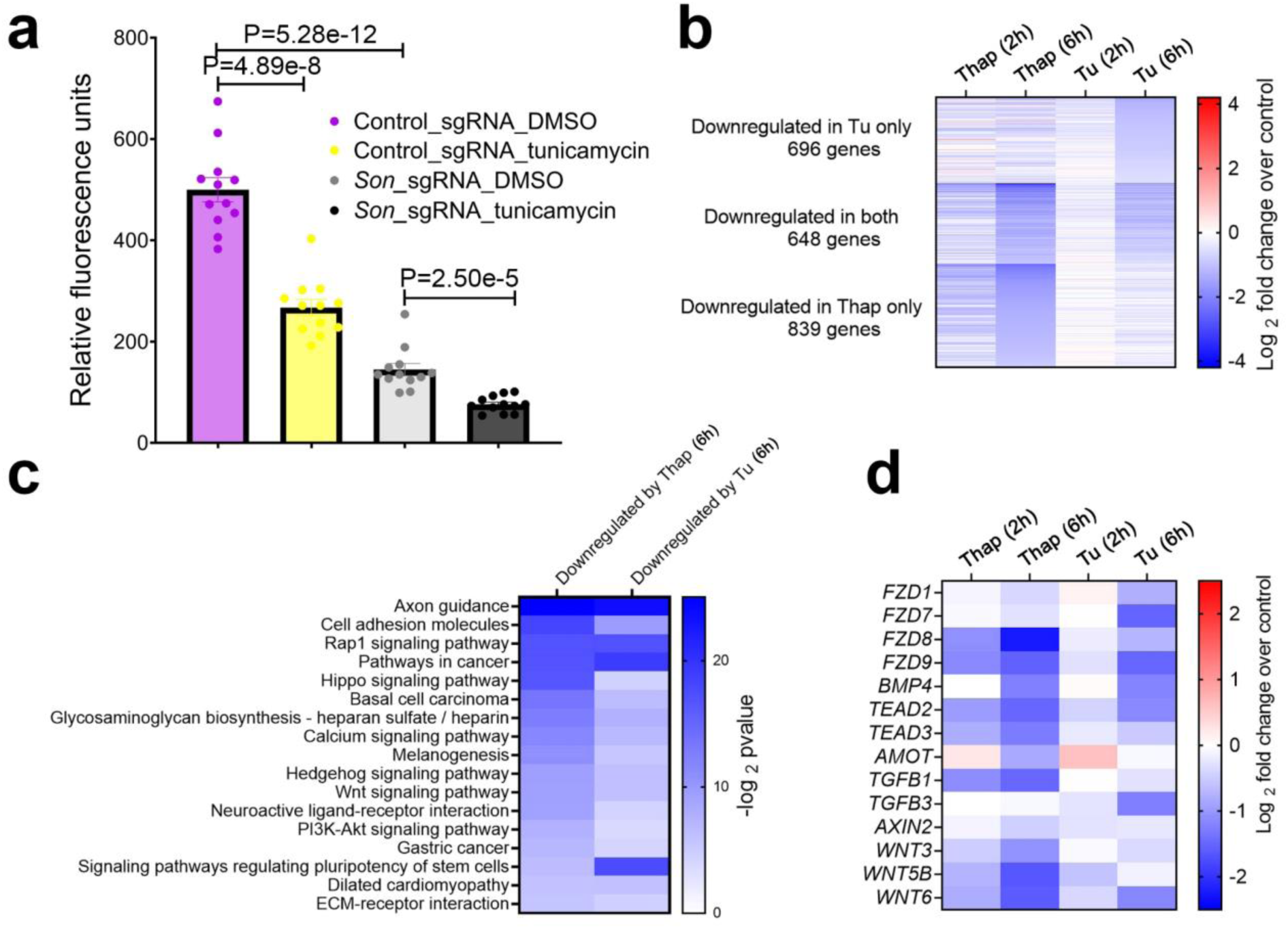
YAP1 transcriptional output is repressed during ER stress. (**a**) TEAD luciferase reporter assay in control and SON OE MEFs in response to 100ng/ml Tu for 6h (n=12 biologically independent samples for all groups). (**b**) Heatmap showing transcriptomes that are significantly downregulated (log _2_-fold change smaller than −0.5 with a p value less than 0.05) either under Tu or Thap treatment at 6h. (**c**) GO analysis of genes that are significantly downregulated either under Tu or Thap treatment at 6h. (**d**) Heatmap of representative YAP1 target genes as in **b**. Data: Mean ± S.E.M. Statistical tests used: unpaired two-tailed Student’s t-test for a.

**Supplementary Figure 4.**
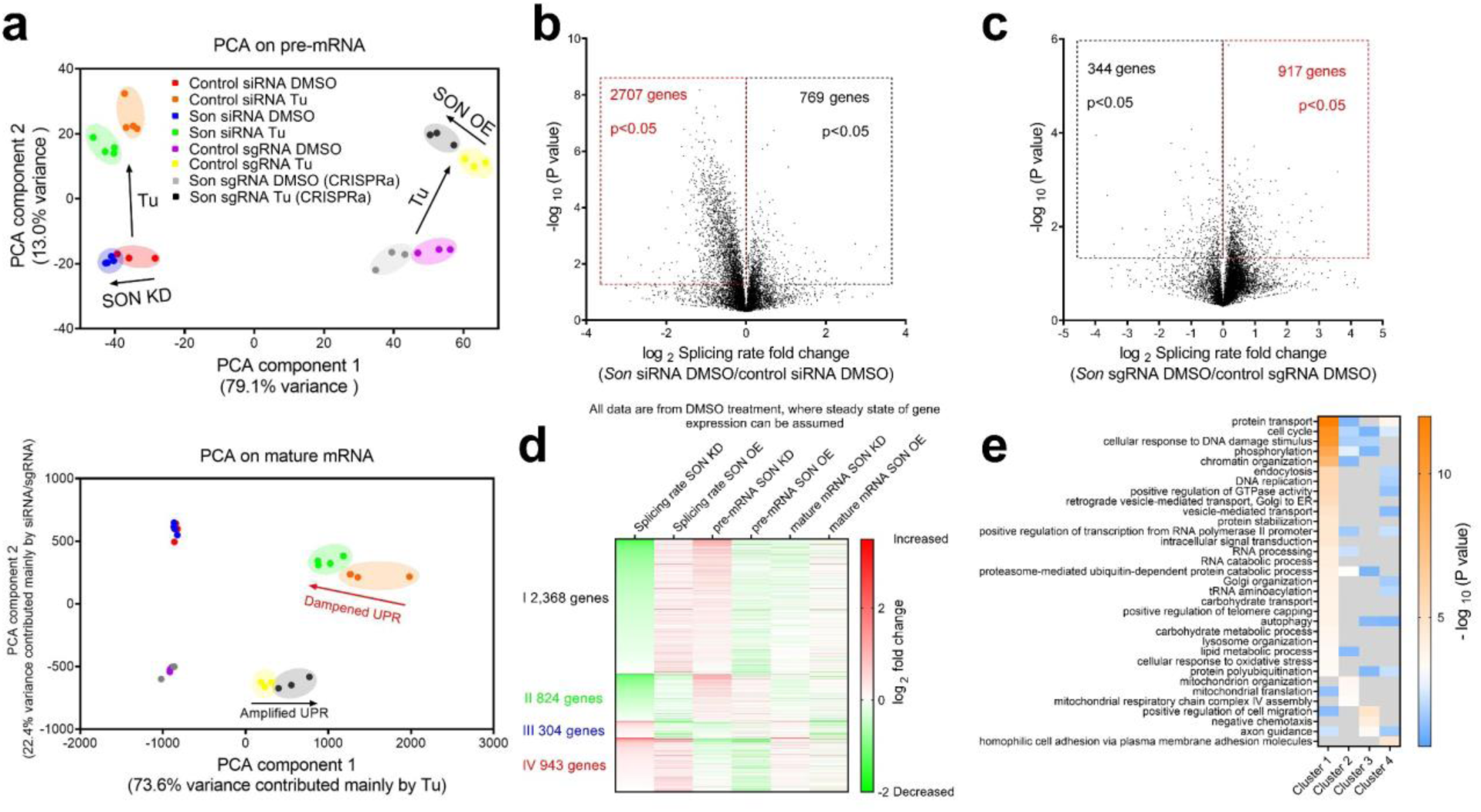
SON regulates mRNA splicing rates under both basal and ER stress conditions (**a**) PCA analysis of global transcriptional response to Tu in the presence of SON OE/KD. Both pre-mRNA (top) and mature mRNA (bottom) are shown. (**b, c**) Volcano plot of mRNA splicing rates changes in SON KD (**b**) or OE (**c**) MEFs under basal condition (DMSO). (**d**) Heat map of fold change of RNA splicing rate, pre and mature mRNA level in SON OE/KD MEFs compared to control MEFs under vehicle (DMSO) condition. Four clusters of genes are shown. (**e**) GO analysis of genes in four clusters showing enriched KEGG pathways.

**Supplementary Figure 5.**
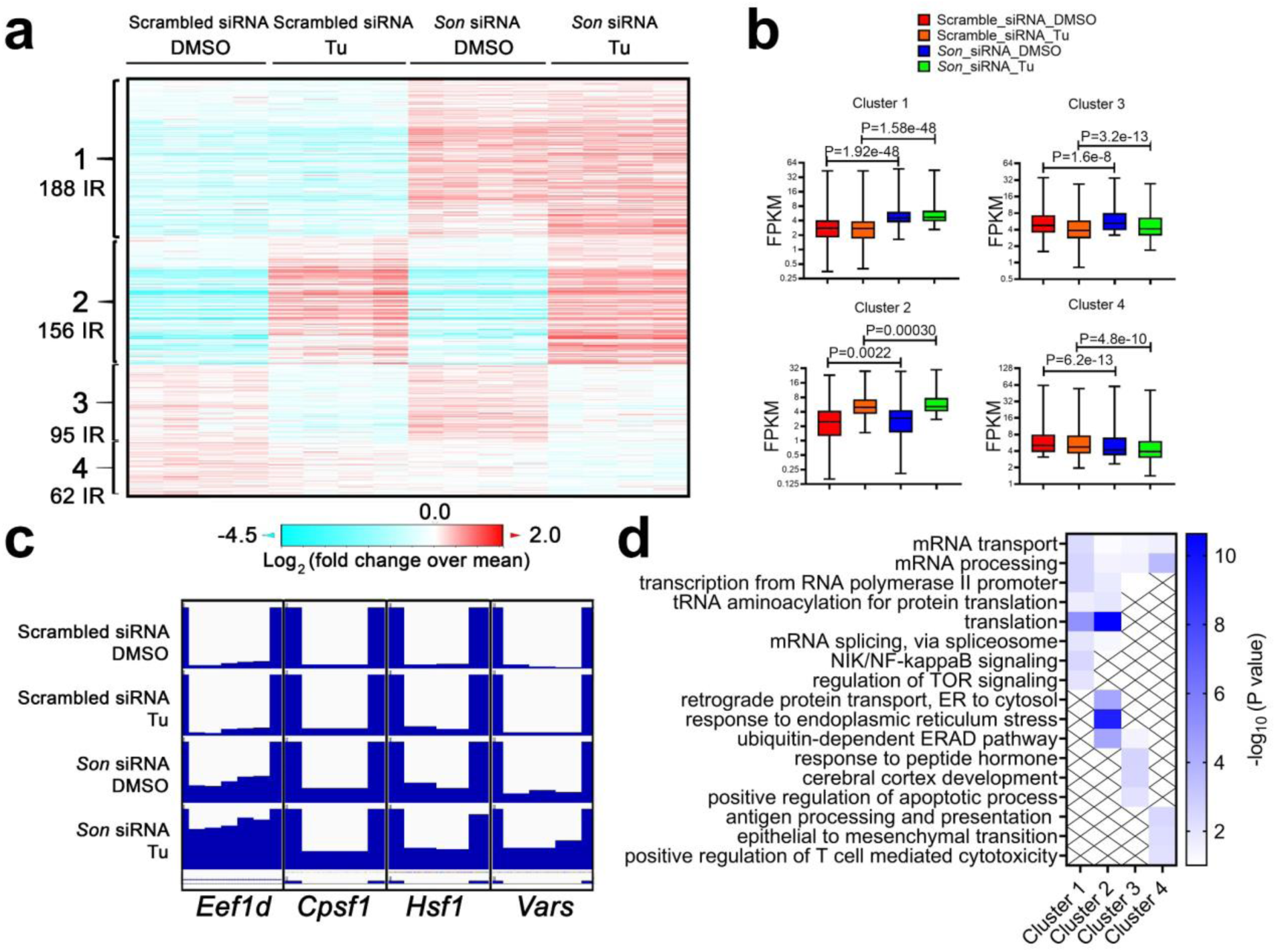
SON knockdown increases intron retention of proteostasis and mRNA metabolism genes. Heat map (**a**) and quantification (**b**) of intron retention events in MEFs with control or SON KD under basal (DMSO) and Tu conditions. Four clusters are shown. (**c**) The Integrative Genome Viewer representation of intron retention in selected genes. (**d**) GO analysis of genes in four clusters showing enriched KEGG pathways. Data: box and whiskers with minimum to maximum. Statistical tests used: unpaired two-tailed Student’s t-test for b.

**Supplementary Figure 6.**
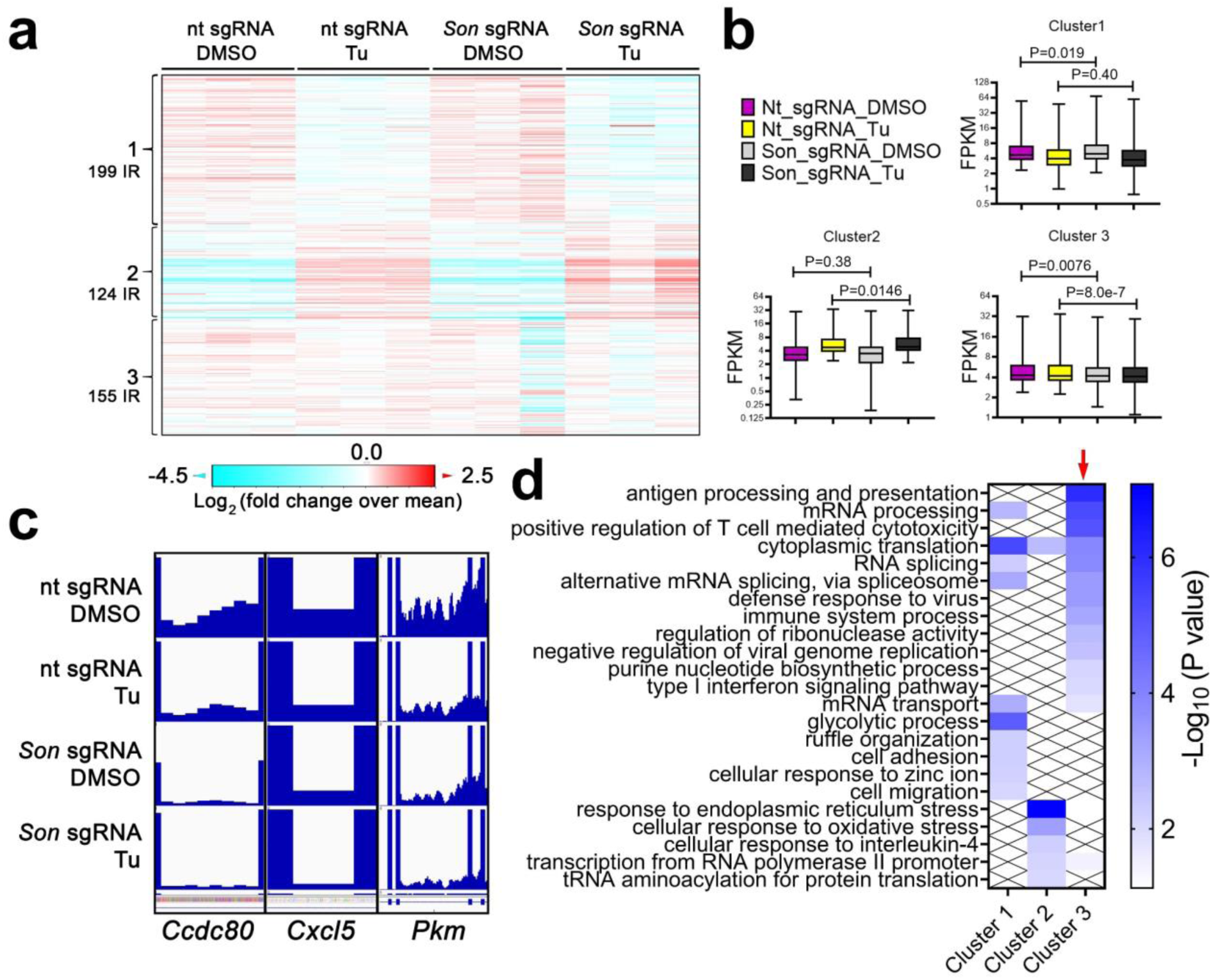
SON overexpression decreases intron retention of protein processing and mRNA metabolism genes. Heat map (**a**) and quantification (**b**) of intron retention events in MEFs with control or SON overexpression under basal (DMSO) and Tu conditions. Three clusters are shown. (**c**) The Integrative Genome Viewer representation of intron retention in selected genes. (**d**) GO analysis of genes in three clusters showing enriched KEGG pathways. Data: box and whiskers with minimum to maximum. Statistical tests used: unpaired two-tailed Student’s t-test for b.

**Supplementary Figure 7.**
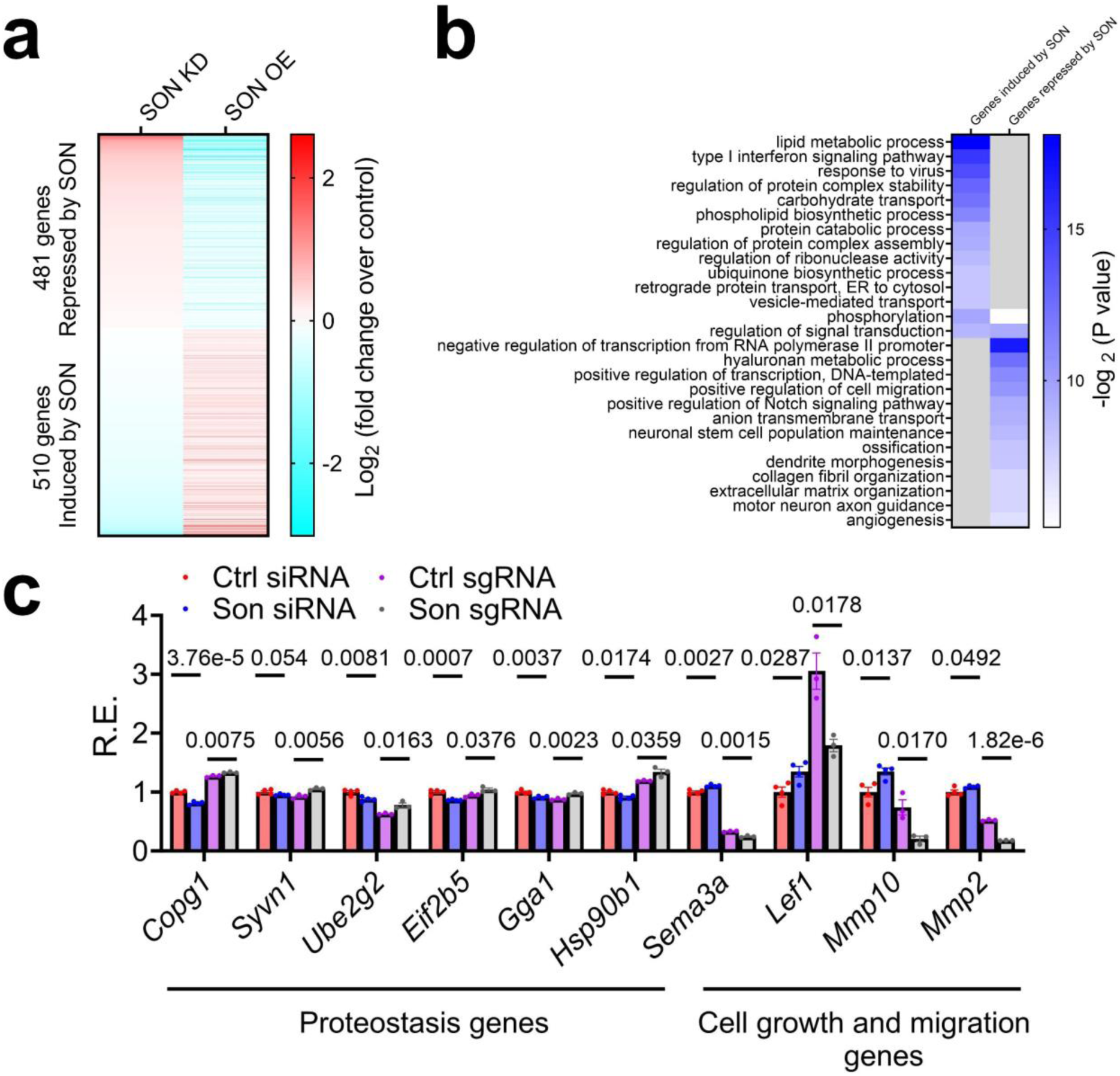
SON reprograms opposing proteostasis and YAP1 transcriptional output under basal conditions. (**a**) Heat map of fold change of mature mRNA level in *Son* OE/KD MEFs compared to control MEFs under vehicle (DMSO) condition. All mature mRNAs in this heatmap are statistically differentially expressed (P<0.05) in both SON OE/KD conditions, compared to their respective controls. (**b**) GO analysis of these 481 and 501 genes. (**c**) Representative mature mRNA expression of proteostasis and YAP1 target genes (n=4 biologically independent samples for Ctrl and *Son* siRNA and n=3 biologically independent samples for Ctrl and *Son* sgRNA). Data: Mean ± S.E.M. Statistical tests used: unpaired two-tailed Student’s t-test for c.

**Supplementary Figure 8.**
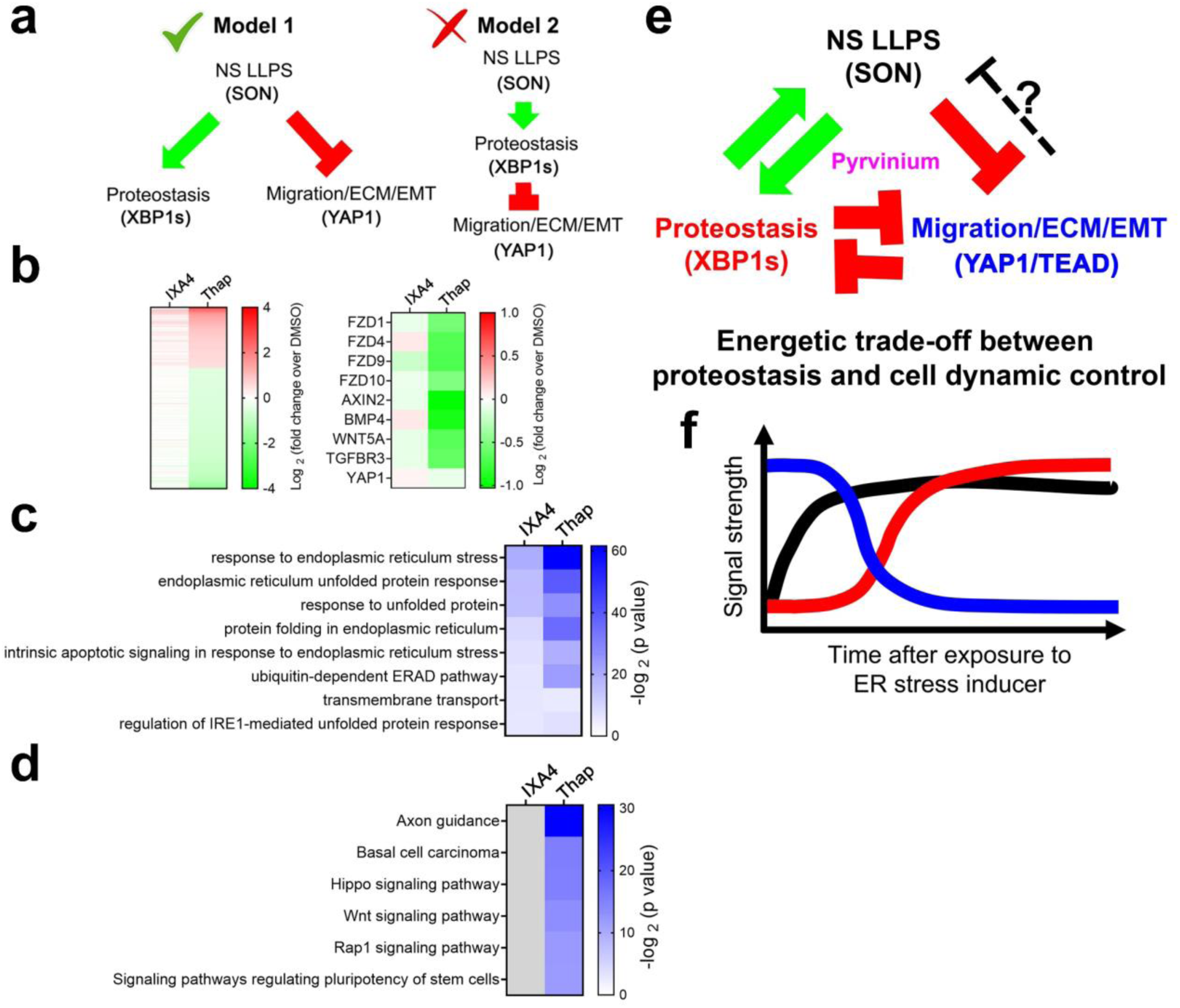
Nuclear speckle LLPS dictates opposing proteostasis and YAP1 signaling. (**a**) Two models explaining the relationship among nuclear speckles LLPS dynamics, proteostasis and YAP1 transcriptional output. Our results support model 1. (**b**) Heatmap demonstrates relative fold change of gene expression relative to DMSO control in IXA4, or Thap treated HEK293T cells. All genes induced or repressed by at least 1.41-fold with p value smaller than 0.05 in Thap condition (left), and representative YAP1-related genes (right). (**c, d**) GO analysis of all upregulated (**c**) or downregulated (**d**) genes in either IXA4 or Thap treatment by at least 1.41-fold with a p-value smaller than 0.05. (**e**) An expanded model of how the LLPS of nuclear speckles can dictate proteostasis and YAP1 transcriptional output. Please see the main text for details. (**f, g**) Diagram showing temporal changes of NS’ LLPS (black), proteostasis (red) and YAP1 transcriptional output (blue) signal during ER stress (**f**).

**Supplementary Figure 9.**
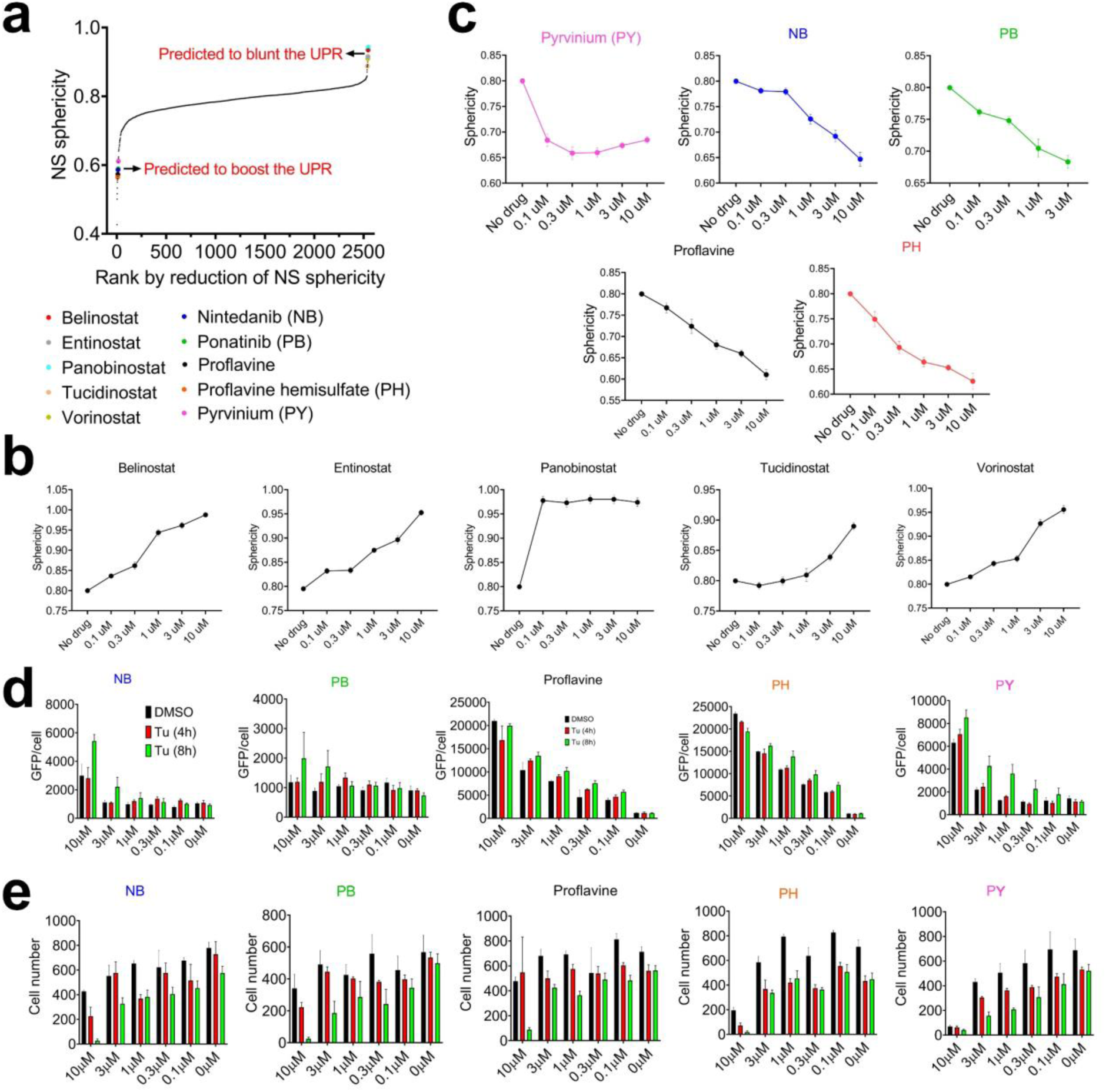
HTS identifies compounds that alter nuclear speckle morphology and the UPR. (**a**) Compounds in the FDA-approved library ranked from lowest to highest on their ability to reduce NS sphericity. (**b**) Five drugs are shown to have a dose-dependent effect on increasing sphericity of NS (n=16 biologically independent samples). (**c**) Dose-dependent effects of drugs on decreasing NS sphericity (n=16 biologically independent samples). (**d, e**) GFP/cell (**d**) or cell number (**e**) measured in *Perk* promoter-driven dGFP reporter MEFs in the presence of Tu for four or eight hours after pre-treatment of different concentrations of drugs or DMSO for 24 hours (n=4 biologically independent samples). Data: Mean ± S.E.M.

**Supplementary Figure 10.**
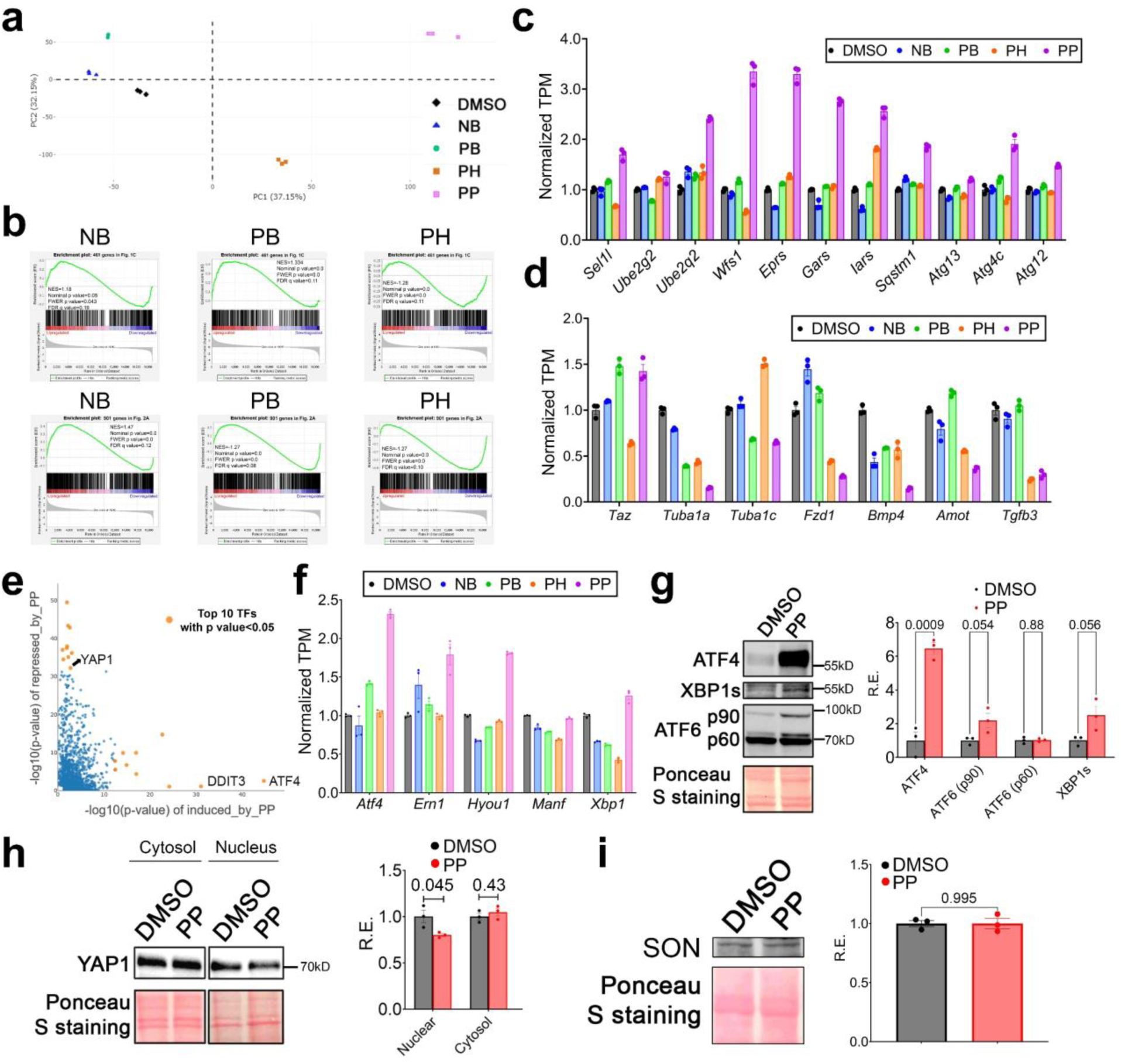
PP is a bona fide nuclear speckle rehabilitator. (**a**) MEFs were treated with 3 μM NB, 1 μM PB, 3 μM PH, and 1 μM PP for 24 hours and RNA-seq was performed. PCA of global transcriptional response to drug treatments. (**b**) For each of the GSEA analysis, genes further activated by SON OE or further repressed by SON OE are compared to the transcriptome signatures of MEFs under different drug treatments. (**c, d**) Gene expression of select protein quality control (**c**) and YAP1 target genes (**d**) determined through mRNA-Seq (n=3 biologically independent samples). (**e**) LISA analysis listing log transformed p values for top predicted transcription regulators for genes upregulated (x-axis) and downregulated (y-axis) by PP. (**f**) Gene expression of select UPR genes determined through mRNA-Seq under different drugs treatment (n=3 biologically independent samples). (**g-i**) Western blot and quantification of UPR TFs (**g**), YAP1 nuclear and cytosol (**h**) and SON (**i**) level in response to 1 μM PP for 24 hours (n=3 biologically independent samples for all data). Data: Mean ± S.E.M. Statistical tests used: unpaired two-tailed Student’s t-test for g-i. All Western blot raw blot images are provided in the source data file.

**Supplementary Figure 11.**
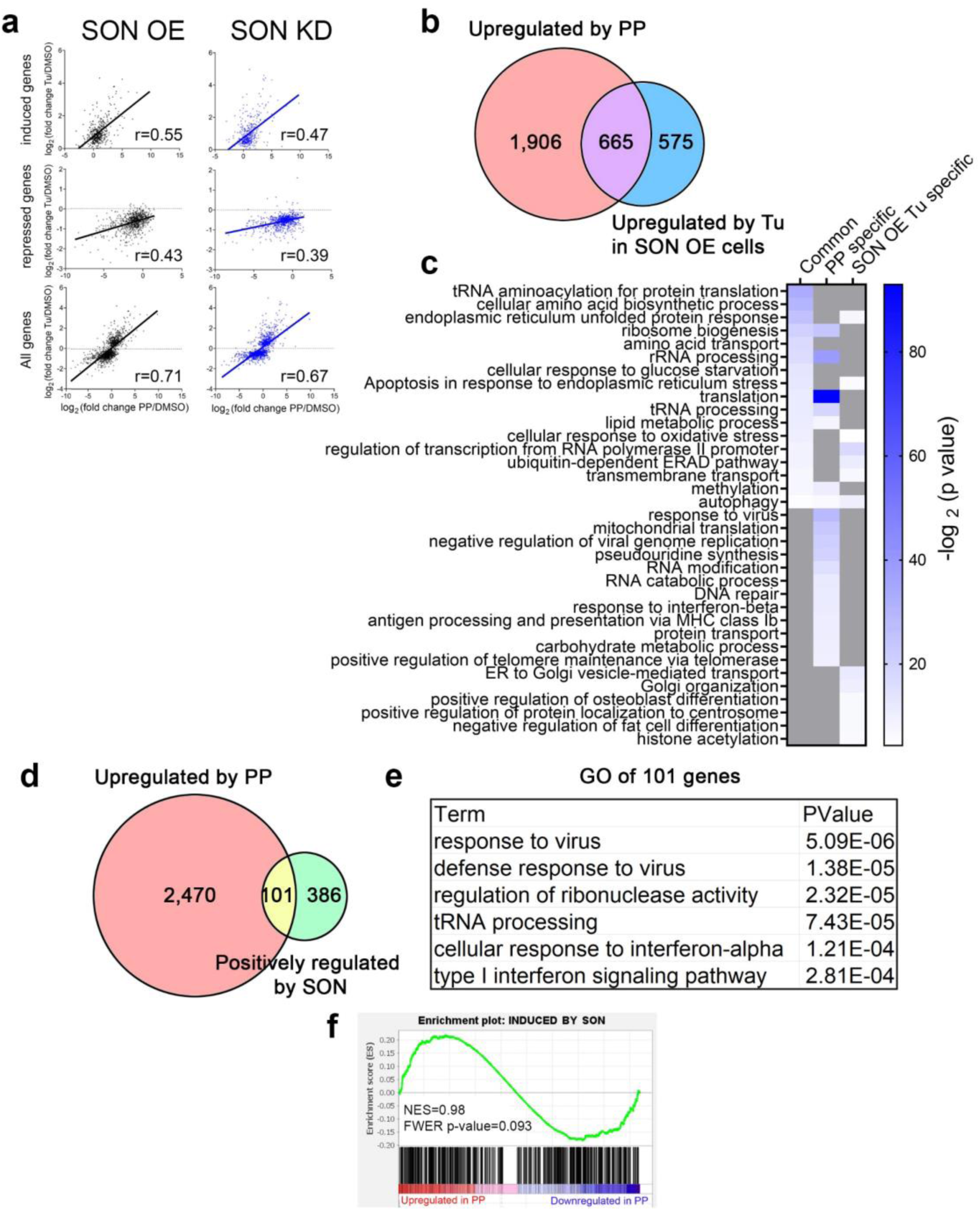
Comparison of upregulated genes by PP and SON OE. (**a**) Scatter plot comparing the fold change of gene expression by PP (x-axis) and by Tu (y-axis) under SON OE or SON KD condition. Correlation coefficient and p value are shown for each plot. Chow test indicates statistically significant coefficients between the two linear regressions with p=0.00195. (**b, c**) Venn diagram showing (**b**) and GO analysis of (**c**) specific and commonly upregulated genes by PP and Tu in SON OE MEFs. (**d**) Venn diagram showing specific and common upregulated genes by PP and SON in MEFs. (**e**) GO analysis of common 101 genes. (**f**) GSEA analysis comparing genes upregulated by SON with those regulated by PP.

**Supplementary Figure 12.**
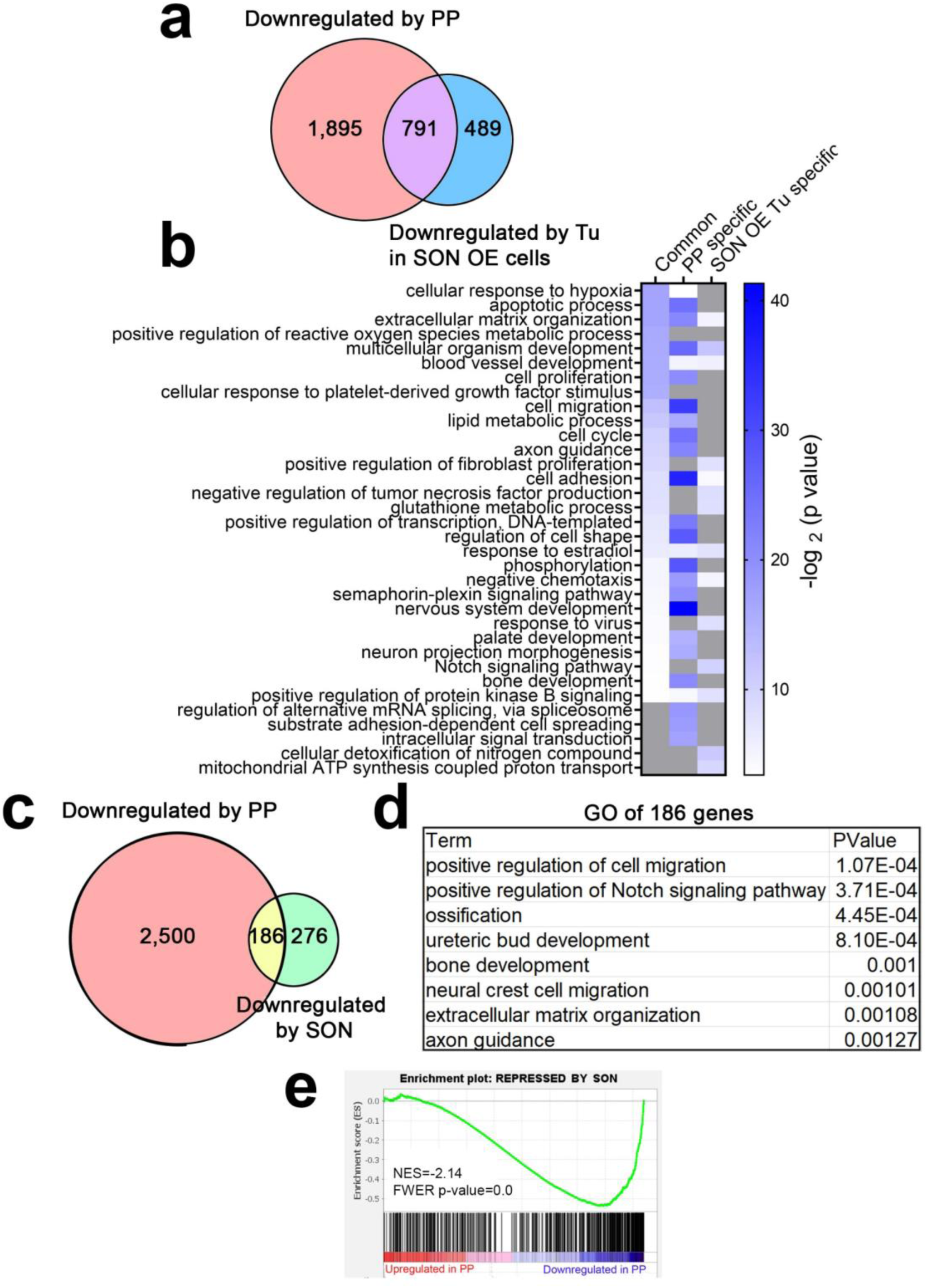
Comparison of downregulated genes by PP and SON OE. (**a-b**) Venn diagram showing (**a**) and GO analysis of (**b**) specific and common downregulated genes by PP and Tu in SON OE MEFs. (**c**) Venn diagram showing specific and common downregulated genes by PP and SON in MEFs. (**d**) GO analysis of 186 common genes. (**e**) GSEA analysis comparing genes downregulated by SON with those regulated by PP.

**Supplementary Figure 13.**
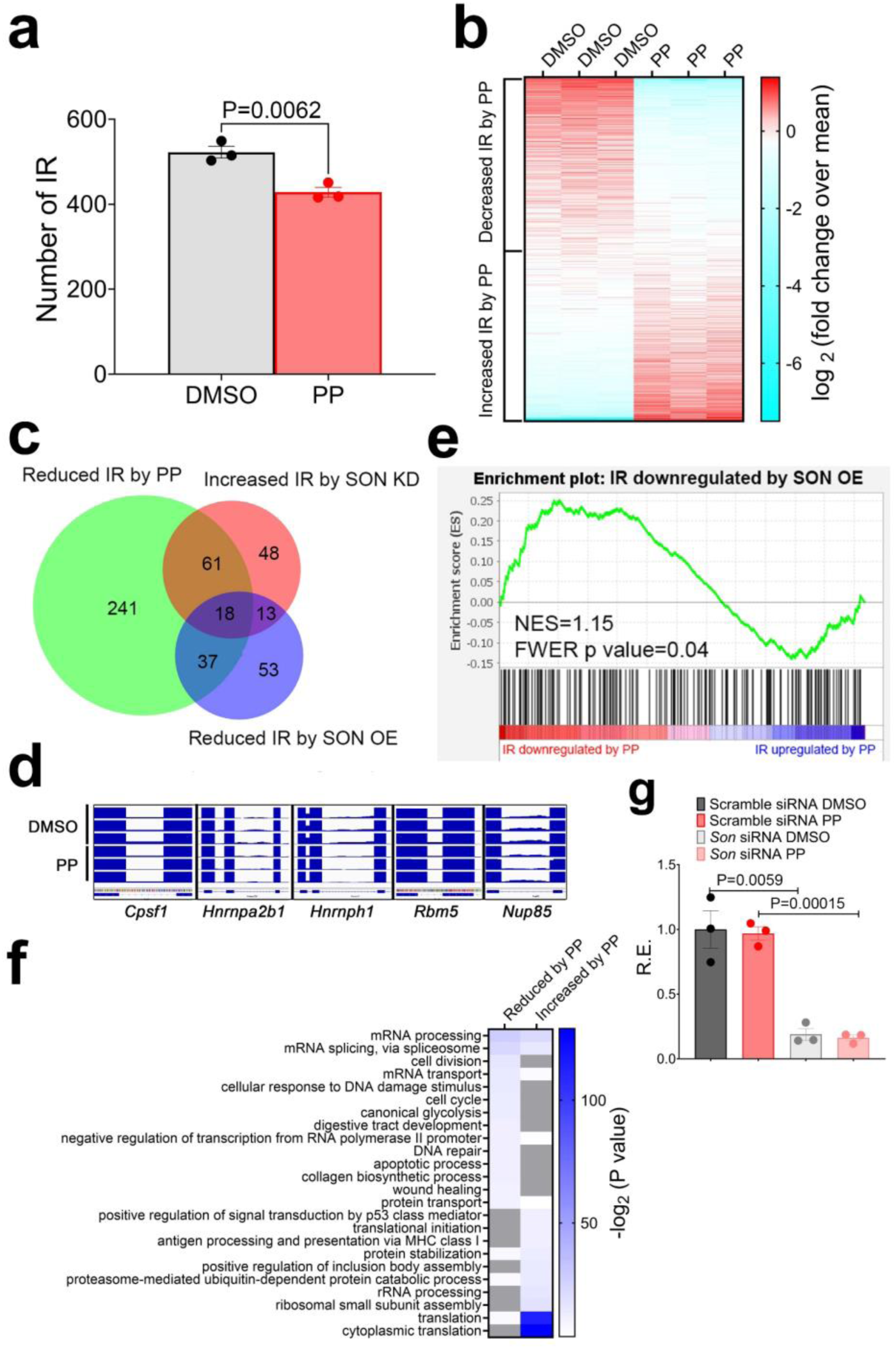
PP downregulates IR of genes involved in mRNA metabolism. (**a**) Quantification of IR under DMSO and PP condition (n=3 biologically independent samples for both treatment). (**b**) Heatmap showing RPKM normalized level of retained introns in DMSO and PP condition. (**c**) Venn diagram comparing genes with specific or common IRs between different conditions. (**d**) Genome browser view of selective genes with reduced IR by PP. (**e**) GSEA comparing IR downregulated by SON OE and PP. (**f**) GO analysis of genes with increased or reduced IR by PP. (**g**) qPCR of *Son* in MEFs transiently transfected scrambled or *Son* siRNA and treated with DMSO or 0.3µM PP for 24 hours (n=3 biologically independent samples). Data: Mean ± S.E.M. Statistical tests used: unpaired two-tailed Student’s t-test for a and g.

**Supplementary Figure 14.**
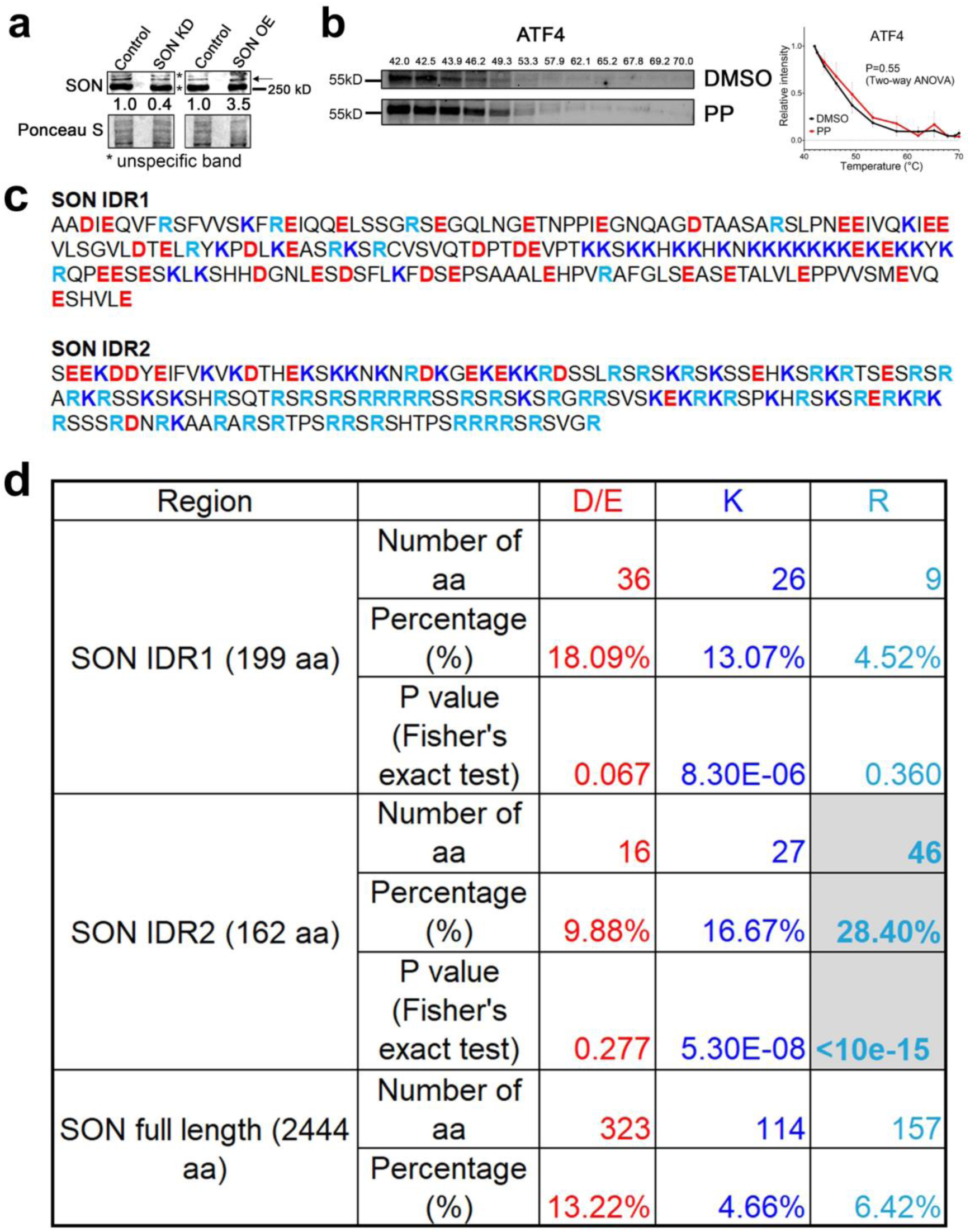
PP directly targets SON IDR2 to modulate nuclear speckle LLPS dynamics. (**a**) Western blots of SON with siRNA-mediated knockdown and SON OE in MEFs. (b) CETSA of ATF4 with 3µM PP. Both representative blot and quantification are shown (n=3 quantifications collected from three independent CETSAs for both DMSO and PP). (**c**) Sequence of SON IDR1 and IDR2. (**d**) The percentage and p values of enrichment (or lack thereof) of different amino acids. Data: Mean ± S.E.M. Statistical tests used: Two-way ANOVA for b. All Western blot raw blot images are provided in the source data file.

**Supplementary Figure 15.**
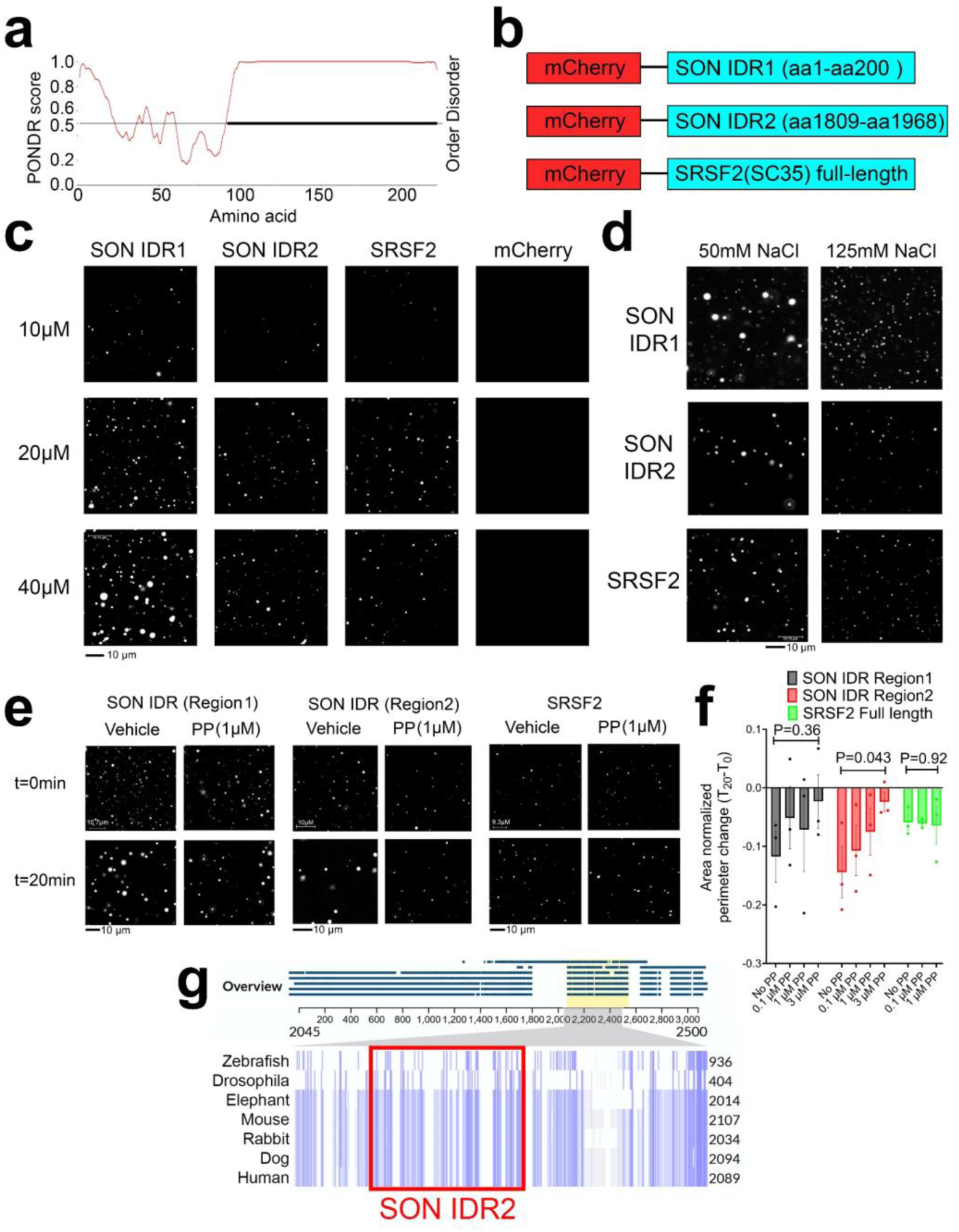
PP reduces the surface tension of nuclear speckle condensates. (**a**) Computational prediction of IDR in mouse SRSF2 (SC35) protein. (**b**) Diagram illustrating the constructs for droplet formation assay (**c**) Representative images of droplet formation assay with different concentrations of recombinant protein at 125mM NaCl. (**d**) Representative images of droplet formation assay with different salt concentrations with 20µM recombinant proteins. (**e, f**) Representative images of droplet formation assay with different recombinant proteins (**e**) and quantification (**f**) of area-normalized perimeter changes in the time span of 20 minutes with 50mM NaCl (n=3 quantifications collected from three independent droplet formation assays for each treatment). (**g**) Alignment of protein sequences of SON orthologs in seven different species. SON IDR2 is located within the most conserved region (highlighted by light yellow). Data: Mean ± S.E.M. Statistical tests used: One-way ANOVA for f.

**Supplementary Figure 16.**
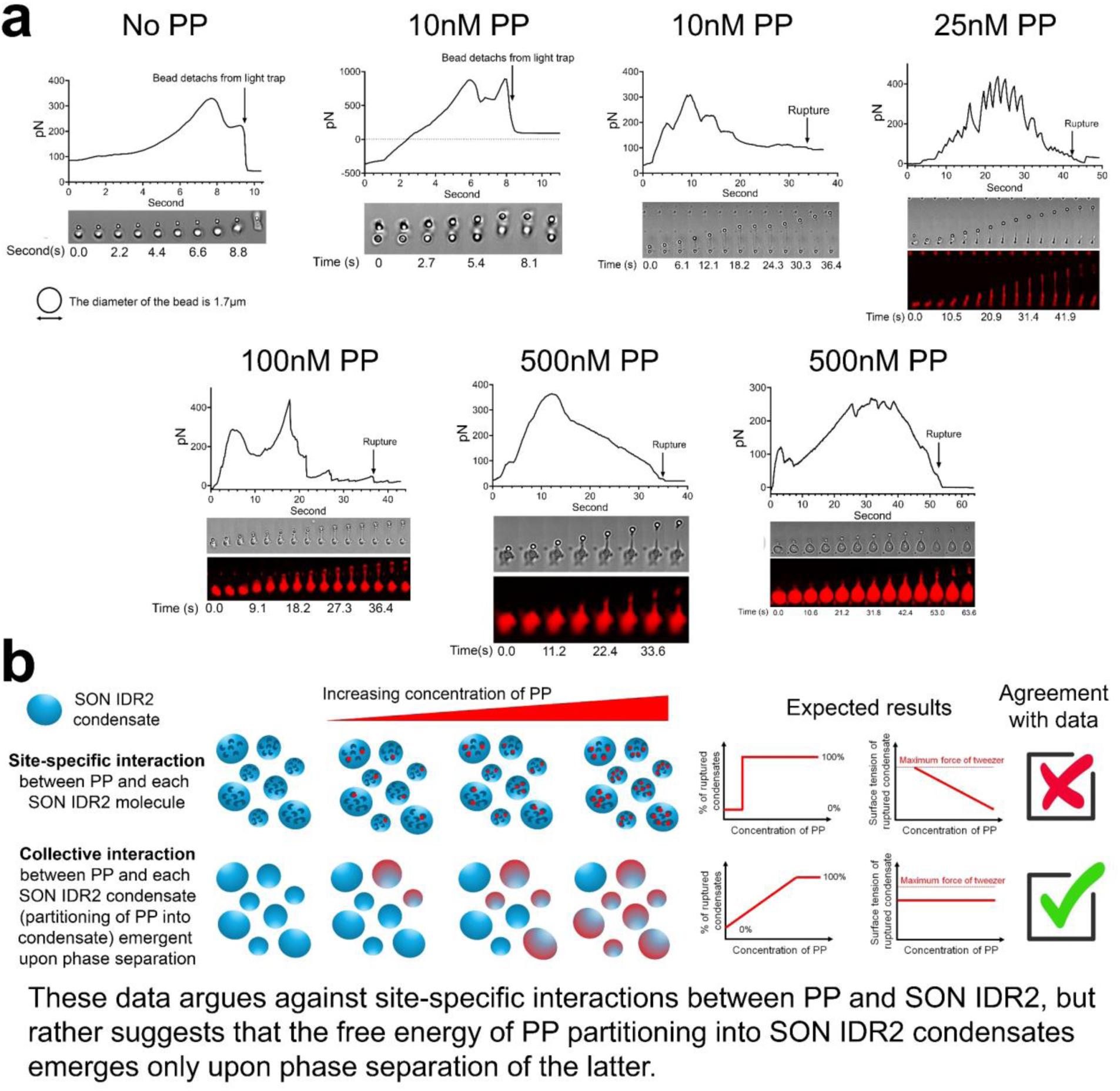
PP reduced the surface tension of SON IDR2 condensate. (**a**) Force recordings as well as time lapse fluorescent and bright field images of condensate stretching with different concentrations of PP. (**b**) A diagram illustrating two models of how PP interacts with SON IDR2, predicted results and whether the predicted results agree with experimental data.

**Supplementary Figure 17.**
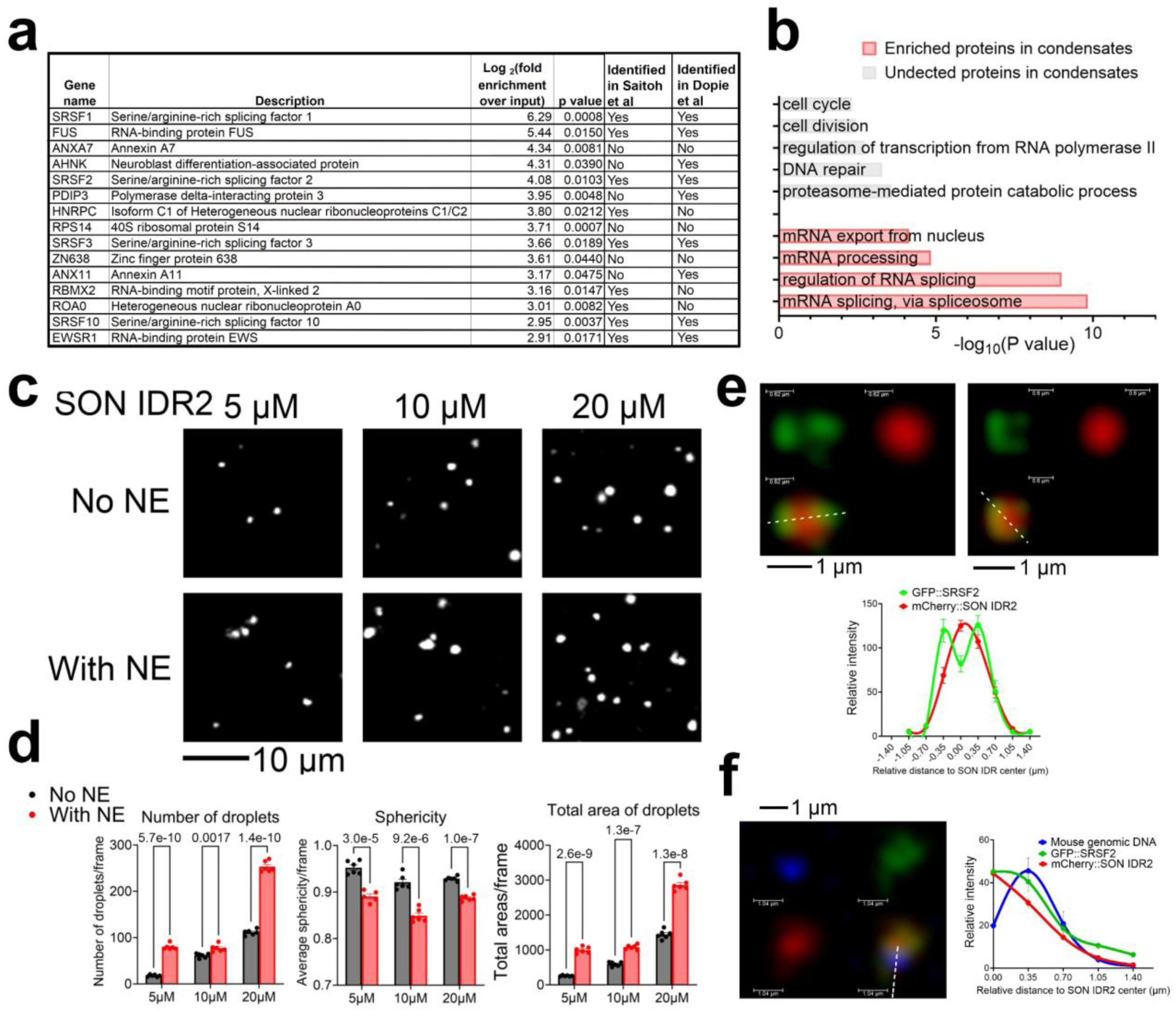
PP modulates nuclear speckle LLPS dynamics in a cell-free system. (**a**) HeLa NE-supplemented SON IDR2 condensates are spun down and subject to mass spectrometry. Top 15 proteins mostly enriched in SON IDR2-compartmentzlied condensates with p value<0.05, and the status of whether these proteins have been identified in nuclear speckles in cells in two datasets ^24, 25^ (**b**) GO of top enriched biological pathways of proteins depleted or enriched in SON IDR2-compartmentzlied condensates. (**c, d**) Representative images of droplet formation assay with increasing concentration of SON IDR2 with or without NE supplementation (c) and quantification (**d**) of the number, sphericity and total areas of droplets (n=6 quantifications from 6 image frames collected evenly across three independent droplet formation assays for each group). (**e**) Representative images and quantification of spatial distribution of mCherry::SON IDR2 and GFP::SRSF2 (n=16 nuclear speckles collected evenly across three independent droplet formation assays). (**f**) Representative images and quantification of spatial distribution of mCherry::SON IDR2, GFP::SRSF2 and mouse genomic DNA (n=43 nuclear speckles collected evenly across three independent droplet formation assays). Data: Mean ± S.E.M. Statistical tests used: unpaired two-tailed Student’s t-test for d.

**Supplementary Figure 18.**
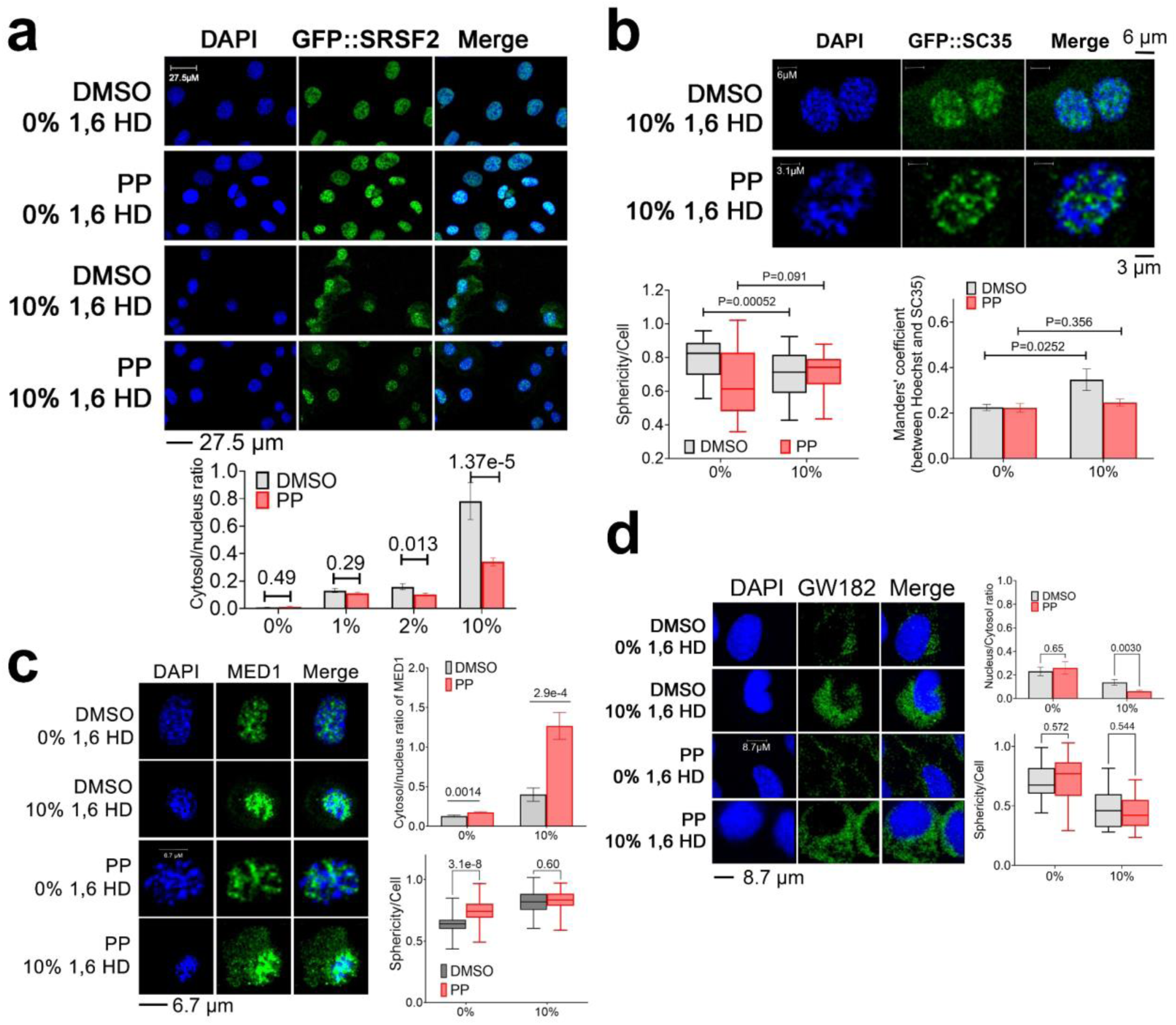
PP alters the sensitivity of different condensates to 1,6 hexanediol. (**a**) 1,6 hexanediol sensitivity assay with representative images and quantification (n=20∼95 cells collected evenly across four separate wells for each treatment) of the ratio of cytosol over nuclear intensity of GFP::SRSF2 signal. (**b**) 1,6 hexanediol sensitivity assay with representative images and quantification of sphericity (n=24∼50 cells collected evenly across four separate wells for each treatment) of GFP::SRSF2 signal and Manders’ coefficient (n=12∼14 cells collected evenly across four separate wells for each treatment) of signals of GFP and Hoechst. (**c**) 1,6 hexanediol sensitivity assay with representative images and quantification of ratio of cytosol to nuclear MED1 signal (n=19∼29 cells collected evenly across four separate wells for each treatment) and sphericity of MED1 signal (n=31∼75 cells collected evenly across four separate wells for each treatment). (**d**) 1,6 hexanediol sensitivity assay with representative images and quantification of ratio of cytosol to nuclear GW182 signal (n=16∼54 cells collected evenly across four separate wells for each treatment) and sphericity (n=19∼97 cells collected evenly across four separate wells for each treatment) of GW182 signal. All data was collected from at least two independent experiments. Data: Mean ± S.E.M. Statistical tests used: unpaired two-tailed Student’s t-test for all data.

**Supplementary Figure 19.**
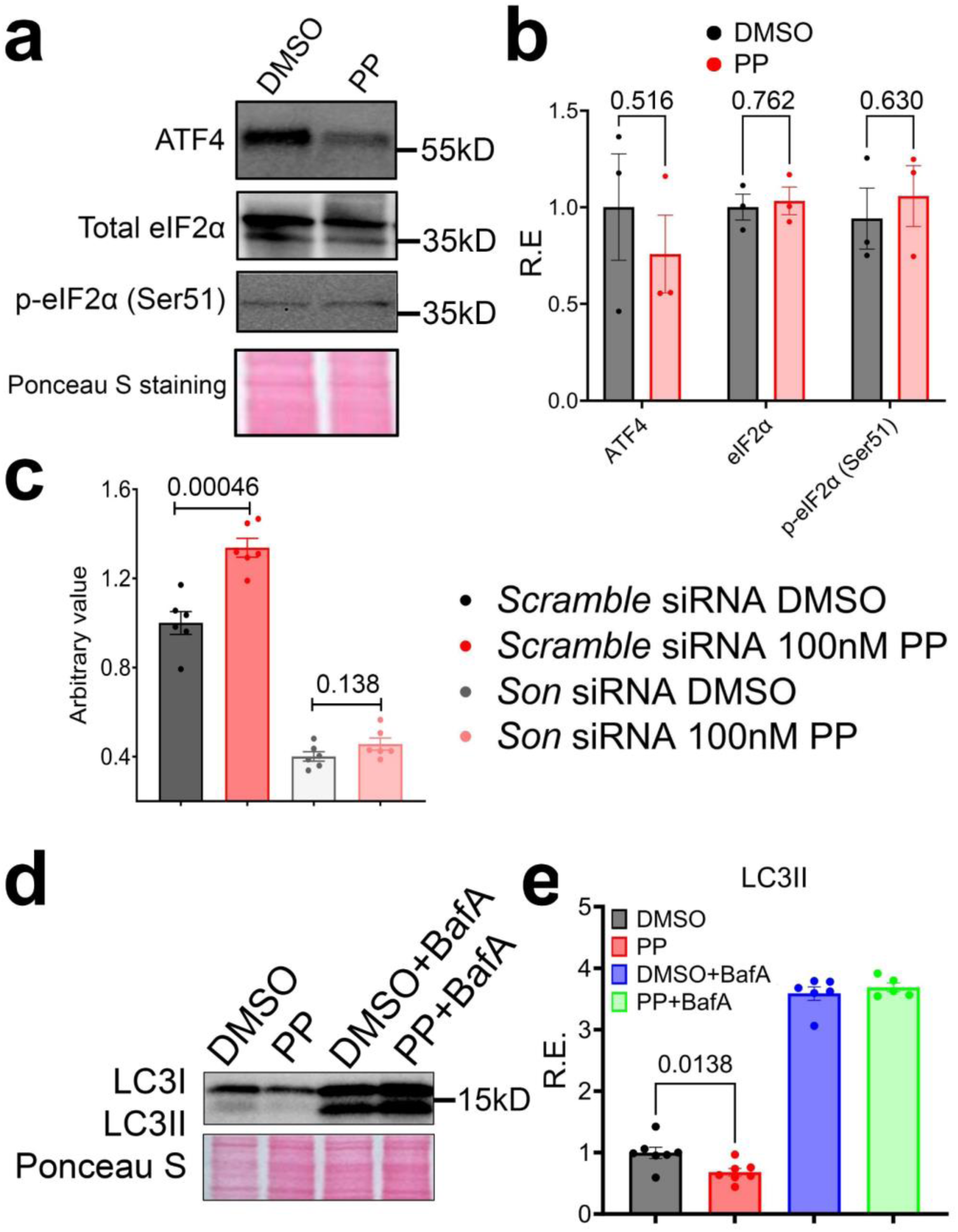
Nanomolar concentration of PP promotes UPS and ALP without inducing cellular stress. (**a, b**) MEFs were treated with DMSO or 100nM PP for 24 hours. Representative western blot (**a**) and quantification (**b**) of different proteins (n=3 biologically independent samples for both treatments). (**c**) MEFs were transiently transfected with scrambled or Son siRNA for 24 hours before treated with DMSO or 100nM PP for another 24 hours. Proteasome activity assay was then performed (n=6 biologically independent samples for each treatment collected from two independent experiments). (**d, e**) MEFs were treated with vehicle control or 0.1µM PP for ∼22 hours and then co-treated with or without Baf A (100nM for 22 hours) (n=5∼7 biologically independent samples for each treatment collected from three independent experiments). Representative western blot image (**d**) and quantification (**e**) of LC3II and LC3II/LC3I ratio. Data: Mean ± S.E.M. Statistical tests used: unpaired two-tailed Student’s t-test for all data. All Western blot raw blot images are provided in the source data file.

**Supplementary Figure 20.**
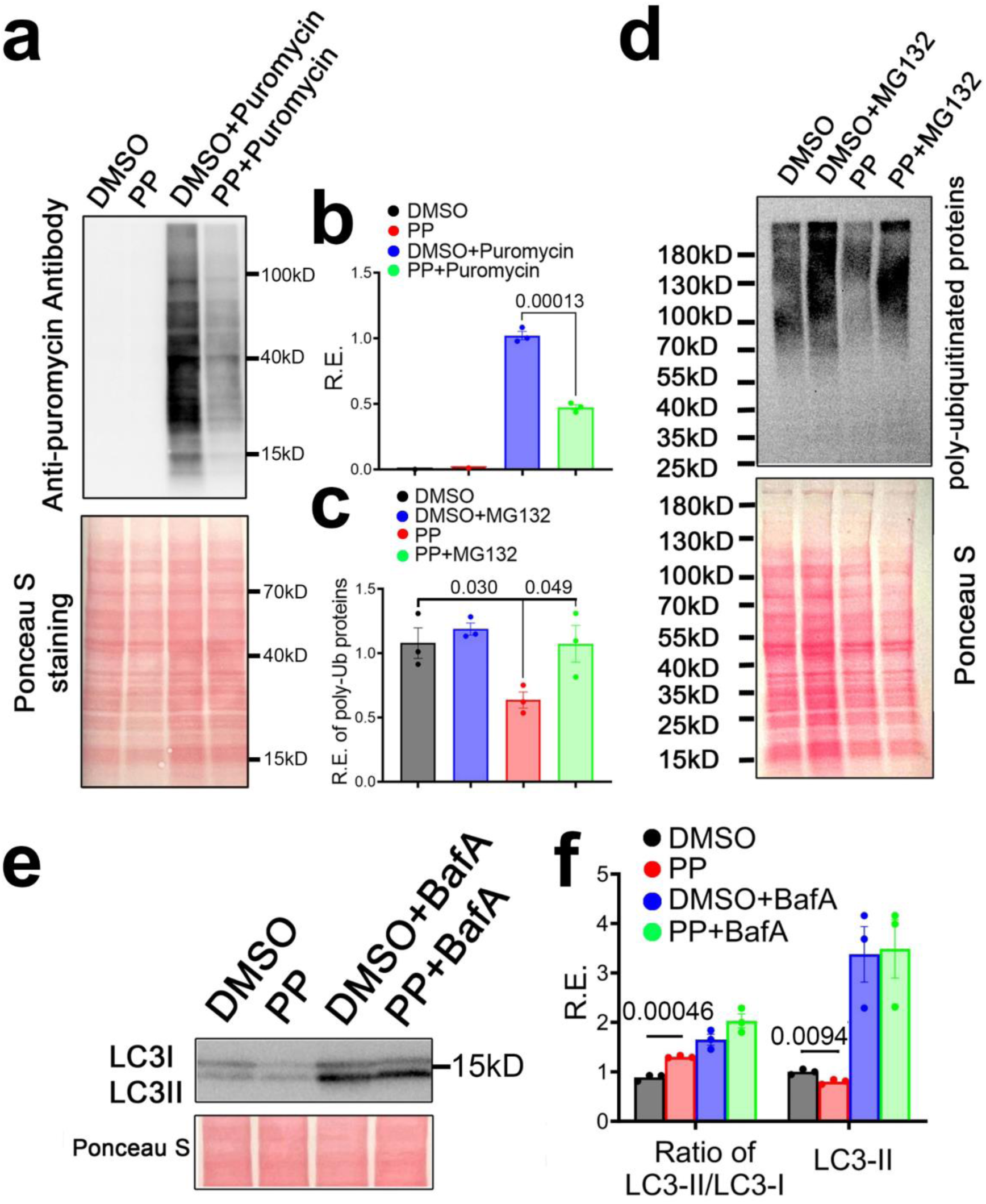
Micromolar PP promotes autophagy and UPS activity and represses translation. MEFs were treated with vehicle control or 1µM PP for ∼24 hours (22 hours for **e** and **f**) and then co-treated with or without puromycin (10 μg/mL for 30 minutes), MG132 (10μM for 110 minutes) or Baf A (100nM for 22 hours) (n=3 biologically independent samples for each treatment, collected from two independent experiments). Western blot and quantification of puromycin-incorporated proteins (**a, b**), poly-ubiquitinated protein (**c, d**) and LC3II and LC3II/LC3I ratio (**e, f**). Data: Mean ± S.E.M. Statistical tests used: unpaired two-tailed Student’s t-test for all data. All Western blot raw blot images are provided in the source data file.

**Supplementary Figure 21.**
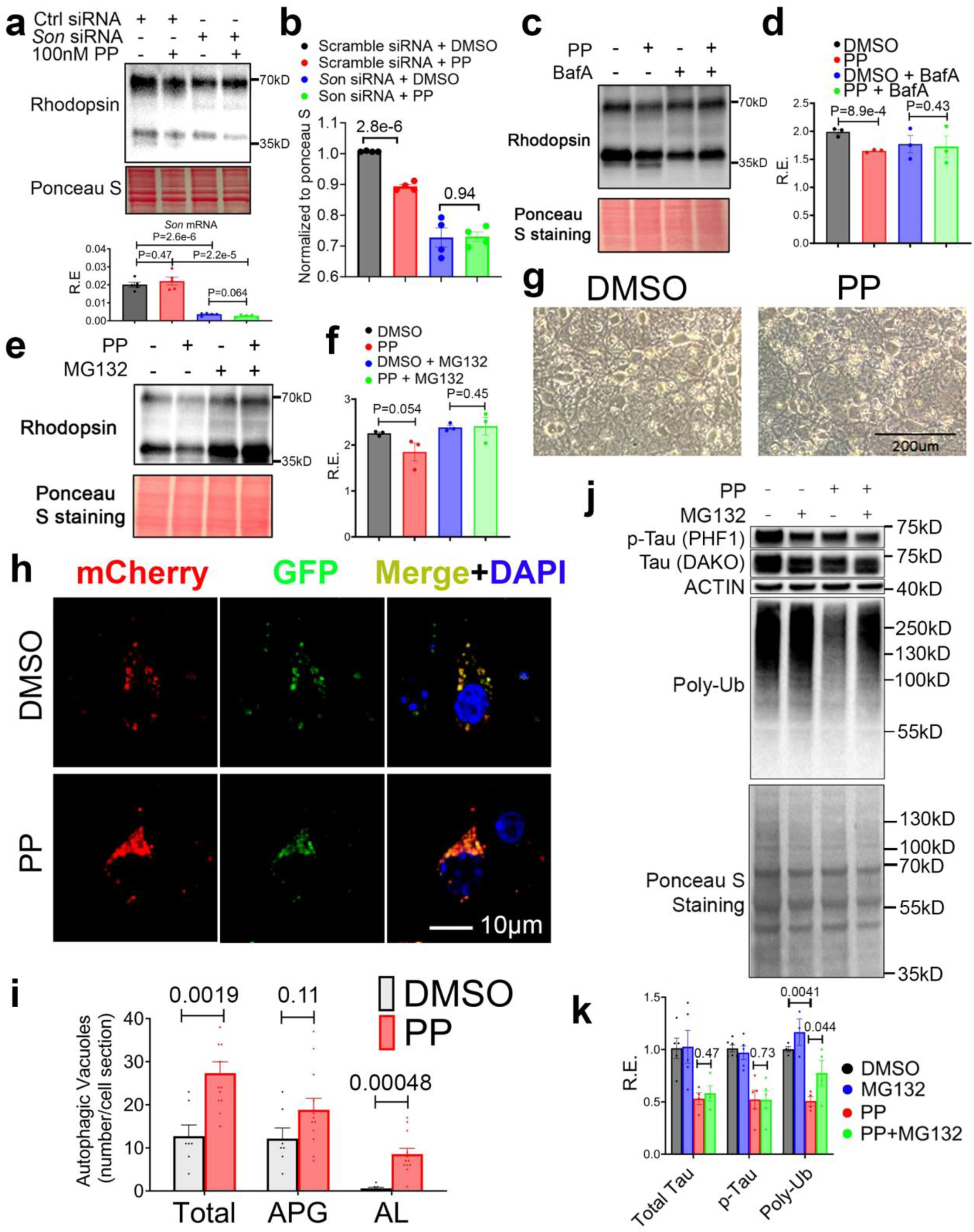
Pyrvinium pamoate reduces pathological Tau and Rhodopsin level by boosting autophagy and UPS activity. (**a, b**) NIH3T3 RHO^P23H^ cells were transfected with scrambled or *Son* siRNA for 24 hours before treated with DMSO or 0.1µM PP for another 24 hours. Western blot and qPCR of *Son* expression (n=5 biologically independent samples for each treatment collected from two independent experiments) (**a**) and quantification (**b**) of RHO^P23H^ level (n=4 biologically independent samples for each treatment). (**c, d**) NIH3T3 RHO^P23H^ cells were treated with 0.1µM PP and co-treated with or without BafA (100nM) for 24 hours. Western blot (**c**) and quantification (**d**) of RHO^P23H^ level (n=3 biologically independent samples for each treatment). (**e, f**) NIH3T3 RHO^P23H^ cells were co-treated with or without MG132 (10μM for 120 minutes). Western blot (**e**) and quantification (**f**) of RHO^P23H^ level (n=3 biologically independent samples for each treatment). (**g**) Representative images of primary mouse neurons treated with DMSO or 0.5 µM PP for 24 hours. (**h, i**) Representative images showing an increase of the number of mCherry positive puncta in primary neurons cultured in the presence of 0.1µM PP for 12 hours (**h**). Quantification of the number of total vacuoles, autophagosome and autolysosomes (n=7 and 12 cells for DMSO and PP, respectively, collected evenly across four separate wells for each treatment from two independent experiments) (**i**). (**j, k**) Tau P301S-expressing primary neurons were co-treated with vehicle or 0.1µM PP in the presence or absence of MG132 (10µM) for 12 hours and western blot (**j**) and quantification (**k**) of different proteins (n=3∼6 biologically independent samples for each treatment, collected from three independent experiments). All data mean ± S.E.M. Statistical tests used: unpaired two-tailed Student’s t-test for b, and i. Paired two-tailed Student’s t-test for k. Unpaired one-tailed Student’s t-tests for d and f, which are justified due to strong previous evidence in Fig. 5g-j demonstrating unidirectional change of Rhodopsin by PP treatment in NIH3T3 RHO^P23H^ cells (seven biological repeats). All Western blot raw blot images are provided in the source data file.

**Supplementary Figure 22.**
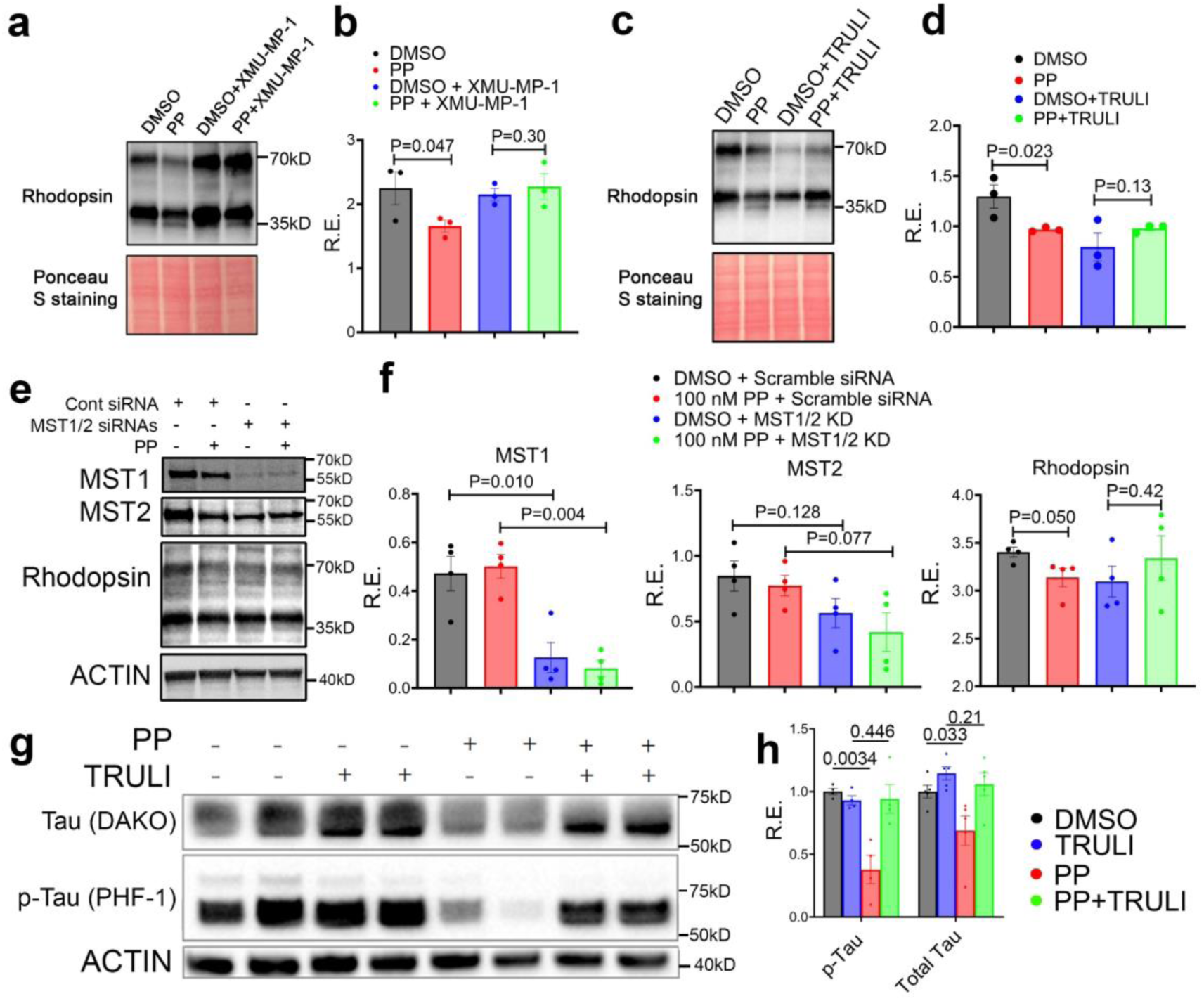
Pyrvinium pamoate reduces pathological Rhodopsin level in a manner that depends on reduced YAP1 activity. (**a, b**) NIH3T3 RHO^P23H^ cells were treated with 0.1µM PP for 24 hours and co-treated with or without XMU-MP-1 (1µM). Western blot (**a**) and quantification (**b**) of RHO^P23H^ level (n=3 biologically independent samples for each treatment). (**c, d**) NIH3T3 RHO^P23H^ cells were treated with 0.1µM PP for 24 hours and co-treated with or without TRULI (1µM). Western blot (**c**) and quantification (**d**) of RHO^P23H^ level (n=3 biologically independent samples for each treatment). (**e, f**) NIH3T3 RHO^P23H^ cells were transiently transfected with scrambled or Mst1/Mst2 siRNAs for 24 hours and then treated with DMSO or 0.1µM PP for another 24 hours. Western blot (**e**) and quantification (**f**) of MST1/2 and RHO^P23H^ level (n=4 biologically independent samples for each treatment). (**g, h**) Tau P301S-expressing primary neurons were co-treated with vehicle or 0.1µM PP in the presence or absence of YAP1 activator TRULI (10µM) for 12 hours and western blot (**g**) and quantification (**h**) of different proteins (n=4∼5 biologically independent samples for each treatment, collected from three independent experiments). All data: mean ± S.E.M. Statistical tests used: unpaired two-tailed Student’s t-test for f. Unpaired one-tailed Student’s t-tests for b and d, which are justified due to strong previous evidence in Fig. 5g-j demonstrating unidirectional change of Rhodopsin by PP treatment in NIH3T3 RHO^P23H^ cells (7 biological replicates). Paired one-tailed Student’s t-tests for h, which is justified due to strong previous evidence in Fig. 5k-m demonstrating unidirectional change of both total and phosphor-Tau by PP treatment in Tau P301S-expressing primary neurons (8∼12 biological replicates). All Western blot raw blot images are provided in the source data file.

**Supplementary Figure 23.**
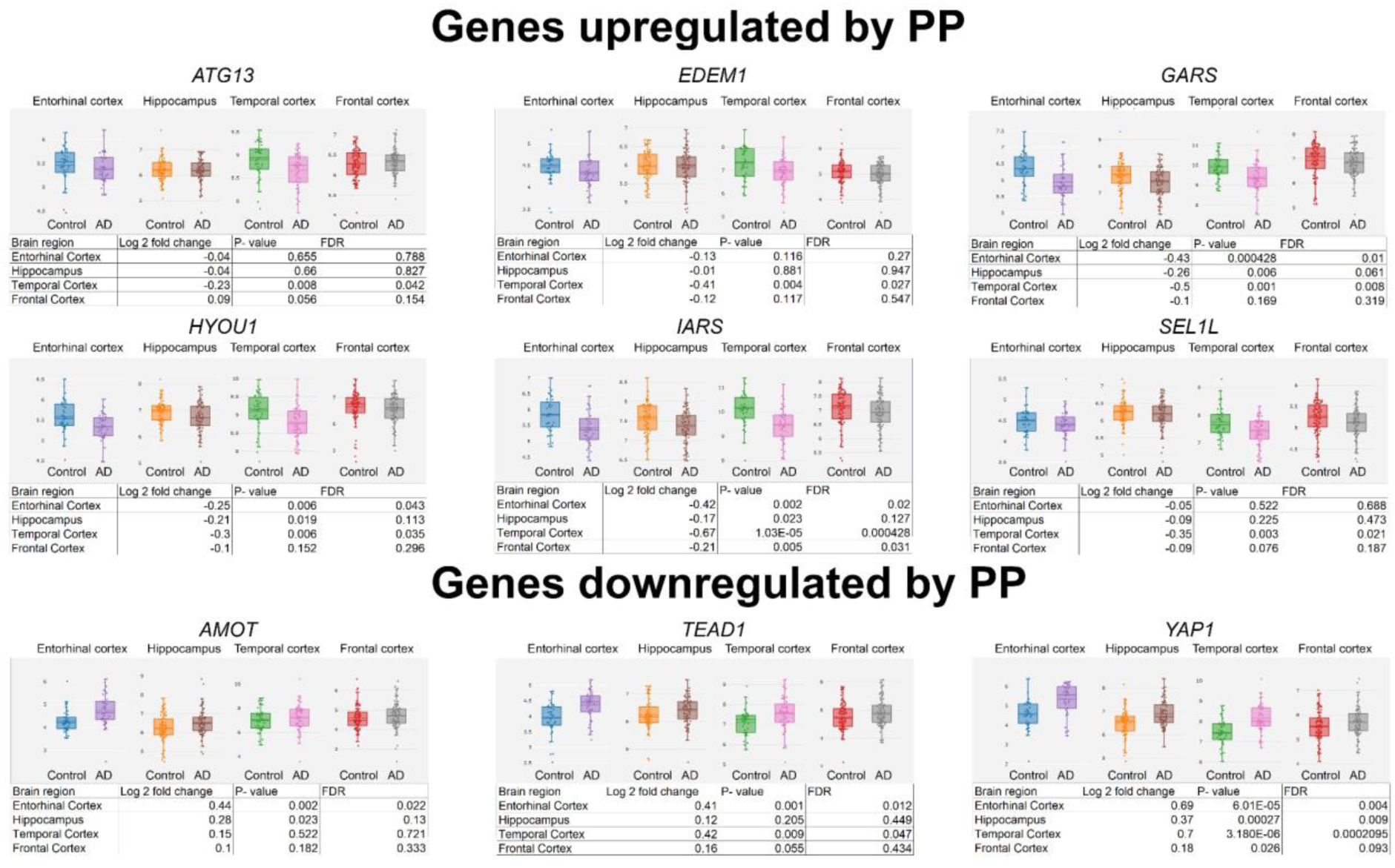
Gene signatures in human AD subjects are opposite from those regulated by PP revealed by bulk RNA-Seq. Relative gene expressions in different brain regions of human AD subjects normalized to control subjects as reported in ^86^.

**Supplementary Figure 24.**
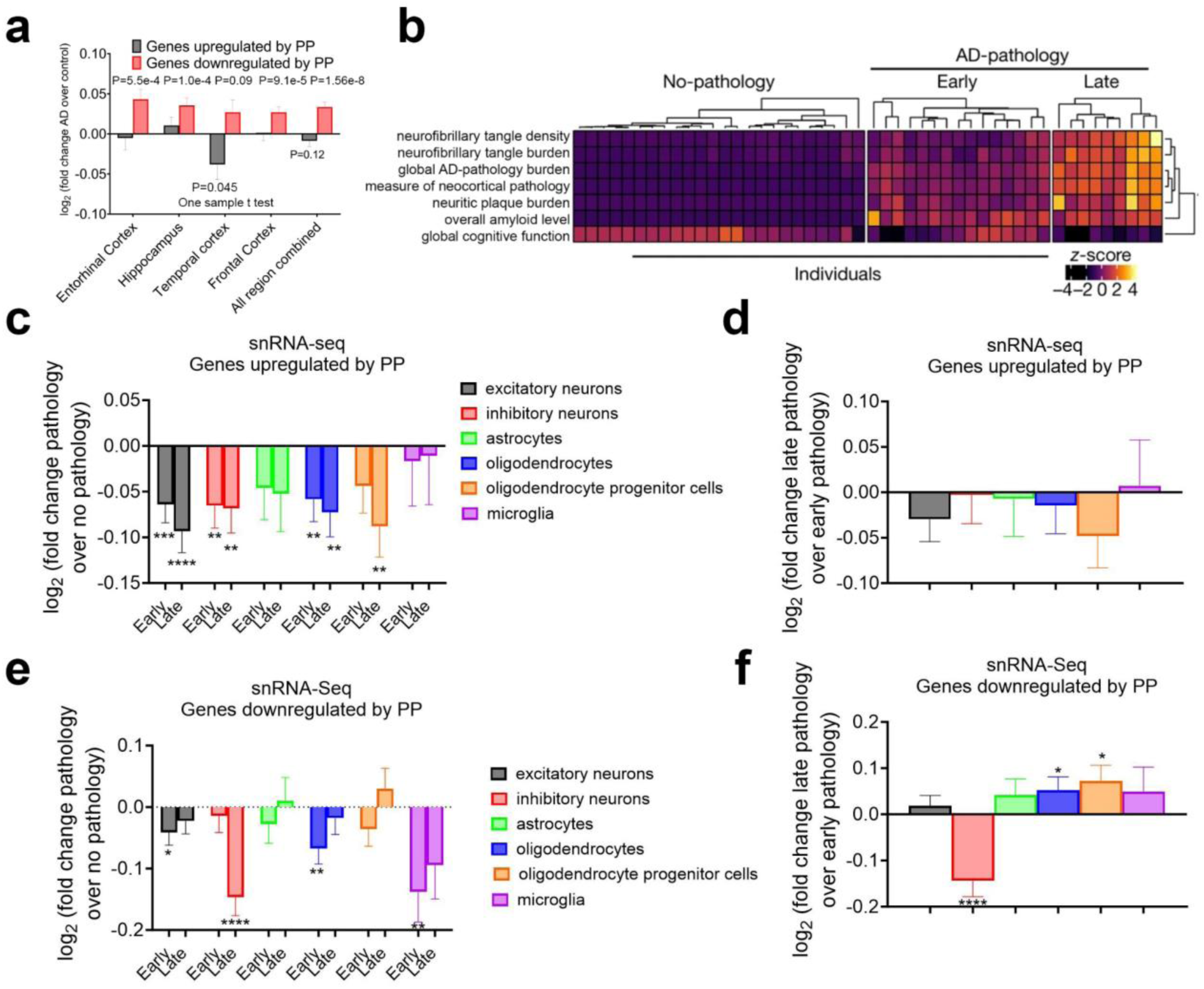
Gene signatures in late-stage human AD subjects with severe tauopathy are opposite from those regulated by PP. (**a**) Log _2_ transformed values of fold change of gene expression of different individual or combined brain regions of AD versus control human subjects for top genes that were either upregulated or downregulated by PP (with a log_2_ fold change > 1.5) in MEFs. (**b**) Phenotypic clustering of 48 individuals (columns) using seven clinicopathological variables as reported and adapted from ^87^. (**c-f**) Log _2_ transformed values of fold change of mean gene expression of different cell types of early or late AD versus no pathology human subjects for top genes that were either upregulated (**c**) or downregulated by PP in MEFs (**e**). Log 2 transformed values of fold change of mean gene expression of different cell types of late AD versus early AD human subjects for top genes that were either upregulated (**d**) or downregulated by PP in MEFs (**f**). Data: Mean ± S.E.M. Statistical tests used: Two-tailed one sample t-test (different from 0) for all. * p<0.05, ** p<0.01, *** p<0.001, **** p<0.0001 for c-f.

**Supplementary Figure 25.**
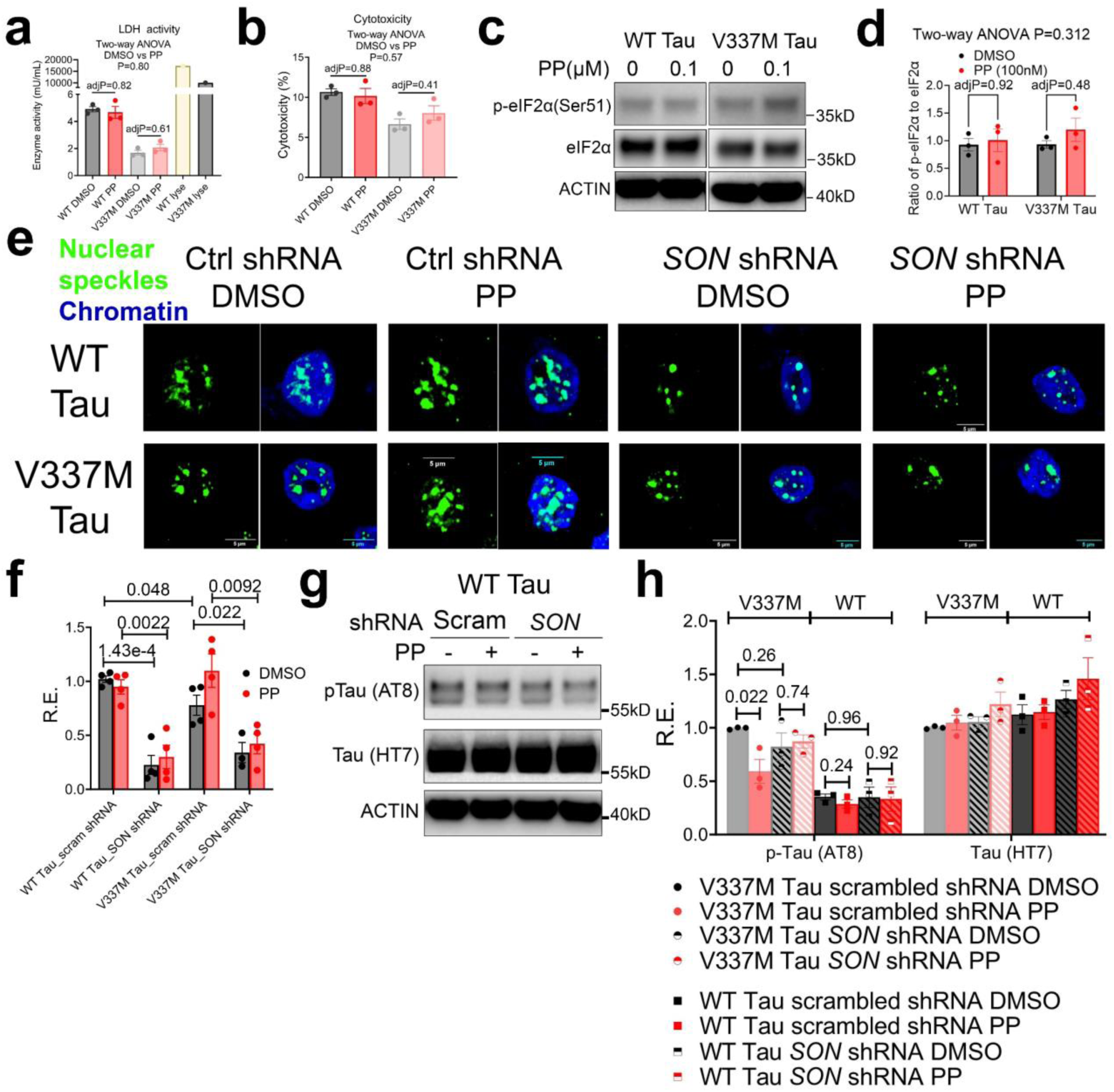
PP reduced V337M Tau level in iPSC neurons. (**a, b**) WT and V337M Tau iPSC neurons were treated with DMSO or 100nM PP for 12 hours. LDH activities in the media were measured. Media from triton-x lysed neurons were included for the maximum amount of LDH release. Both raw activity data (**a**) and triton-x normalized cytotoxicity (**b**) were shown (n=3 biologically independent samples for each treatment). (**c, d**) Representative Western blot images (**c**) and quantification (**d**) showing levels of different proteins in lysates of WT and V337M Tau iPSC neurons treated with DMSO or 100nM PP for 12 hours (n=3 biologically independent samples for each treatment). (**e-h**) WT and V337M Tau iPSC-neurons were infected with scrambled or SON shRNA lentivirus and treated with DMSO or 100nM PP for 12 hours. Representative images of IF against SRRM2 (**e**). qPCR of SON expression (n=3∼4 biologically independent samples for each treatment) (**f**). Representative western blot of WT p-Tau and total Tau (**g**) and quantification (**h**) (n=3 biologically independent samples for each treatment). For h, the V337M Tau data is a replicate of Fig. 7l. Data: Mean ± S.E.M. Statistical tests used: Two-way ANOVA and Šídák’s multiple comparisons test for a, b and d, unpaired two-tailed Student’s t-test for f and h. All biologically independent samples used for Western blot analyses were collected from three independent experiments. The corresponding raw blot images are provided in the source data file.

## Supplementary Movie legends

**Supplementary Movie 1.** Time lapse imaging of droplet formation with 20µm SON IDR1 in 125mM NaCl.

**Supplementary Movie 2.** Time lapse imaging of droplet formation with 20µm SON IDR2 in 125mM NaCl.

**Supplementary Movie 3.** Bright field time lapse imaging of SON IDR2 condensate stretching with bead detachment as the outcome with no PP, example 1.

**Supplementary Movie 4.** Bright field time lapse imaging of SON IDR2 condensate stretching with bead detachment as the outcome with no PP, example 2.

**Supplementary Movie 5.** Bright field time lapse imaging of SON IDR2 condensate stretching with no rupture with no PP.

**Supplementary Movie 6.** Bright field time lapse imaging of SON IDR2 condensate stretching with rupture with 25nM PP.

**Supplementary Movie 7.** Fluorescent time lapse imaging of SON IDR2 condensate stretching with rupture with 25nM PP.

**Supplementary Movie 8.** Bright field time lapse imaging of SON IDR2 condensate stretching with rupture with 100nM PP.

**Supplementary Movie 9.** Fluorescent time lapse imaging of SON IDR2 condensate stretching with rupture with 100nM PP.

**Supplementary Movie 10.** Bright field time lapse imaging of SON IDR2 condensate stretching with rupture with 500nM PP, example 1.

**Supplementary Movie 11.** Fluorescent time lapse imaging of SON IDR2 condensate stretching with rupture with 500nM PP, example 1.

**Supplementary Movie 12.** Bright field time lapse imaging of SON IDR2 condensate stretching with rupture with 500nM PP, example 2.

**Supplementary Movie 13.** Fluorescent time lapse imaging of SON IDR2 condensate stretching with rupture with 500nM PP, example 2.

**Supplementary Movie 14.** Time lapse imaging of droplet formation with 10µm SON IDR2 supplemented with 0.6mg/ml GFP::SRSF2 MEF NE.

## Supplementary Table legends

**Table S1.** FPKM normalization of RNA-seq of SON OE and KD cells in the absence or presence of Tunicamycin in MEFs.

**Table S2.** TPM normalization of RNA-seq of different chemical treatments in MEFs.

## Notes

### Summary of Updates

We have revised all stats and also included optical tweezer experiments showing that PP reduces the surface tension of nuclear speckle condensates in a cell-free system.

## References

1. Gestwicki JE, Garza D. Protein quality control in neurodegenerative disease. Prog Mol Biol Transl Sci 107, 327–353 (2012).

2. Golde TE, Borchelt DR, Giasson BI, Lewis J. Thinking laterally about neurodegenerative proteinopathies. J Clin Invest 123, 1847–1855 (2013).

3. Zhu B, et al. A Cell-Autonomous Mammalian 12 hr Clock Coordinates Metabolic and Stress Rhythms. Cell Metab 25, 1305–1319 e1309 (2017).

4. Zhu B, Dacso CC, O’Malley BW. Unveiling “Musica Universalis” of the Cell: A Brief History of Biological 12-Hour Rhythms. J Endocr Soc 2, 727–752 (2018).

5. Meng H, et al. XBP1 links the 12-hour clock to NAFLD and regulation of membrane fluidity and lipid homeostasis. Nature Communications 11, 6215 (2020).

6. Pan Y, et al. 12-h clock regulation of genetic information flow by XBP1s. PLOS Biology 18, e3000580 (2020).

7. Ballance H, Zhu B. Revealing the hidden reality of the mammalian 12-h ultradian rhythms. Cellular and Molecular Life Sciences, (2021).

8. Asher G, Zhu B. Beyond circadian rhythms: Emerging roles of ultradian rhythms in control of liver functions. Hepatology **n/a**, (2022).

9. Dion W, et al. Four-dimensional nuclear speckle phase separation dynamics regulate proteostasis. Science Advances 8, eabl4150 (2022).

10. Meng H, et al. Defining the mammalian coactivation of hepatic 12-h clock and lipid metabolism. Cell Reports 38, 110491 (2022).

11. Scott MR, et al. Twelve-hour rhythms in transcript expression within the human dorsolateral prefrontal cortex are altered in schizophrenia. PLOS Biology 21, e3001688 (2023).

12. Zhu B, et al. Evidence for conservation of a primordial 12-hour ultradian gene program in humans. bioRxiv, 2023.2005.2002.539021 (2023).

13. Zhu B, Liu S. Preservation of ∼12-h ultradian rhythms of gene expression of mRNA and protein metabolism in the absence of canonical circadian clock. Frontiers in Physiology 14, (2023).

14. Zhu B, et al. Evidence for ∼12-h ultradian gene programs in humans. npj Biological Timing and Sleep 1, 4 (2024).

15. Spector DL, Lamond AI. Nuclear speckles. Cold Spring Harb Perspect Biol 3, (2011).

16. Hasenson SE, Shav-Tal Y. Speculating on the Roles of Nuclear Speckles: How RNA-Protein Nuclear Assemblies Affect Gene Expression. Bioessays 42, e2000104 (2020).

17. Liao SE, Regev O. Splicing at the phase-separated nuclear speckle interface: a model. Nucleic Acids Research 49, 636–645 (2020).

18. Alexander KA, et al. Nuclear speckles regulate HIF-2α programs and correlate with patient survival in kidney cancer. *bioRxiv*, 2023.2009.2014.557228 (2023).

19. Alexander KA, et al. p53 mediates target gene association with nuclear speckles for amplified RNA expression. Mol Cell 81, 1666–1681 e1666 (2021).

20. Dion W, et al. Four-dimensional nuclear speckle phase separation dynamics regulate proteostasis. Science Advances 8, eabl4150 (2022).

21. Jia M, et al. Transcriptional changes of the aging lung. Aging Cell 22, e13969 (2023).

22. Chavez A, et al. Comparison of Cas9 activators in multiple species. Nature methods 13, 563–567 (2016).

23. Qin Q, et al. Lisa: inferring transcriptional regulators through integrative modeling of public chromatin accessibility and ChIP-seq data. Genome Biology 21, 32 (2020).

24. Dopie J, Sweredoski MJ, Moradian A, Belmont AS. Tyramide signal amplification mass spectrometry (TSA-MS) ratio identifies nuclear speckle proteins. Journal of Cell Biology 219, (2020).

25. Saitoh N, Spahr CS, Patterson SD, Bubulya P, Neuwald AF, Spector DL. Proteomic analysis of interchromatin granule clusters. Mol Biol Cell 15, 3876–3890 (2004).

26. Cretenet G, Le Clech M, Gachon F. Circadian clock-coordinated 12 Hr period rhythmic activation of the IRE1alpha pathway controls lipid metabolism in mouse liver. Cell Metab 11, 47–57 (2010).

27. Pocaterra A, Romani P, Dupont S. YAP/TAZ functions and their regulation at a glance. Journal of Cell Science 133, jcs230425 (2020).

28. Panciera T, Azzolin L, Cordenonsi M, Piccolo S. Mechanobiology of YAP and TAZ in physiology and disease. Nature Reviews Molecular Cell Biology 18, 758–770 (2017).

29. Piccolo S, Dupont S, Cordenonsi M. The Biology of YAP/TAZ: Hippo Signaling and Beyond. Physiological Reviews 94, 1287–1312 (2014).

30. Reich S, et al. A multi-omics analysis reveals the unfolded protein response regulon and stress-induced resistance to folate-based antimetabolites. Nature Communications 11, 2936 (2020).

31. Lu X, Ng H-H, Bubulya PA. The role of SON in splicing, development, and disease. Wiley Interdiscip Rev RNA 5, 637–646 (2014).

32. Ahn EY, et al. SON controls cell-cycle progression by coordinated regulation of RNA splicing. Mol Cell 42, 185–198 (2011).

33. Kim J-H, et al. SON drives oncogenic RNA splicing in glioblastoma by regulating PTBP1/PTBP2 switching and RBFOX2 activity. Nature Communications 12, 5551 (2021).

34. Li H-D, Funk CC, Price ND. iREAD: a tool for intron retention detection from RNA-seq data. BMC Genomics 21, 128 (2020).

35. Sureau A, Gattoni R, Dooghe Y, Stévenin J, Soret J. SC35 autoregulates its expression by promoting splicing events that destabilize its mRNAs. Embo j 20, 1785–1796 (2001).

36. Ding F, Su CJ, Edmonds KK, Liang G, Elowitz MB. Dynamics and functional roles of splicing factor autoregulation. Cell Rep 39, 110985 (2022).

37. Grandjean JMD, et al. Pharmacologic IRE1/XBP1s activation confers targeted ER proteostasis reprogramming. Nat Chem Biol 16, 1052–1061 (2020).

38. Madhavan A, et al. Pharmacologic IRE1/XBP1s activation promotes systemic adaptive remodeling in obesity. Nature Communications 13, 608 (2022).

39. Grandjean JMD, Wiseman RL. Small molecule strategies to harness the unfolded protein response: where do we go from here? Journal of Biological Chemistry 295, 15692–15711 (2020).

40. Molina DM, et al. Monitoring Drug Target Engagement in Cells and Tissues Using the Cellular Thermal Shift Assay. Science 341, 84–87 (2013).

41. Jafari R, et al. The cellular thermal shift assay for evaluating drug target interactions in cells. Nature Protocols 9, 2100–2122 (2014).

42. Ilik İA, Malszycki M, Lübke AK, Schade C, Meierhofer D, Aktaş T. SON and SRRM2 are essential for nuclear speckle formation. eLife 9, e60579 (2020).

43. Fei J, et al. Quantitative analysis of multilayer organization of proteins and RNA in nuclear speckles at super resolution. Journal of Cell Science 130, 4180–4192 (2017).

44. Sharma A, Takata H, Shibahara K, Bubulya A, Bubulya PA. Son is essential for nuclear speckle organization and cell cycle progression. Mol Biol Cell 21, 650–663 (2010).

45. Liao SE, Regev O. Splicing at the phase-separated nuclear speckle interface: a model. Nucleic Acids Res 49, 636–645 (2021).

46. Xu S, Lai SK, Sim DY, Ang WSL, Li HY, Roca X. SRRM2 organizes splicing condensates to regulate alternative splicing. Nucleic Acids Res 50, 8599–8614 (2022).

47. Gouveia B, Kim Y, Shaevitz JW, Petry S, Stone HA, Brangwynne CP. Capillary forces generated by biomolecular condensates. Nature 609, 255–264 (2022).

48. Sabari BR, et al. Coactivator condensation at super-enhancers links phase separation and gene control. Science 361, eaar3958 (2018).

49. Chen Z, et al. Screening membraneless organelle participants with machine-learning models that integrate multimodal features. Proceedings of the National Academy of Sciences 119, e2115369119 (2022).

50. Romero, Obradovic, Dunker K. Sequence Data Analysis for Long Disordered Regions Prediction in the Calcineurin Family. Genome Inform Ser Workshop Genome Inform 8, 110–124 (1997).

51. Walsh I, Martin AJM, Di Domenico T, Tosatto SCE. ESpritz: accurate and fast prediction of protein disorder. Bioinformatics 28, 503–509 (2011).

52. Greig JA, et al. Arginine-Enriched Mixed-Charge Domains Provide Cohesion for Nuclear Speckle Condensation. Molecular Cell 77, 1237–1250.e1234 (2020).

53. Santos J, Calero N, Trujillo-Cayado LA, Garcia MC, Muñoz J. Assessing differences between Ostwald ripening and coalescence by rheology, laser diffraction and multiple light scattering. Colloids and Surfaces B: Biointerfaces 159, 405–411 (2017).

54. Ghosh A, Kota D, Zhou H-X. Shear relaxation governs fusion dynamics of biomolecular condensates. Nature Communications 12, 5995 (2021).

55. Ambadi Thody S, Clements HD, Baniasadi H, Lyon AS, Sigman MS, Rosen MK. Small-molecule properties define partitioning into biomolecular condensates. Nature Chemistry 16, 1794–1802 (2024).

56. Riback JA, et al. Viscoelasticity and advective flow of RNA underlies nucleolar form and function. Molecular Cell 83, 3095–3107.e3099 (2023).

57. Pancholi A, et al. RNA polymerase II clusters form in line with liquid phase wetting of chromatin. bioRxiv, 2021.2002.2003.429626 (2021).

58. Liu X, et al. Time-dependent effect of 1,6-hexanediol on biomolecular condensates and 3D chromatin organization. Genome Biology 22, 230 (2021).

59. Liu J, Rivas FV, Wohlschlegel J, Yates JR, 3rd, Parker R, Hannon GJ. A role for the P-body component GW182 in microRNA function. Nat Cell Biol 7, 1261–1266 (2005).

60. Ravi V, Jain A, Mishra S, Sundaresan NR. Measuring Protein Synthesis in Cultured Cells and Mouse Tissues Using the Non-radioactive SUnSET Assay. Curr Protoc Mol Biol 133, e127 (2020).

61. Pakos-Zebrucka K, Koryga I, Mnich K, Ljujic M, Samali A, Gorman AM. The integrated stress response. EMBO Rep 17, 1374–1395 (2016).

62. Mauvezin C, Neufeld TP. Bafilomycin A1 disrupts autophagic flux by inhibiting both V-ATPase-dependent acidification and Ca-P60A/SERCA-dependent autophagosome-lysosome fusion. Autophagy 11, 1437–1438 (2015).

63. Harada Y, Ishii I, Hatake K, Kasahara T. Pyrvinium pamoate inhibits proliferation of myeloma/erythroleukemia cells by suppressing mitochondrial respiratory complex I and STAT3. Cancer Letters 319, 83–88 (2012).

64. Ishii I, Harada Y, Kasahara T. Reprofiling a classical anthelmintic, pyrvinium pamoate, as an anti-cancer drug targeting mitochondrial respiration. Frontiers in Oncology 2, (2012).

65. Lee E-J, Chan P, Chea L, Kim K, Kaufman RJ, Lin JH. ATF6 is required for efficient rhodopsin clearance and retinal homeostasis in the P23H rho retinitis pigmentosa mouse model. Scientific Reports 11, 16356 (2021).

66. Chiang W-C, et al. Robust Endoplasmic Reticulum-Associated Degradation of Rhodopsin Precedes Retinal Degeneration. Molecular Neurobiology 52, 679–695 (2015).

67. Chiang W-C, Hiramatsu N, Messah C, Kroeger H, Lin JH. Selective Activation of ATF6 and PERK Endoplasmic Reticulum Stress Signaling Pathways Prevent Mutant Rhodopsin Accumulation. Investigative Ophthalmology & Visual Science 53, 7159–7166 (2012).

68. Lobanova ES, et al. Increased proteasomal activity supports photoreceptor survival in inherited retinal degeneration. Nat Commun 9, 1738 (2018).

69. Liu X, et al. Pharmacological clearance of misfolded rhodopsin for the treatment of RHO-associated retinitis pigmentosa. The FASEB Journal 34, 10146–10167 (2020).

70. Lee MJ, Lee JH, Rubinsztein DC. Tau degradation: The ubiquitin–proteasome system versus the autophagy-lysosome system. Progress in Neurobiology 105, 49–59 (2013).

71. Limanaqi F, Biagioni F, Gambardella S, Familiari P, Frati A, Fornai F. Promiscuous Roles of Autophagy and Proteasome in Neurodegenerative Proteinopathies. Int J Mol Sci 21, (2020).

72. Samelson AJ, et al. CRISPR screens in iPSC-derived neurons reveal principles of tau proteostasis. *bioRxiv*, 2023.2006.2016.545386 (2023).

73. Bravo CP, et al. Human iPSC 4R tauopathy model uncovers modifiers of tau propagation. *bioRxiv*, 2023.2006.2019.544278 (2023).

74. Bugiani O, et al. Frontotemporal dementia and corticobasal degeneration in a family with a P301S mutation in tau. Journal of neuropathology and experimental neurology 58, 667–677 (1999).

75. Chen X, et al. Promoting tau secretion and propagation by hyperactive p300/CBP via autophagy-lysosomal pathway in tauopathy. Molecular Neurodegeneration 15, 2 (2020).

76. Triastuti E, et al. Pharmacological inhibition of Hippo pathway, with the novel kinase inhibitor XMU-MP-1, protects the heart against adverse effects during pressure overload. Br J Pharmacol 176, 3956–3971 (2019).

77. Hao X, et al. XMU-MP-1 attenuates osteoarthritis via inhibiting cartilage degradation and chondrocyte apoptosis. Frontiers in Bioengineering and Biotechnology 10, (2022).

78. Kastan N, et al. Small-molecule inhibition of Lats kinases may promote Yap-dependent proliferation in postmitotic mammalian tissues. Nature Communications 12, 3100 (2021).

79. Qin F, Tian J, Zhou D, Chen L. Mst1 and Mst2 kinases: regulations and diseases. Cell & Bioscience 3, 31 (2013).

80. Arno G, et al. Mutations in REEP6 Cause Autosomal-Recessive Retinitis Pigmentosa. Am J Hum Genet 99, 1305–1315 (2016).

81. Pi S, et al. Fully automated OCT-based tissue screening system. Opt Lett 49, 4481–4484 (2024).

82. Wittmann CW, et al. Tauopathy in Drosophila: Neurodegeneration Without Neurofibrillary Tangles. Science 293, 711–714 (2001).

83. Berger C, Renner S, Lüer K, Technau GM. The commonly used marker ELAV is transiently expressed in neuroblasts and glial cells in the Drosophila embryonic CNS. Developmental Dynamics 236, 3562–3568 (2007).

84. Hirabayashi S, Baranski TJ, Cagan RL. Transformed Drosophila cells evade diet-mediated insulin resistance through wingless signaling. Cell 154, 664–675 (2013).

85. Wang L, et al. JNK modifies neuronal metabolism to promote proteostasis and longevity. Aging Cell 18, e12849 (2019).

86. Xu M, et al. A systematic integrated analysis of brain expression profiles reveals YAP1 and other prioritized hub genes as important upstream regulators in Alzheimer’s disease. Alzheimers Dement 14, 215–229 (2018).

87. Mathys H, et al. Single-cell transcriptomic analysis of Alzheimer’s disease. Nature 570, 332–337 (2019).

88. Wang C, et al. Scalable Production of iPSC-Derived Human Neurons to Identify Tau-Lowering Compounds by High-Content Screening. Stem Cell Reports 9, 1221–1233 (2017).

89. Wu R, et al. Disruption of nuclear speckle integrity dysregulates RNA splicing in C9ORF72-FTD/ALS. Neuron, (2024).

90. Lester E, et al. Tau aggregates are RNA-protein assemblies that mislocalize multiple nuclear speckle components. Neuron 109, 1675–1691.e1679 (2021).

91. McMillan PJ, et al. Pathological tau drives ectopic nuclear speckle scaffold protein SRRM2 accumulation in neuron cytoplasm in Alzheimer’s disease. Acta Neuropathologica Communications 9, 117 (2021).

92. Mordes D, et al. Pre-mRNA splicing and retinitis pigmentosa. Mol Vis 12, 1259–1271 (2006).

93. Lester E, et al. Tau aggregates are RNA-protein assemblies that mislocalize multiple nuclear speckle components. Neuron 109, 1675–1691 e1679 (2021).

94. Hsieh YC, et al. Tau-Mediated Disruption of the Spliceosome Triggers Cryptic RNA Splicing and Neurodegeneration in Alzheimer’s Disease. Cell Rep 29, 301–316 e310 (2019).

95. Wheeler RJ. Therapeutics-how to treat phase separation-associated diseases. Emerg Top Life Sci 4, 307–318 (2020).

96. Conti BA, Oppikofer M. Biomolecular condensates: new opportunities for drug discovery and RNA therapeutics. Trends in Pharmacological Sciences 43, 820–837 (2022).

97. Abramson J, et al. Accurate structure prediction of biomolecular interactions with AlphaFold 3. Nature 630, 493–500 (2024).

98. Cai D, Sukenik S, Feliciano D, Gruebele M, Lippincott-Schwartz J. Phase Separation of YAP Reprograms Cells for Long-term YAP Target Gene Expression. bioRxiv, 438416 (2018).

99. Yang S, et al. Bip-Yorkie interaction determines oncogenic and tumor-suppressive roles of Ire1/Xbp1s activation. Proceedings of the National Academy of Sciences 119, e2202133119 (2022).

100. Rivera-Reyes A, et al. YAP1 enhances NF-κB-dependent and independent effects on clock-mediated unfolded protein responses and autophagy in sarcoma. Cell Death Dis 9, 1108 (2018).

101. Stashi E, et al. SRC-2 is an essential coactivator for orchestrating metabolism and circadian rhythm. Cell reports 6, 633–645 (2014).

102. Román-Fernández Á, et al. The phospholipid PI(3,4)P(2) is an apical identity determinant. Nat Commun 9, 5041 (2018).

103. Vallazza-Deschamps G, et al. Excessive activation of cyclic nucleotide-gated channels contributes to neuronal degeneration of photoreceptors. Eur J Neurosci 22, 1013–1022 (2005).

104. Vats A, et al. Nonretinoid chaperones improve rhodopsin homeostasis in a mouse model of retinitis pigmentosa. JCI Insight 7, (2022).

105. Pi S, et al. Retinal capillary oximetry with visible light optical coherence tomography. Proceedings of the National Academy of Sciences of the United States of America 117, 11658–11666 (2020).

106. Fanning S, Selkoe D, Dettmer U. Parkinson’s disease: proteinopathy or lipidopathy? NPJ Parkinsons Dis 6, 3 (2020).

107. Wulansari N, et al. Neurodevelopmental defects and neurodegenerative phenotypes in human brain organoids carrying Parkinson’s disease-linked DNAJC6 mutations. Sci Adv 7, (2021).

108. Dion W, et al. Four-dimensional nuclear speckle phase separation dynamics regulate proteostasis. Sci Adv 8, eabl4150 (2022).

109. Dupont S, et al. Role of YAP/TAZ in mechanotransduction. Nature 474, 179–183 (2011).

110. Stirling DR, Swain-Bowden MJ, Lucas AM, Carpenter AE, Cimini BA, Goodman A. CellProfiler 4: improvements in speed, utility and usability. BMC bioinformatics 22, 1–11 (2021).

111. Liu X, et al. Pharmacological clearance of misfolded rhodopsin for the treatment of RHO-associated retinitis pigmentosa. Faseb j 34, 10146–10167 (2020).

112. Schneider CA, Rasband WS, Eliceiri KW. NIH Image to ImageJ: 25 years of image analysis. Nature methods 9, 671–675 (2012).

113. Zhu B, et al. Coactivator-dependent oscillation of chromatin accessibility dictates circadian gene amplitude via REV-ERB loading. Molecular cell 60, 769–783 (2015).

114. Zhu B, et al. Coactivator-Dependent Oscillation of Chromatin Accessibility Dictates Circadian Gene Amplitude via REV-ERB Loading. Mol Cell 60, 769–783 (2015).

115. Zhu B, et al. A Cell-Autonomous Mammalian 12 hr Clock Coordinates Metabolic and Stress Rhythms. Cell Metabolism 25, 1305–1319.e1309 (2017).

116. Zhu B, et al. Peroxisome proliferator-activated receptor beta/delta cross talks with E2F and attenuates mitosis in HRAS-expressing cells. Mol Cell Biol 32, 2065–2082 (2012).

117. Bolger AM, Lohse M, Usadel B. Trimmomatic: a flexible trimmer for Illumina sequence data. Bioinformatics 30, 2114–2120 (2014).

118. Kim D, Langmead B, Salzberg SL. HISAT: a fast spliced aligner with low memory requirements. Nat Methods 12, 357–360 (2015).

119. Quinlan AR, Hall IM. BEDTools: a flexible suite of utilities for comparing genomic features. Bioinformatics 26, 841–842 (2010).

120. Blankenberg D, et al. Galaxy: A Web-Based Genome Analysis Tool for Experimentalists. Current Protocols in Molecular Biology 89, 19.10.11–19.10.21 (2010).

121. Krueger F. Trim Galore.) (2021).

122. Patro R, Duggal G, Love MI, Irizarry RA, Kingsford C. Salmon provides fast and bias-aware quantification of transcript expression. Nature methods 14, 417–419 (2017).

123. Guo W, et al. 3D RNA-seq: a powerful and flexible tool for rapid and accurate differential expression and alternative splicing analysis of RNA-seq data for biologists. RNA biology 18, 1574–1587 (2021).

124. Langmead B, Salzberg SL. Fast gapped-read alignment with Bowtie 2. Nat Methods 9, 357–359 (2012).

125. Li B, Dewey CN. RSEM: accurate transcript quantification from RNA-Seq data with or without a reference genome. BMC Bioinformatics 12, 323 (2011).

126. Robinson MD, McCarthy DJ, Smyth GK. edgeR: a Bioconductor package for differential expression analysis of digital gene expression data. Bioinformatics 26, 139–140 (2010).

127. Huang da W, Sherman BT, Lempicki RA. Systematic and integrative analysis of large gene lists using DAVID bioinformatics resources. Nat Protoc 4, 44–57 (2009).

128. He HH, et al. Nucleosome dynamics define transcriptional enhancers. Nature Genetics 42, 343–347 (2010).

129. Zhang MJ, Pisco AO, Darmanis S, Zou J. Mouse aging cell atlas analysis reveals global and cell type-specific aging signatures. Elife 10, (2021).

130. A single-cell transcriptomic atlas characterizes ageing tissues in the mouse. Nature 583, 590–595 (2020).

131. Jiang J, Wang C, Qi R, Fu H, Ma Q. scREAD: A Single-Cell RNA-Seq Database for Alzheimer’s Disease. iScience 23, 101769 (2020).

